# X-SPEAR: An Extensible Framework for Spatial Panel Evaluation and Refinement

**DOI:** 10.64898/2026.10.13.756865

**Authors:** Rushil Patel, Bryant Duong, Veer Patel, Rachel Lyn-Sue, Kyungbin Oh, Yuyan Cheng

## Abstract

Targeted spatial transcriptomics platforms such as 10x Genomics Xenium measure a predefined panel of genes. A gene is usually detected in fewer cells in the spatial assay than in the single-cell RNA sequencing (scRNA-seq) reference used to choose it, by an amount that differs from gene to gene. We present eXtensible Spatial Panel Evaluation And Refinement (X- SPEAR), a framework for designing and checking Xenium gene panels. Its marker-selection method, Decision-rule Aggregation for Rare-cell Typing (DART), learns short Boolean gene rules and keeps several gene-disjoint rules per cell type. DART first thins the reference to per-gene detection rates measured in a previous experiment, which we call a Xenium prior, so no Xenium data from the target sample are needed before the panel is ordered. A Xenium prior helped most when it matched the target’s assay chemistry and species, even when it came from a different tissue, and a mismatched prior could lower accuracy. With such a prior, DART outperformed four marker-selection methods on held-out Xenium data from ovarian cancer, mouse brain and cervical cancer at matched panel size: by 0.04–0.10 macro-averaged *F*_1_ score (macro-*F*_1_) over the Wilcoxon rank-sum test, 0.04–0.08 over COSine similarity-based marker Gene identification (COSG), 0.10–0.17 over Necessary and Sufficient Forest (NS-Forest) and 0.04–0.08 over the probe-set selection method Spapros. It reached the Wilcoxon test’s best accuracy with 42–69% fewer genes. DART’s rules agree with the classifier that uses the panel and name recognisable marker programs for each cell type. X-SPEAR also checks the probes of any panel for predicted off-target transcripts that are expressed in other cell types. X-SPEAR is available as an open-source Python package.

## 1 Introduction

Targeted in situ transcriptomics platforms such as 10x Genomics Xenium measure hundreds to thousands of genes at subcellular resolution in intact tissue, so that cell types, subtypes and states can be identified in their spatial context (Janesick et al., 2023). They require the user to choose the gene panel before the experiment. The panel is usually chosen from an annotated single-cell RNA sequencing (scRNA-seq) or single-nucleus RNA sequencing (snRNA-seq) reference, and the choice determines which cell identities the experiment can resolve. A probe that hybridises to unintended transcripts can also produce misleading spatial signal, so the designed probes need checking.

Several computational methods select markers or probe sets for this purpose: differential-expression rankings such as the Wilcoxon rank-sum test (Wilcoxon, 1945); specificity scores such as COSine similarity-based marker Gene identification (COSG) (Dai et al., 2022); minimal marker combinations such as Necessary and Sufficient Forest (NS-Forest) (Aevermann et al., 2021); and probe-set optimisation such as Spapros (Kuemmerle et al., 2024). Two problems remain. First, single-gene markers are often insufficient for rare populations, closely related subtypes and transient states, and methods that find gene combinations usually reduce each cell type to one combination, which fails if any of its genes is poorly detected. Second, a gene’s detection in a targeted spatial assay differs from its detection in the single-cell reference. Across the ovarian cancer panel studied here, the median gene is detected in Xenium in only about half as many cells as in the matched reference, and the loss varies strongly from gene to gene. A marker that looks excellent in the reference may be nearly invisible in situ. The methods we compare choose genes from the reference as measured and do not model this loss.

Here we present eXtensible Spatial Panel Evaluation And Refinement (X-SPEAR), a framework with two modules. The first is Decision-rule Aggregation for Rare-cell Typing (DART), which learns short Boolean gene rules and selects several gene-disjoint rules for each cell type. Before learning, DART thins the reference to each gene’s measured single-cell-to-Xenium detection rate, so that rules relying on poorly detected genes score poorly and are not chosen. The detection rates come from a *Xenium prior*: a previous experiment in which a single-cell reference and a Xenium run were both measured, usually on a different tissue, because a designer never has Xenium data from the target sample before ordering its panel. The second module checks the probes of any panel, whichever method chose it, for predicted off-target transcripts that are expressed in other cell types. The current version targets cell-type marker selection for Xenium. The “X” denotes extensibility, toward other targeted platforms and other selection objectives.

We first ask which Xenium priors help DART, using every combination of prior and target among five paired single-cell and Xenium datasets. We then benchmark DART against the Wilcoxon test, COSG, NS-Forest and Spapros on three targets (ovarian cancer, mouse brain and cervical cancer), with a deployment rule throughout: nothing measured on the target’s Xenium informs the panel, and its labels are used only for scoring. With a Xenium prior from a different tissue, DART outperformed all four methods at matched panel size on every target. Its rules agree with the downstream classifier and name interpretable marker programs. On 45 mouse retinal ganglion cell (RGC) subtypes, held out within a single-cell atlas, DART matched the best methods and led COSG and NS-Forest at every subtype abundance.

## 2 Results

### 2.1 X-SPEAR designs a detection-aware panel and checks its probes

X- SPEAR takes an annotated single-cell reference of the tissue to be profiled, together with a Xenium prior, and returns a Xenium gene panel with evidence for classification performance and probe-level off-target risk (Fig. 1). It is a Python package that can be used as a library, from the command line or as a pipeline of twelve stages (below). Panel design with DART proceeds in three steps (Fig. 2).

**Figure 1:**
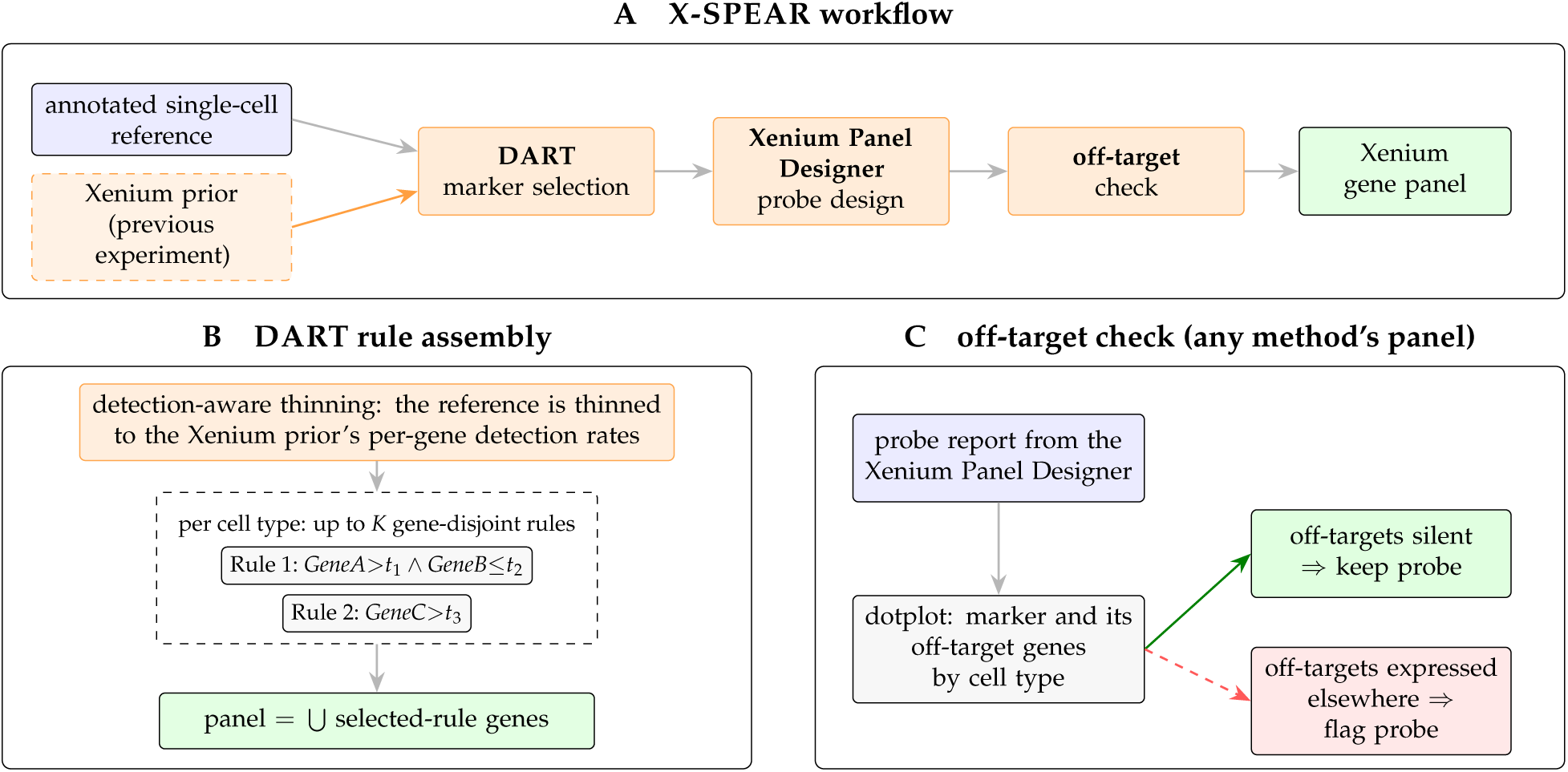
X-SPEAR designs a detection-aware Xenium panel and checks its probes. **(A)** Workflow: DART marker selection from an annotated single-cell reference and a Xenium prior, probe design in the Xenium Panel Designer, and the off-target check. **(B)** DART keeps up to *K* gene-disjoint Boolean rules per cell type, learned on a reference thinned to the Xenium prior’s per-gene detection rates. A rule can require a gene to be high (*>*) or low (*≤*). **(C)** The off-target check reads the probe report of any panel, draws each marker beside its predicted off-target genes across the reference’s cell types, and flags probes whose off- target genes are expressed in cell types the marker does not mark.

**Figure 2:**
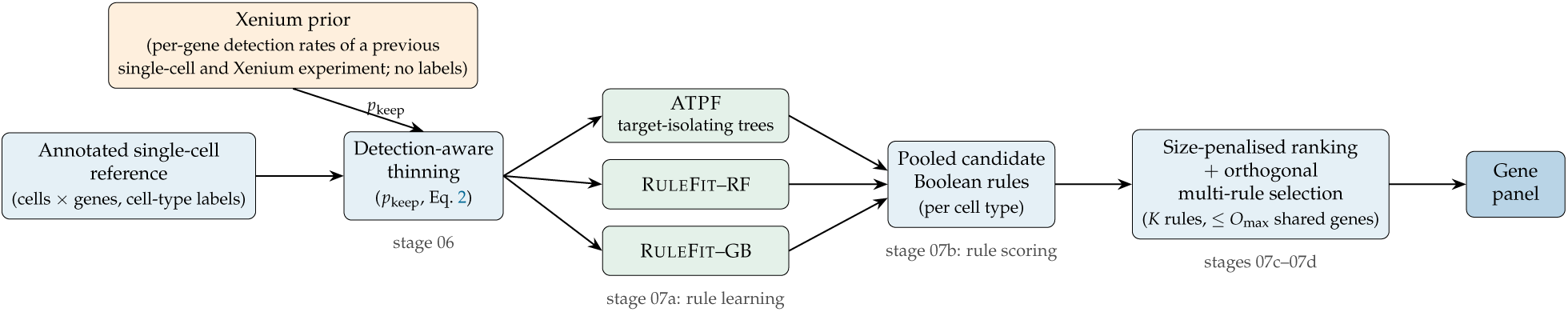
The DART algorithm. The single-cell reference is first thinned so that each gene is detected as often as in the Xenium prior: each nonzero count is kept with probability *p*_keep_ (Eq. 2). The prior is a previous experiment with both a single-cell reference and a Xenium run, usually of another tissue, and its labels are not used. Three learners then propose candidate Boolean rules per cell type: ATPF, whose trees isolate one cell type, and R_ULE_F_IT_ on random-forest (RF) and gradient-boosted (GB) ensembles. Candidates are ranked by *F*_1_ with a penalty per additional gene, and up to *K* rules sharing at most *O*_max_ genes are kept per type. The panel is the union of the selected rules’ genes. Grey labels give the pipeline stage that runs each step (Fig. 3).

The first step is detection-aware thinning. For each gene, a keep probability *p*_keep_ is the ratio of its detection rate (the fraction of cells with a nonzero count) in the Xenium prior’s Xenium run to its detection rate in that experiment’s single-cell reference (Methods). Each nonzero count of the single-cell reference is kept with probability *p*_keep_ and zeroed otherwise, so that the thinned reference reproduces each gene’s expected in situ detection. The prior’s cells need no labels, and the target’s Xenium is never used.

The second step is rule learning. Three learners propose candidate Boolean rules for each cell type: thresholded conjunctions such as *GeneA > t*_1_ *∧ GeneB ≤ t*_2_, which require one gene to be high and another to be low. The Asymmetric Tree Panel Finder (ATPF), developed here, grows decision trees whose splits isolate one target cell type rather than separating all classes. RULEFIT (Friedman and Popescu, 2008), a rule-ensemble method, is applied to random-forest and gradient-boosted ensembles; it extracts rules from the trees and keeps those with nonzero *L*_1_-penalised coefficients.

The third step is orthogonal selection. Candidates are ranked by *F*_1_ minus a penalty per additional gene, and up to *K* rules are kept per cell type, sharing at most *O*_max_ genes. The default is gene-disjoint rules (*O*_max_ = 0). The panel is the union of the selected rules’ genes.

The off-target module reads the probe report that the Xenium Panel Designer returns for a submitted panel, which lists each probe’s predicted off-target genes. For each marker it draws a dotplot of the marker beside its off-target genes across the reference’s cell types. A probe whose offtarget genes are expressed in cell types that the marker does not mark would add false signal there and should be replaced; the dotplot shows which probes these are. Because the check needs only the probe report and the reference, it applies to a panel from any selection method (Section 2.12).

X-SPEAR’s pipeline runs these steps as twelve numbered stages, each configured by its own file and each optional (Fig. 3). Stages 00–04 load and prepare the single-cell reference: ingest, quality control, a cap on the cells per cell type, normalisation and, if wanted, an embedding. Stage 06 thins a copy of the reference to the Xenium prior’s detection rates, and stage 07 selects the panel, with DART (as four substages: rule learning, rule scoring, ranking and orthogonal selection) or with one of the other selectors compared here. Stage 10 exports the panel together with the reference bundle that the Xenium Panel Designer takes, and stage 11 checks the probe report it returns. Where a labelled Xenium run of the target exists, stages 05, 08 and 09 load it, label its cells with a classifier trained on the reference and score the labels, as the evaluations in this study do. Every selector writes the same panel file, which is all the later stages read, so the classification, export and off-target stages serve a panel from any method. The interfaces and each stage are described in the Methods (Section 4.1).

**Figure 3:**
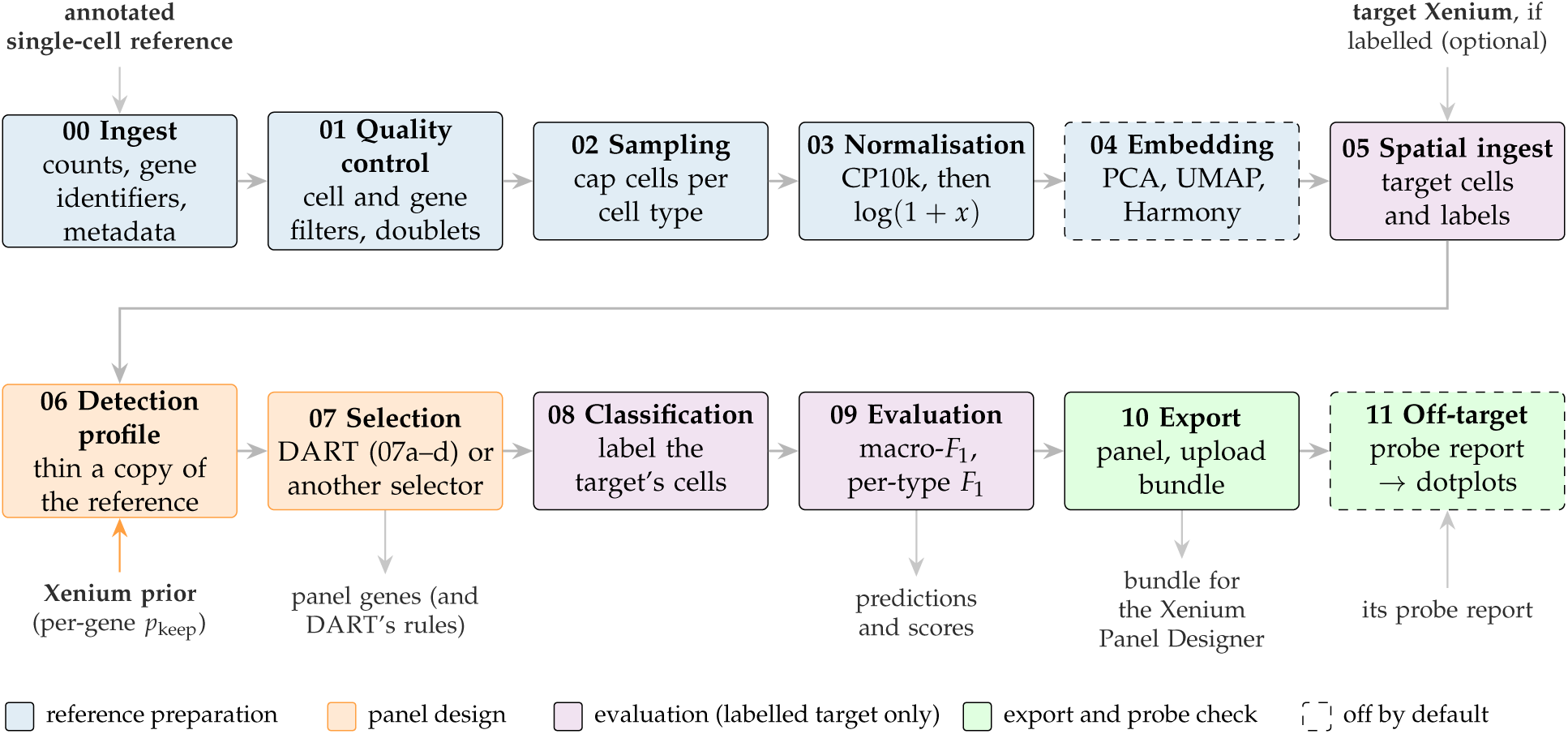
The X-SPEAR pipeline. Twelve numbered stages run in this order, each configured by its own file and each optional. Stages 00–04 prepare the annotated single-cell reference. Stage 06 thins a copy of it to the Xenium prior’s per-gene detection rates, and stage 07 selects the panel: DART as four substages (07a rule learning, 07b rule scoring, 07c ranking, 07d orthogonal selection) or another selector (the Wilcoxon test, COSG, NS-Forest, Spapros or a curated list). Stage 10 exports the panel and the reference bundle for the Xenium Panel Designer, and stage 11 checks the probe report the Designer returns. Stages 05, 08 and 09 evaluate a panel on a labelled Xenium run of the target, when one exists. CP10k, counts per 10,000; PCA, principal component analysis; UMAP, uniform manifold approximation and projection.

### 2.2 Datasets

We used five paired datasets, each with a single-cell reference and a Xenium run, and one single-cell atlas (Table 1). Three pairs serve as *targets*, which need cell-type labels on the Xenium cells to score a panel: ovarian cancer, cervical cancer and mouse brain. All five pairs serve as Xenium priors. Ovarian and cervical cancer are the reference cases: formalin-fixed paraffin-embedded (FFPE) tissue profiled with the 10x Genomics Chromium Flex assay (Flex) and Xenium Prime 5K on the same tissue block, with 10x’s cell-type annotation of the Xenium cells. Mouse brain adds a fresh-frozen (FF) target of another species. The mouse retina pair supplies a same-species prior for mouse brain; it has no cell-type labels, so it cannot be a target. The breast cancer pair is used only as a prior: its Flex reference and its Xenium run come from different tumours, so it is not suitable as a target. The mouse RGC atlas, with 45 annotated subtypes, has no Xenium counterpart with subtype labels, so it is used for a held-out single-cell benchmark of fine subtypes.

**Table 1:**
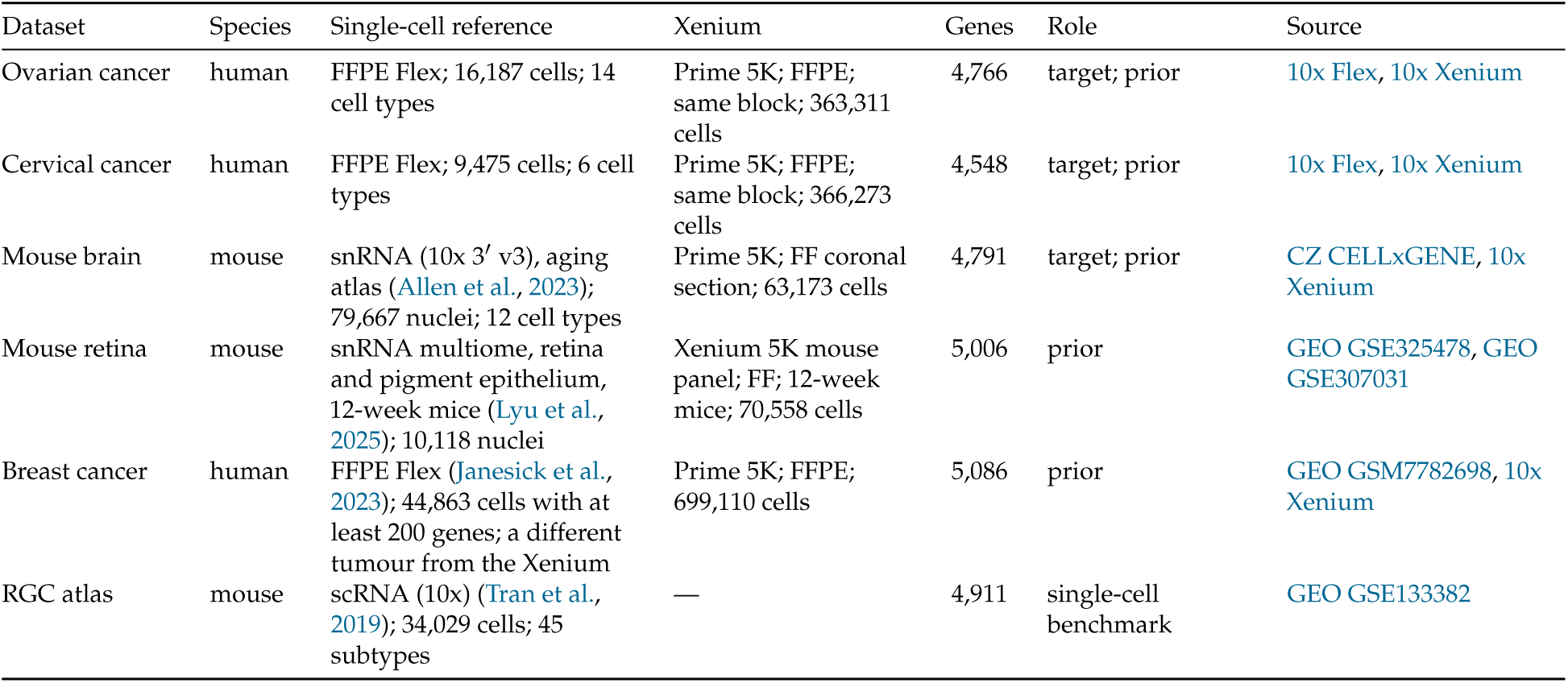
Datasets used in this study. Targets are scored on the held-out half of their Xenium run; every pair also serves as a Xenium prior. Cell counts are those used here (for the targets’ Xenium runs, after quality control). Genes: the genes measured in both the single-cell reference and the Xenium panel (for the RGC atlas, its genes on the mouse Xenium 5K panel). FFPE, formalin-fixed paraffin-embedded; FF, fresh-frozen; Flex, 10x Genomics Chromium Flex; snRNA, single-nucleus RNA sequencing; scRNA, single-cell RNA sequencing. CZ, Chan Zuckerberg; GEO, Gene Expression Omnibus. Full links are in the Data availability statement.

| Dataset | Species | Single-cell reference | Xenium | Genes | Role | Source |
| --- | --- | --- | --- | --- | --- | --- |
| Ovarian cancer | human | FFPE Flex; 16,187 cells; 14 cell types | Prime 5K; FFPE; same block; 363,311 cells | 4,766 | target; prior | 10x Flex, 10x Xenium |
| Cervical cancer | human | FFPE Flex; 9,475 cells; 6 cell types | Prime 5K; FFPE; same block; 366,273 cells | 4,548 | target; prior | 10x Flex, 10x Xenium |
| Mouse brain | mouse | snRNA (10x 3’ v3), aging atlas (Allen et al., 2023); 79,667 nuclei; 12 cell types | Prime 5K; FF coronal section; 63,173 cells | 4,791 | target; prior | CZ CELLxGENE, 10x Xenium |
| Mouse retina | mouse | snRNA multiome, retina and pigment epithelium, 12-week mice (Lyu et al., 2025); 10,118 nuclei | Xenium 5K mouse panel; FF; 12-week mice; 70,558 cells | 5,006 | prior | GEO GSE325478, GEO GSE307031 |
| Breast cancer | human | FFPE Flex (Janesick et al., 2023); 44,863 cells with at least 200 genes; a different tumour from the Xenium | Prime 5K; FFPE; 699,110 cells | 5,086 | prior | GEO GSM7782698, 10x Xenium |
| RGC atlas | mouse | scRNA (10x) (Tran et al., 2019); 34,029 cells; 45 subtypes | — | 4,911 | single-cell benchmark | GEO GSE133382 |

Several other datasets were considered and set aside. A skin melanoma pair was removed because its single-cell reference turned out to be from mouse lemur while its Xenium run was human, so its detection rates measured a species mismatch. A dermatomyositis skin pair had a nearly flat detection profile from a shallow reference and was not validated. A second ovarian cancer Xenium dataset was measured with an earlier 5K panel version lacking about 30% of the genes, and in situ whole-transcriptome Atera data show no detection loss to correct, so neither tests detection-aware design. A human spleen pair was set aside because the source of its Xenium run could not be documented.

### 2.3 Detection loss is large, gene-specific and reproducible across experiments

A Xenium prior is only useful if a gene’s detection loss measured in one experiment predicts its loss in another. On the ovarian cancer Xenium, the median gene is detected in 52% as many cells as in the matched single-cell reference (4,259 genes detected in at least 0.1% of reference cells), and the loss varies widely between genes (Fig. 4a). The loss is reproducible. Per-gene *p*_keep_ in ovarian and cervical cancer, two different tissues profiled on the same Xenium panel, correlate at Spearman *ρ* = 0.88 over 3,056 shared genes. That is more strongly than the genes’ single-cell detection breadth correlates between the two tissues (*ρ* = 0.62), so *p*_keep_ captures a property of the gene and its probes rather than shared biology (Fig. 4b).

**Figure 4:**
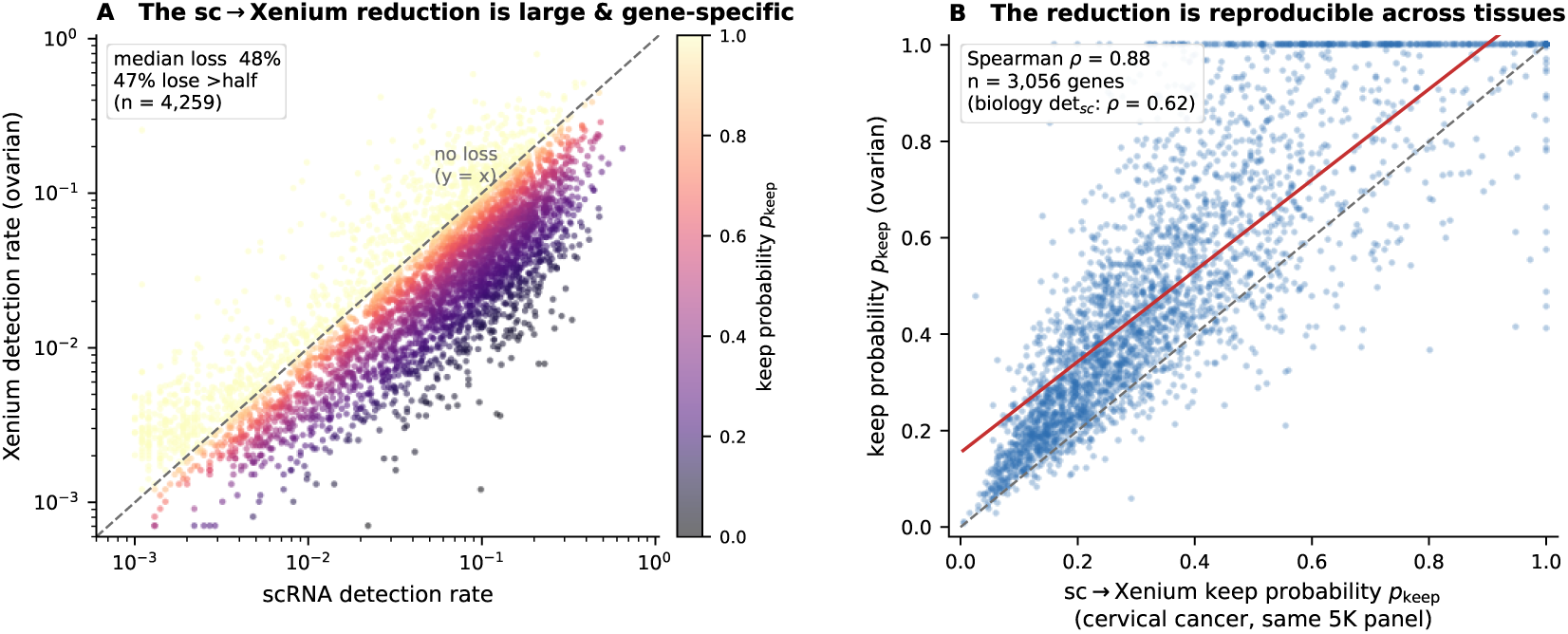
The single-cell-to-Xenium detection loss is large, gene-specific and reproducible across tissues. **(A)** Per-gene detection rate (fraction of cells with a nonzero count) in the ovarian single-cell reference versus ovarian Xenium, for the 4,259 genes detected in at least 0.1% of reference cells. Colour is *p*_keep_. The median gene keeps 52% of its detection. **(B)** Per-gene *p*_keep_ in ovarian versus cervical cancer, two tissues profiled on the same Xenium 5K panel, for the 3,056 genes expressed in at least 1% of reference cells in both (Spearman *ρ* = 0.88; dashed line *y* = *x*, red line least-squares fit). The loss reproduces more faithfully than single-cell detection breadth (*ρ* = 0.62).

### 2.4 Designing a panel without the target’s Xenium

Every panel in this study is designed and evaluated in the same way (Fig. 5). Three datasets are involved. The *single-cell reference* is annotated single-cell data of the target tissue; the panel is chosen from its genes. The *Xenium prior* supplies per-gene detection rates from a previous experiment that measured both a single-cell reference and a Xenium run; only its detection rates are used. The *target Xenium* is the Xenium run of the target tissue. Its Xenium cells were split into two spatial halves by a checkerboard, and only the held-out half was scored. Nothing from the target Xenium informs the panel. To score a panel, a classifier is trained on the single-cell reference restricted to the panel’s genes, it predicts the cell type of every held-out target Xenium cell, and the predictions are compared with the target’s labels by macro-*F*_1_, the unweighted mean of the per-cell-type *F*_1_ scores.

**Figure 5:**
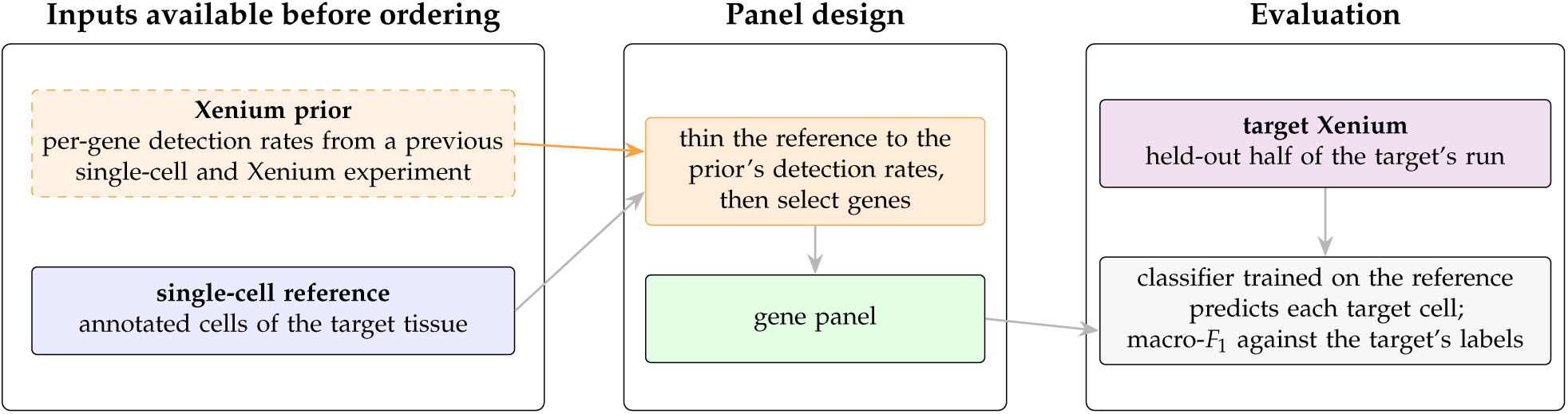
Designing and evaluating a panel without the target’s Xenium. The Xenium prior and the single-cell reference are available before the panel is ordered. The reference is thinned to the prior’s pergene detection rates (DART) or used as measured (the other methods), and a panel is selected. The panel is evaluated on the held-out half of the target Xenium: a classifier trained on the reference, restricted to the panel’s genes, predicts each cell, and the predictions are scored by macro-*F*_1_ against the target’s labels, which are used only here.

### 2.5 A Xenium prior helps when it matches the target’s chemistry and species

We first asked which Xenium priors help DART. For each target we designed DART panels with every available prior, and with none, and compared them at the target’s default panel size (Fig. 6; Methods). The target’s own Xenium is shown as an upper bound that no designer has before ordering.

**Figure 6:**
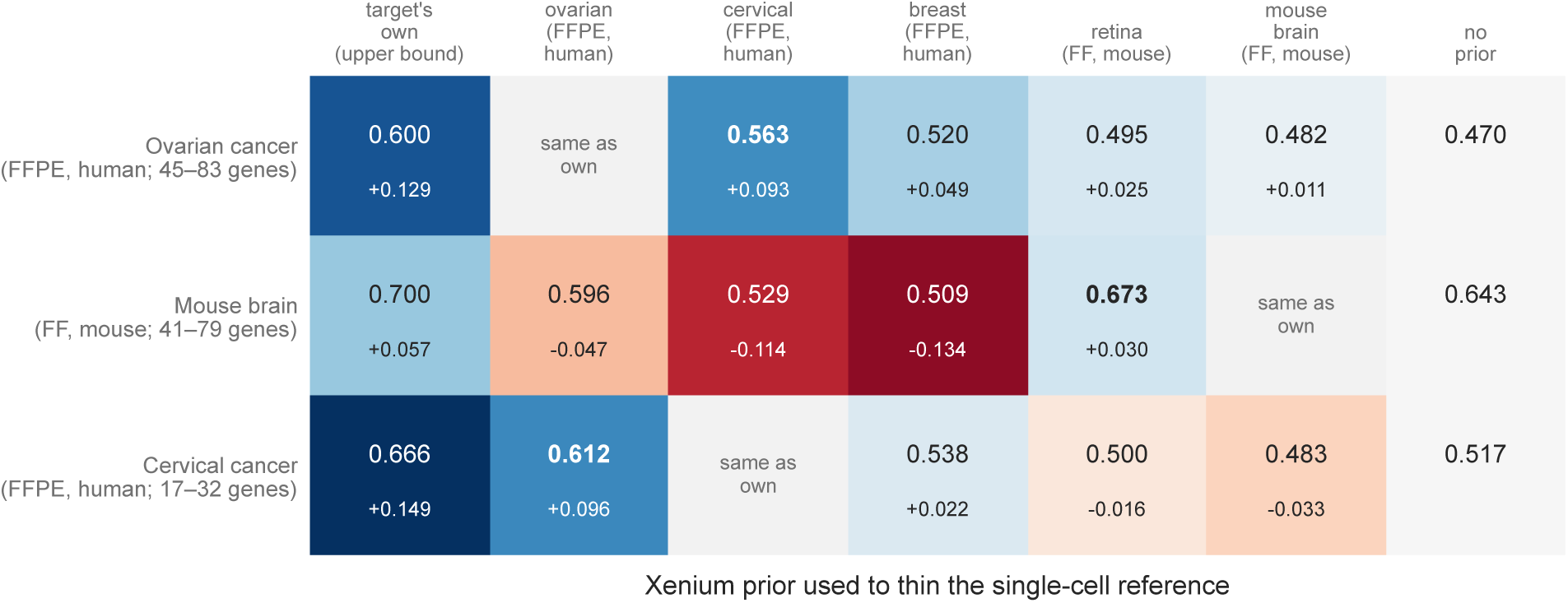
Which Xenium priors help DART. macro-*F*_1_ on the held-out target Xenium of DART panels designed with each Xenium prior (columns) for each target (rows). Each prior’s curve (*K* = 1–10) is interpolated at the sizes of the target’s benchmark panels with *K* = 3–6 (range in the row label) and averaged. The upper number is macro-*F*_1_; the lower number and the colour give the difference from designing with no prior. “Target’s own” uses the target’s own Xenium, which is unavailable before ordering, and is shown as an upper bound. A prior from the other species reaches only the genes whose symbols agree (Methods). FFPE, formalin-fixed paraffin-embedded; FF, fresh-frozen.

The prior that helped most, other than the target’s own, always matched the target’s chemistry and species. For the two human FFPE tumours it was the other tumour, profiled with the same Flex reference and Xenium chemistry: cervical cancer raised ovarian cancer from 0.470 macro-*F*_1_ without a prior to 0.563, against 0.600 with the target’s own Xenium, and ovarian cancer raised cervical cancer from 0.517 to 0.612 (own 0.666). The breast pair, of the same chemistry but with its reference and Xenium run from different tumours, helped less (0.520 and 0.538). For the fresh-frozen mouse brain, the fresh-frozen mouse retina prior helped (0.643 to 0.673; own 0.700), whereas every human FFPE prior lowered accuracy, by 0.05 to 0.13. Conversely, the two fresh-frozen mouse priors helped ovarian cancer little (+0.011 and +0.025) and lowered cervical cancer (*−*0.016 and *−*0.033). A prior from the other species reaches only the genes whose human and mouse symbols agree, about half of the target’s genes (Methods). The best results therefore need a prior of the same chemistry and species as the planned experiment, which can come from a different tissue.

### 2.6 Benchmark against existing marker-selection methods

We then compared DART with four existing methods: the Wilcoxon rank-sum test, COSG, NS-Forest and Spapros. Each target used a Xenium prior from a different tissue, chosen by how closely its samples match the target’s and not by any score on the target: for ovarian and cervical cancer, each other (FFPE tissue with a Flex reference and Xenium Prime 5K on the same block), and for mouse brain, the mouse retina pair, the only other mouse pair, which like the target is fresh-frozen tissue with a single-nucleus reference and the mouse 5K panel (Fig. 7). The other methods select genes from the single-cell reference as measured, as they are normally used. Because panel size drives accuracy, DART was run with *K* = 1–10 rules per cell type and the Wilcoxon test and COSG with 1–12 genes per cell type, and methods were compared at matched realised panel size by interpolating along each accuracy curve. NS-Forest’s panel barely changes with its budget (21– 24 genes on ovarian cancer and one identical panel on each of the other targets), and Spapros was run at its default size, so neither gives an accuracy curve and DART was compared with them at their own panel sizes.

**Figure 7:**
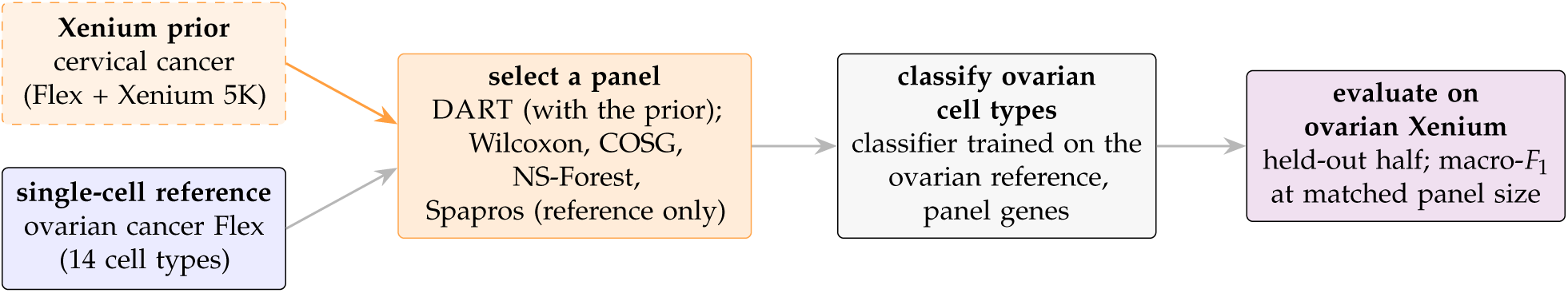
Benchmark design, shown for the ovarian cancer target. The cervical cancer Xenium prior and the ovarian cancer single-cell reference are used to select a panel; a classifier trained on the ovarian reference with the panel’s genes classifies the cells of the held-out half of the ovarian Xenium, and macro-*F*_1_ is compared at matched panel size. The same design is used for the cervical cancer target (ovarian prior) and the mouse brain target (mouse retina prior). Only DART uses the prior; the other methods select from the reference as measured.

### 2.7 DART outperforms existing marker-selection methods

At matched panel size, DART outperformed the Wilcoxon test, COSG, NS-Forest and Spapros on every target (Fig. 8; Table 2). On ovarian cancer, the mean advantage over the Wilcoxon test was +0.064 macro-*F*_1_, ranging from +0.036 to +0.118 across panel sizes, and DART reached the Wilcoxon test’s best accuracy (0.581 with 144 genes) with 83 genes. On mouse brain the advantage was +0.105, and DART needed 41 genes to match the Wilcoxon test’s best, against its 132. On cervical cancer it was +0.044, with 28 genes against 58. Across the three targets the margins were +0.038 to +0.079 over COSG, +0.099 to +0.175 over NS-Forest and +0.044 to +0.082 over Spapros. When DART’s thinning and rule learning were repeated at four seeds, every margin stayed positive (the smallest was +0.039, over Spapros on ovarian cancer), and the seed-to-seed standard deviation of a margin was 0.005–0.017 (Methods).

**Figure 8:**
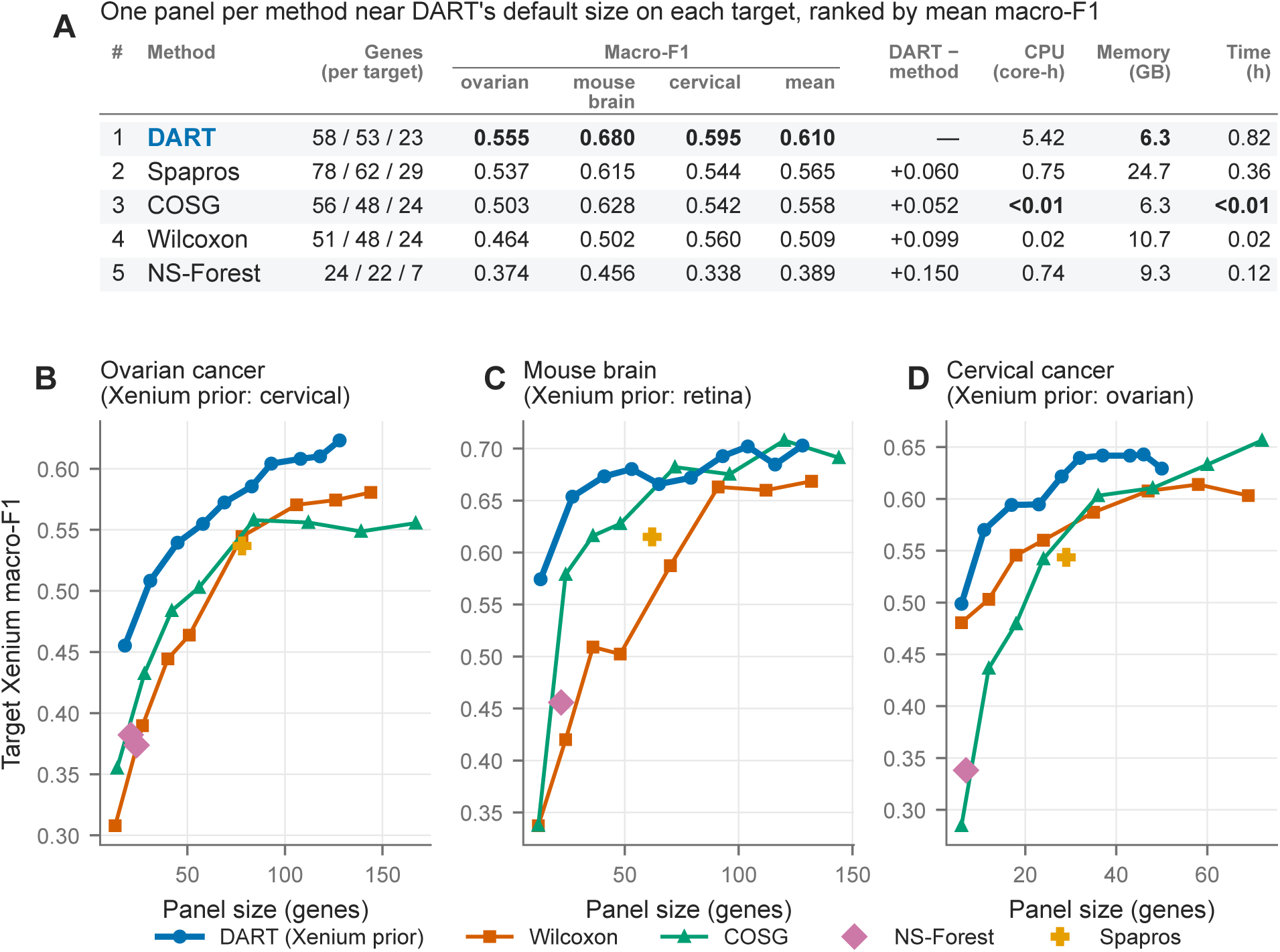
DART outperforms existing marker-selection methods. **(A)** Ranked summary over the three targets. On each target, each method is represented by one panel near DART’s default size (DART with *K* = 4; the Wilcoxon test and COSG at the per-type budget closest in size; NS-Forest at its largest panel; Spapros at its default), and methods are ranked by their mean macro-*F*_1_. Columns give the panel sizes (ovarian / mouse brain / cervical), macro-*F*_1_ on each target and its mean, the difference between DART and the method at the method’s panel size averaged over the targets, and compute measured on this study’s runs: central processing unit (CPU) time and wall time averaged over the targets, and the largest peak memory (Supplementary Table S1). Bold marks the best value in each column. **(B–D)** macro-*F*_1_ on the held-out target Xenium against realised panel size for DART with a Xenium prior (*K* = 1–10) and for the Wilcoxon test and COSG (1–12 genes per cell type), NS-Forest and Spapros. Mouse brain is scored on rebuilt labels (Supplementary S3).

**Table 2:**
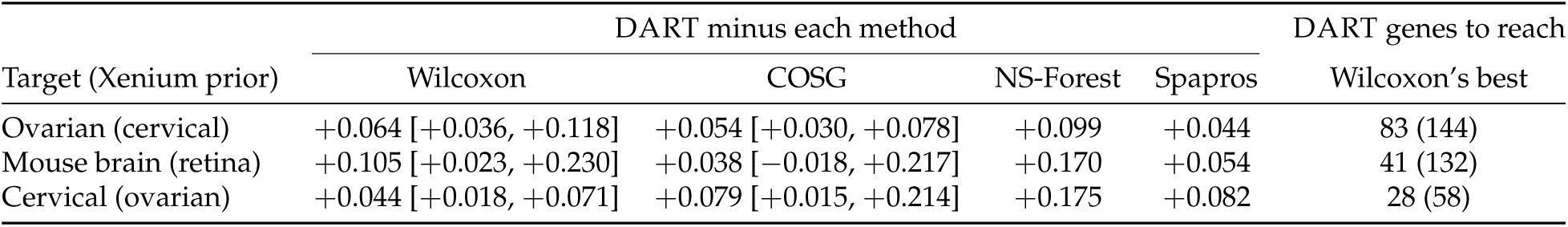
DART (Xenium prior) versus each method at matched panel size. Mean macro-*F*_1_ difference over DART’s realised sizes, with the range across sizes in brackets. NS-Forest (mean over its three budgets) and Spapros are compared at their own panel sizes. The last column gives the number of genes DART needs to match the Wilcoxon test’s best accuracy and, in brackets, the number the Wilcoxon test itself uses. Seed 42; across four seeds the standard deviation of each margin is 0.005–0.017 (Methods). Mouse brain is scored on rebuilt labels (Supplementary S3).

| Target (Xenium prior) | DART minus each method |  |  |  | DART genes to reach |
| --- | --- | --- | --- | --- | --- |
|  | Wilcoxon | COSG | NS-Forest | Spapros | Wilcoxon’s best |
| Ovarian (cervical) | +0.064 [+0.036, +0.118] | +0.054 [+0.030, +0.078] | +0.099 | +0.044 | 83 (144) |
| Mouse brain (retina) | +0.105 [+0.023, +0.230] | +0.038 [−0.018, +0.217] | +0.170 | +0.054 | 41 (132) |
| Cervical (ovarian) | +0.044 [+0.018, +0.071] | +0.079 [+0.015, +0.214] | +0.175 | +0.082 | 28 (58) |

Two qualifications apply to COSG. First, on mouse brain DART’s lead over COSG is concentrated in small panels: it is +0.041 to +0.217 below about 55 genes, but from about 65 genes upward the two are level (*−*0.018 to +0.017), and COSG’s best point (0.708 with 120 genes) is slightly above DART’s best (0.703). On cervical, COSG’s best point (0.656 with 72 genes) lies beyond the largest panel on DART’s curve (50 genes). Second, COSG’s accuracy depends on how the reference is normalised; normalising over the full transcriptome rather than the shared gene universe would raise COSG by about 0.015 on ovarian and 0.036 on cervical (Supplementary S2).

Fig. 8a ranks one panel per method near DART’s default size on each target. DART ranked first on every target and by mean macro-F1 (0.610), ahead of Spapros (0.565), COSG (0.558), the Wilcoxon test (0.509) and NS-Forest (0.389). At each method’s own panel size, DART led Spapros by 0.060, COSG by 0.052, the Wilcoxon test by 0.099 and NS-Forest by 0.150 on average over the three target

The methods also differ in their compute (Fig. 8a; Supplementary Fig. S3). DART splits its rule learning into independent tasks, one per cell type and learner, and runs them on parallel worker processes. It used the most CPU time, 1.2–8.9 core-hours per target against 0.3–1.1 for Spapros and at most 1.8 for the marker selectors, but with eight workers it finished in 12–82 minutes, six to seven times faster than on one core, and its largest process needed at most 6.3 GB of memory. Spapros took 11–28 minutes and needed 16–25 GB, and the Wilcoxon test and COSG finished in under three minutes.

### 2.8 Per-cell-type accuracy on the target Xenium

The differences in macro-*F*_1_ come from particular cell types (Fig. 9). On ovarian cancer, DART’s panel gave the highest *F*_1_ of the five methods for 6 of the 14 cell types and Spapros’s larger panel for 5; DART scored above the Wilcoxon test on 11 types and above NS-Forest on 12. Its largest gain was on T and natural killer (NK) cells (0.50, against 0.14–0.27 for the other methods). It fell furthest behind the best method, by 0.14–0.17, on two tumour states, proliferative and VEGFA-positive tumour cells, which the Wilcoxon test and Spapros separated better. On mouse brain, DART gave the highest *F*_1_ for 6 of 12 cell types and scored above the Wilcoxon test on 11, most of all on neurons (0.86 against 0.24). The two rarest mouse brain types, macrophages (48 held-out cells) and T cells (36), were poorly classified by every method (*F*_1_ at most 0.38). On cervical cancer, with six broad cell types, DART’s advantage came from macrophages (0.64, against 0.09–0.30), and it was weakest on squamous carcinoma (0.28, against 0.29–0.43).

**Figure 9:**
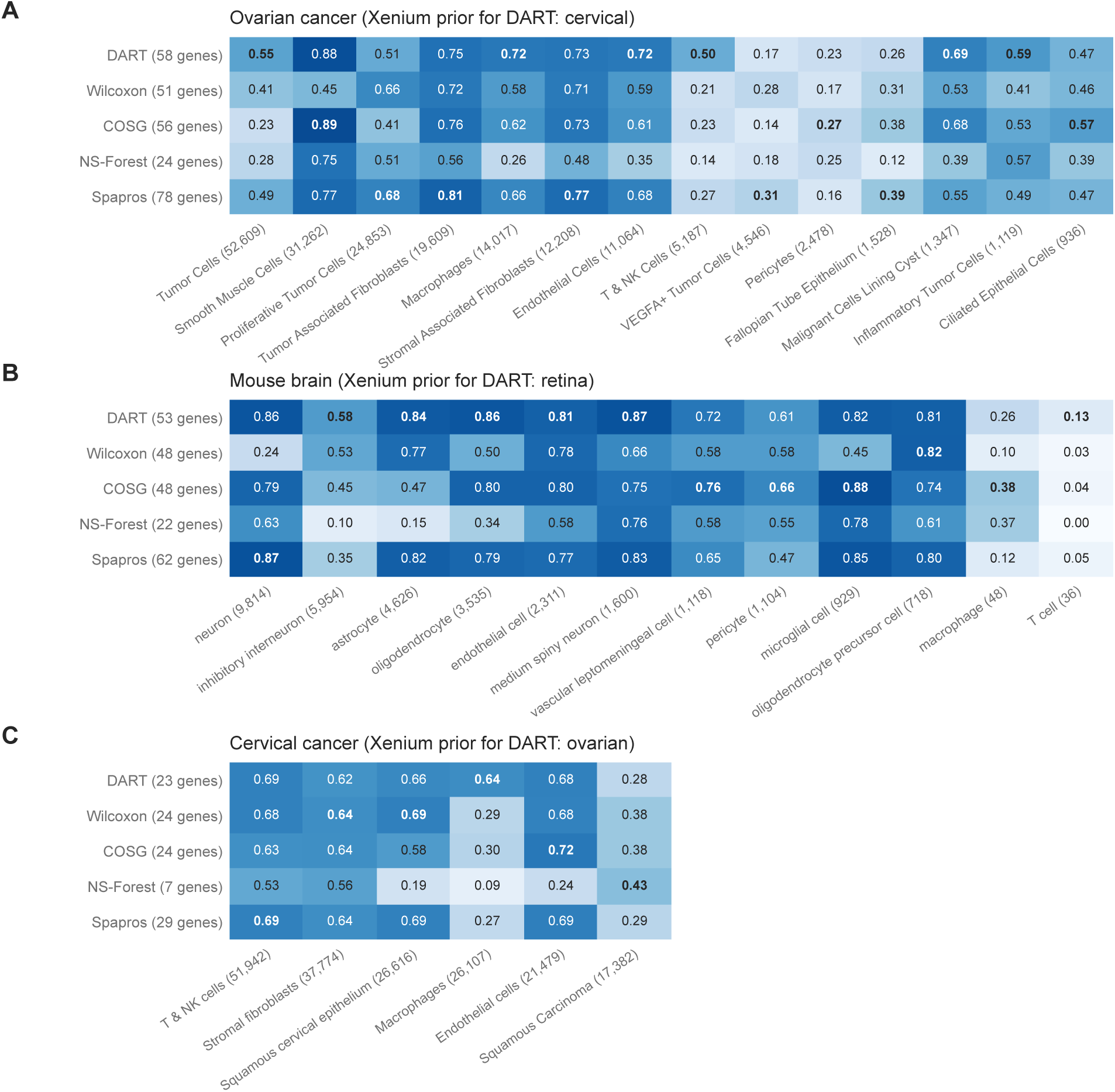
Per-cell-type *F*_1_ on each target Xenium. *F*_1_ of each cell type (columns, ordered by the number of held-out Xenium cells, in brackets) for one panel per method near DART’s default size (rows; panel size in brackets), as in Fig. 8a. Bold marks the best method for each cell type. **(A)** Ovarian cancer (cervical prior). **(B)** Mouse brain (retina prior; rebuilt labels). **(C)** Cervical cancer (ovarian prior).

### 2.9 Number of rules per cell type and rule overlap

Two settings control a DART panel: *K*, the number of rules kept per cell type, and *O*_max_, the number of genes that two rules of the same cell type may share (Fig. 10). Each additional rule adds genes, so *K* also sets the panel size. macro-*F*_1_ rose with *K* on every target, steeply from *K* = 1 to *K* = 3 and more slowly afterwards. On ovarian cancer it rose from 0.455 with one rule per type (18 genes) to 0.555 at the default *K* = 4 (58 genes) and 0.623 at *K* = 10 (128 genes); on mouse brain from 0.574 to 0.680 and 0.703; and on cervical cancer from 0.499 to 0.595, reaching 0.643 at *K* = 9 before falling to 0.629 at *K* = 10. The effect of letting rules share one gene (*O*_max_ = 1) depended on the target. On ovarian cancer it lowered macro-*F*_1_ at every *K* above 1, by 0.085 at *K* = 4. On mouse brain the two settings differed by at most 0.024, with no consistent sign. On cervical cancer sharing was better from *K* = 4 upward, by 0.014 at *K* = 4 and 0.043 at *K* = 10.

**Figure 10:**
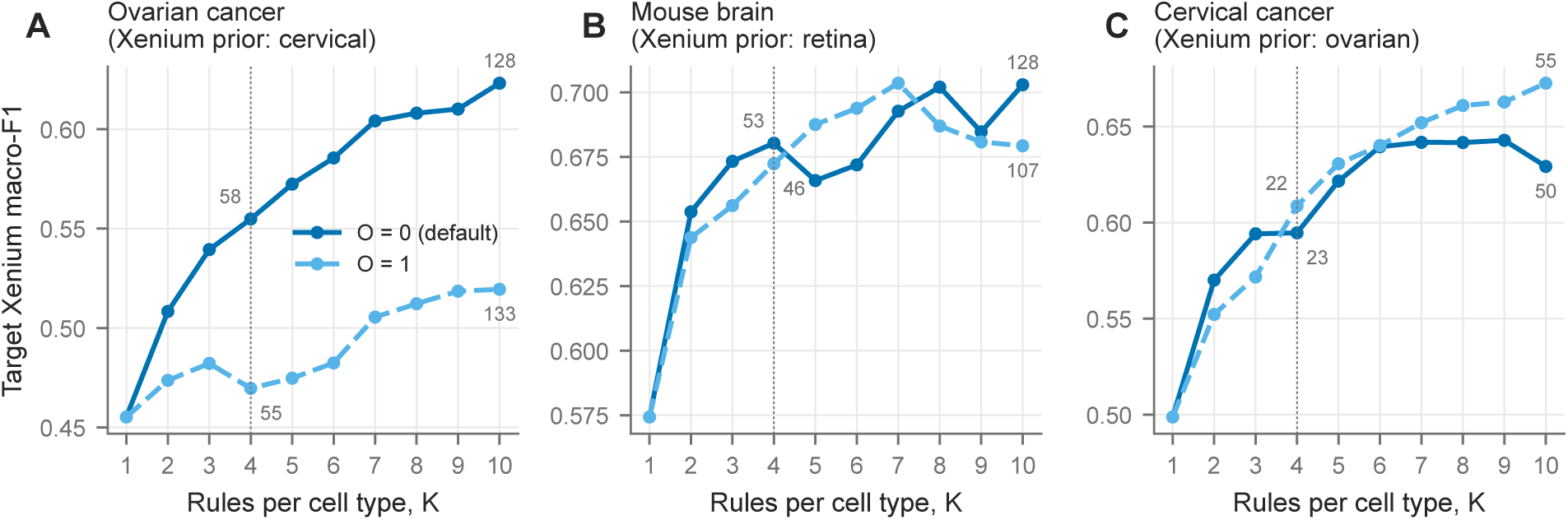
DART accuracy as a function of the number of rules per cell type and their overlap. macro-*F*_1_ on the held-out target Xenium against *K*, the number of rules kept per cell type, for gene-disjoint rules (*O*_max_ = 0, the default) and rules that may share one gene (*O*_max_ = 1), with each target’s Xenium prior. Numbers give the panel size at *K* = 4 and *K* = 10. The dotted line marks the default *K* = 4.

### 2.10 DART’s rules are interpretable and agree with the classifier

Each DART panel comes with explicit rules, such as a gene above a threshold, alone or combined with others. These rules name recognisable marker programs (Table 3). For the interferon-high inflammatory tumour cells of the ovarian cancer reference, DART’s four gene-disjoint rules use *MX1* (in a conjunction that also requires *ESR1* high and *NAMPT* and *CD200* low), *IFIT1*, *IFIT3* and *DDX58*, all interferon-stimulated genes. The rules for ciliated epithelial cells use cilia genes (*CFAP100*, *CDHR4*, *TEKT1*, *DNAH9*), those for proliferative tumour cells use cell-cycle genes (*CIP2A*, *RRM2*, *TYMS*, *TPX2*), and those for T and natural killer cells use *TRAC*, *CD8A*, *CXCR4* and *IL7R*. A ranked gene list carries no threshold and does not say which genes act together.

**Table 3:**
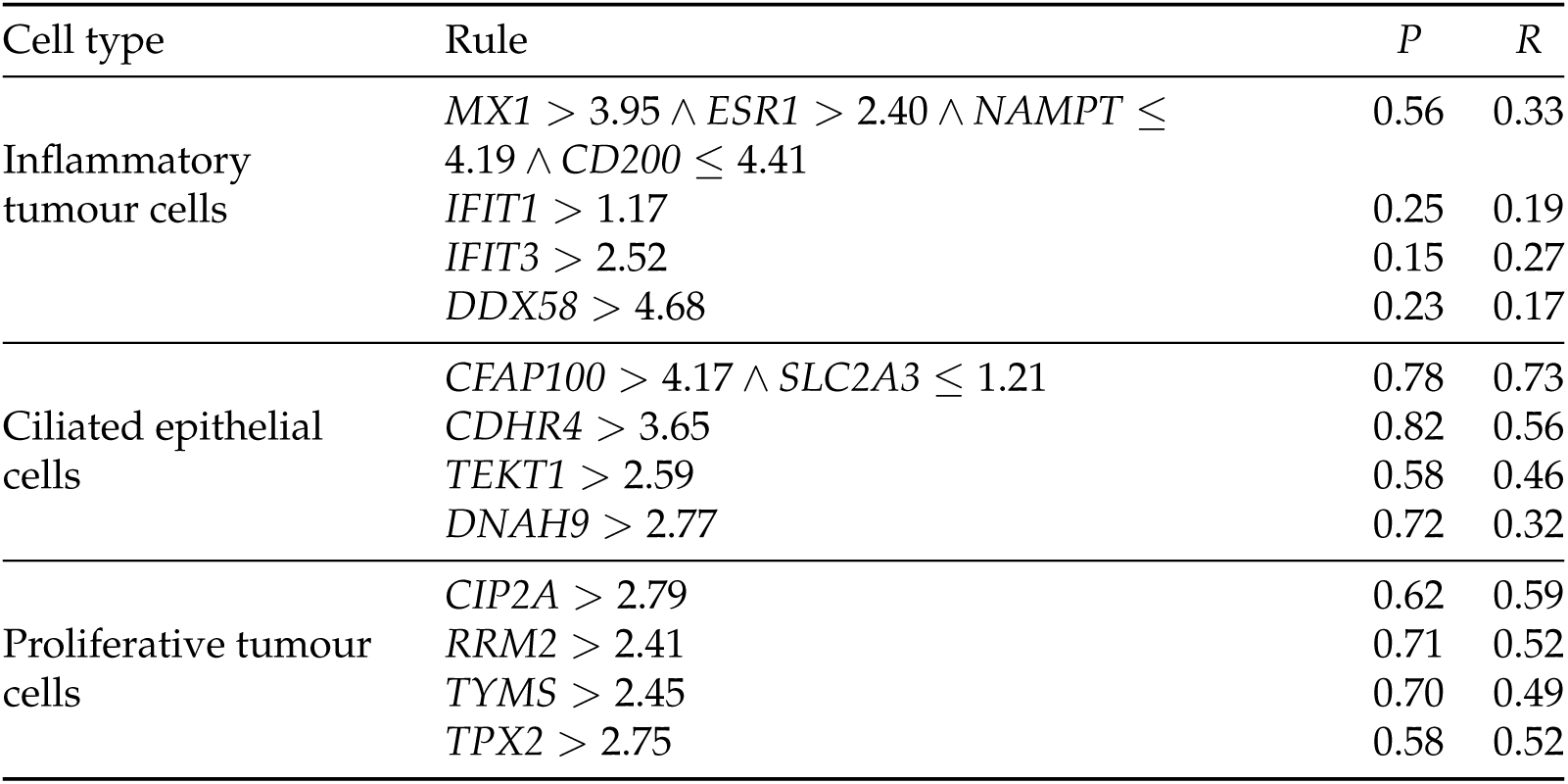
Example DART rules. The four gene-disjoint rules kept for selected cell types of the ovarian cancer reference (cervical Xenium prior, *K* = 4). Thresholds are on the log-normalised scale of the thinned reference. *P* and *R*, precision and recall of the rule in the thinned reference.

| Cell type | Rule | $P$ | $R$ |
| --- | --- | --- | --- |
| Inflammatory tumour cells | $MX1 > 3.95 \wedge ESR1 > 2.40 \wedge NAMPT \leq 4.19 \wedge CD200 \leq 4.41$ | 0.56 | 0.33 |
| | $IFIT1 > 1.17$ | 0.25 | 0.19 |
| | $IFIT3 > 2.52$ | 0.15 | 0.27 |
| | $DDX58 > 4.68$ | 0.23 | 0.17 |
| Ciliated epithelial cells | $CFAP100 > 4.17 \wedge SLC2A3 \leq 1.21$ | 0.78 | 0.73 |
| | $CDHR4 > 3.65$ | 0.82 | 0.56 |
| | $TEKT1 > 2.59$ | 0.58 | 0.46 |
| | $DNAH9 > 2.77$ | 0.72 | 0.32 |
| Proliferative tumour cells | $CIP2A > 2.79$ | 0.62 | 0.59 |
| | $RRM2 > 2.41$ | 0.71 | 0.52 |
| | $TYMS > 2.45$ | 0.70 | 0.49 |
| | $TPX2 > 2.75$ | 0.58 | 0.52 |

Panels are deployed with a classifier that never sees the rules, yet the classifier’s coefficients recover them (Table 4). A “*gene > t*” condition in a cell type’s rule carries a positive coefficient for that type in 98–100% of cases on the three Xenium targets and in 96.5% across the 45 RGC subtypes. Rule genes carry 2.0–3.0 times the weight of the panel’s other genes, and a cell type’s largest coefficient belongs to one of its rule genes for every cell type of the three targets and for 92% of the RGC subtype panels. The rules are therefore an accurate, human-readable account of why the classifier calls a cell type.

**Table 4:** DART’s rules agree with the deployed classifier. The classifier that scores each panel (Methods), for the benchmark’s *K* = 4 panel of each Xenium target and, for the RGC atlas, each fold’s *K* = 4 panel with that fold’s classifier, pooled over the five folds. “*>*” and “*≤*” are the fractions of rule conditions whose gene has a positive or negative coefficient, respectively, for the rule’s cell type (number of conditions in brackets). Weight ratio is the mean |coefficient| of a type’s rule genes over that of the panel’s other genes.

| Panel (cell types) | “$>$” → positive | “≤” → negative | weight ratio | top coefficient is a rule gene |
| --- | --- | --- | --- | --- |
| Ovarian (14) | 100% (57) | 83% (6) | 2.3× | 100% |
| Mouse brain (12) | 98% (49) | 86% (7) | 2.2× | 100% |
| Cervical (6) | 100% (20) | 100% (6) | 2.0× | 100% |
| RGC subtypes (45; 5 folds) | 96.5% (1,522) | 83.8% (597) | 3.0× | 92% |

### 2.11 Fine retinal ganglion cell subtypes

The Xenium targets have 6–14 broad cell types, which a single marker usually separates; on them most DART rules use one gene. RGCs, the projection neurons of the retina, comprise more than 40 transcriptionally related subtypes (Tran et al., 2019; Sanes and Masland, 2015), many without a unique marker. No Xenium dataset has RGC subtype labels, so we evaluated the 45 subtypes of a published atlas by five-fold out-of-fold prediction within the single-cell data (Methods): each panel is designed on four folds and scored on the held-out fold. Here 75% of DART’s rules are multi-gene conjunctions, with a mean of 2.4 genes per rule (*K* = 4 panels of the five folds), against 7–15% on the three Xenium targets.

DART, the Wilcoxon test and Spapros reached the same accuracy at matched size, about 0.955 to 0.96 macro-*F*_1_ from 150 genes upward (Fig. 11a): DART scored 0.959 with 184 genes and 0.960 with 223, the Wilcoxon test 0.957 with 191 and 0.958 with 224, and Spapros 0.956 with 175. COSG was 0.04–0.06 lower at matched size and reached 0.945 only with 393 genes, and NS-Forest’s single panel (71 genes) scored 0.879. DART led COSG (+0.028 to +0.037) and NS-Forest (+0.052 to +0.076) in every third of subtype abundance, from the rarest to the commonest, whereas against the Wilcoxon test and Spapros the per-subtype differences stayed within *±*0.007 at every abundance (Fig. 11b). No single probe was load-bearing, consistent with each subtype being covered by several gene-disjoint rules: among the 230 single-gene knockouts of one fold’s *K* = 4 panel, the largest drop in macro-*F*_1_ was 0.003.

**Figure 11:**
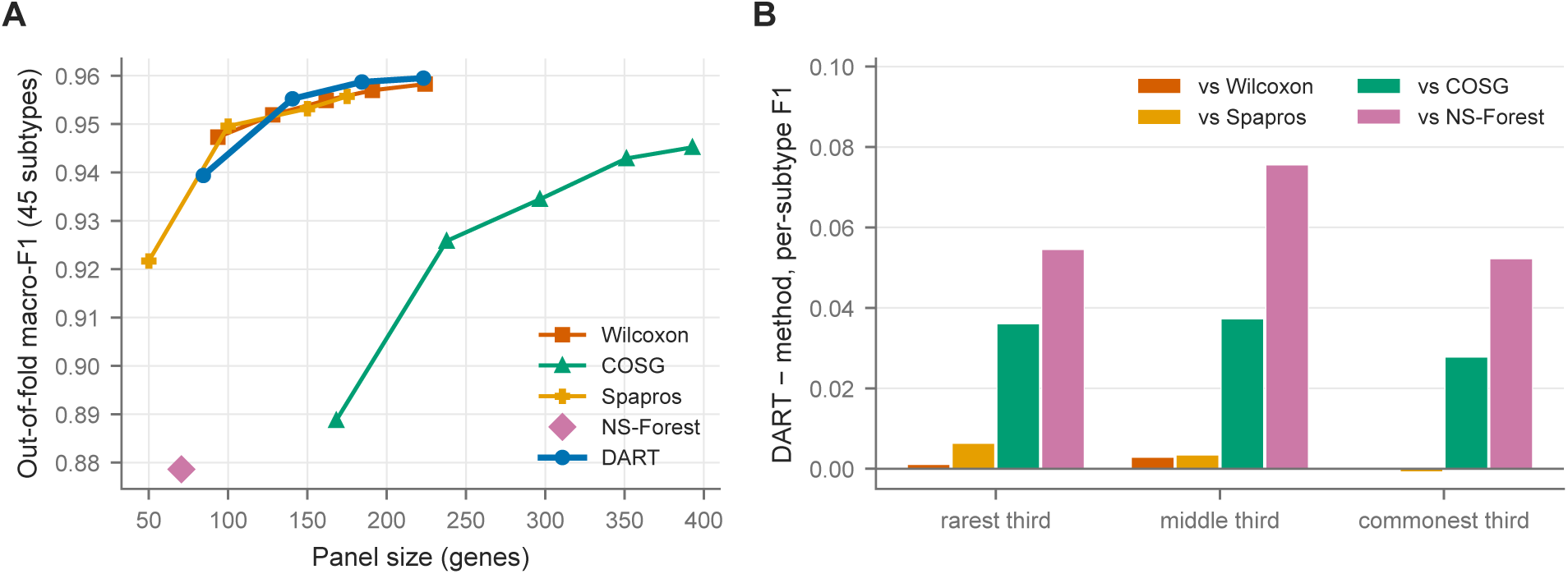
Forty-five RGC subtypes, held out within the single-cell atlas. **(A)** Out-of-fold macro-*F*_1_ against panel size for DART (*K* = 1–4), the Wilcoxon test and COSG (4–12 genes per subtype), Spapros (50, 100 and 150 genes requested, and its default of 170–182 genes) and NS-Forest. **(B)** Mean per-subtype *F*_1_ difference between DART and each method, by subtype abundance (thirds of the 45 subtypes), each at the closest panel sizes: DART with 223 genes against the Wilcoxon test with 224 and COSG with 238, DART with 184 against Spapros with 175, and DART with 84 against NS-Forest with 71.

### 2.12 Checking a panel’s probes for off-target expression

The off-target check applies to a panel from any method once its probes are designed. Fig. 12 shows its output for a 176-gene human retina panel designed in a separate study, using the probe report returned by the Xenium Panel Designer and a single-nucleus atlas of the human retina (Li et al., 2026). The probe for the cone marker *CRX* has one predicted off-target, *RPS6KL1*, which is expressed weakly and evenly across the retinal cell classes, so it cannot mimic the cone pattern and the probe can be kept. Three of the probes for the amacrine-cell marker *PAX6* have predicted off-targets. Of those measured in the atlas, *ALX1* is barely expressed, but *PAX2*, *PAX5* and *PAX8* are strongly expressed in astrocytes, where *PAX6* itself is absent. Those probes could report amacrine-like signal in astrocytes, so they are flagged for replacement.

**Figure 12:**
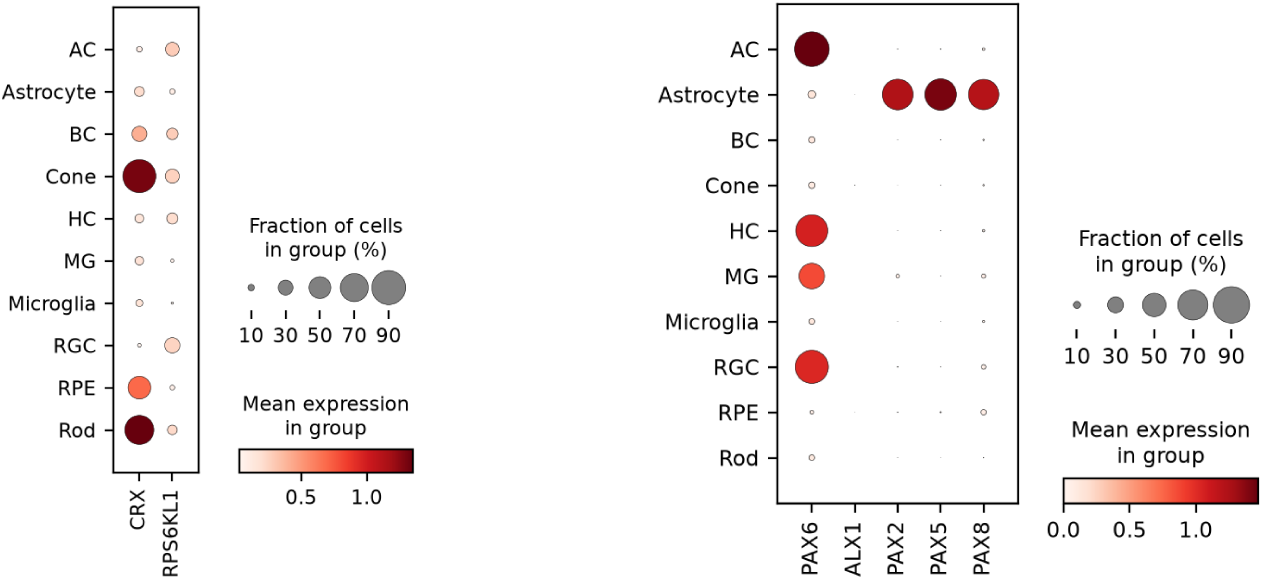
Example output of the off-target check. Dotplots of a marker (left column of each plot) beside its predicted off-target genes across the major cell classes of a human retina single-nucleus atlas (Li et al., 2026); dot size is the fraction of cells expressing the gene and colour its mean expression. **(A, left)** The cone marker *CRX*: its off-target *RPS6KL1* is expressed weakly and evenly across classes, so the probe is kept. **(B, right)** The amacrine-cell marker *PAX6*: of its off-targets, *PAX2*, *PAX5* and *PAX8* are expressed in astrocytes, so its probes are flagged. Panel from a 176-gene human retina design. AC, amacrine cells; BC, bipolar cells; HC, horizontal cells; MG, Müller glia; RGC, retinal ganglion cells; RPE, retinal pigment epithelium.

## 3 Discussion

X- SPEAR turns an annotated single-cell reference into a Xenium panel without any data from the target sample. It makes three contributions:

**1. Detection-aware design from a Xenium prior.** The gap between a gene’s detection in a single-cell reference and in Xenium is large, gene-specific and reproducible across tissues. Thinning the reference to a previous experiment’s detection rates anticipates that gap before ordering. The prior should match the planned experiment’s chemistry and species, and it can come from another tissue; a mismatched prior can lower accuracy.
**2. DART.** With a Xenium prior from a different tissue, DART outperforms the Wilcoxon test, COSG, NS-Forest and Spapros at matched panel size on every target we tested, and reaches the Wilcoxon test’s best accuracy with 42–69% fewer genes. Its rules name interpretable marker programs and agree with the classifier that uses the panel. On fine RGC subtypes it matches the best methods, leads COSG and NS-Forest at every subtype abundance, and no single probe is load-bearing.
**3. A method-independent off-target check.** Dotplots of each marker beside its predicted off-target genes show which probes would add signal in the wrong cell types, for a panel chosen by any method.

Two limitations matter most. First, the benefit depends on a suitable Xenium prior: for the best results, the prior should come from the same assay chemistry and the same species as the target. Second, the evaluation labels of the Xenium targets derive from reference-based annotation: 10x’s for ovarian and cervical cancer and our own label transfer for mouse brain. The comparisons are therefore within the reference’s annotation scheme, although an independent annotation of the ovarian section agreed with the evaluation labels on 82.8% of cells at the level of five lineages (Supplementary S3).

The thinning, selection and off-target modules are separable. Other targeted platforms can be supported once a detection profile and a probe-design output exist for them, and the same architecture can host objectives beyond cell-type identification, such as panels that preserve differential expression across conditions or spatial niches.

## 4 Methods

### 4.1 The X-SPEAR software

#### Interfaces

X-SPEAR is a Python package (Code availability statement) with three interfaces. As a library, design_panel takes an annotated single-cell reference, a dataset configuration and, optionally, a Xenium prior, and returns the panel’s genes, the selected rules (each with its cell type, thresholds, precision and recall) and the full pool of candidate rules. Panel.reselect selects again from that pool at another *K* or *O*_max_ without relearning; this study used it for the RGC panels with *K* = 1–3. classify labels a target’s cells with a panel, and metrics.evaluate scores the labels. A dataset configuration, one JavaScript Object Notation (JSON) file, names the reference’s cell-type column, the cell types to leave out, and the renames and exclusions that map the target’s labels onto the reference’s. On the command line, xspear design, xspear classify and xspear evaluate wrap these calls. xspear prior lists and exports the Xenium priors shipped with the package, xspear profile depth-normalises priors and xspear kcurve plots accuracy against *K* (Supplementary S6). xspear init writes annotated configuration files for the pipeline, and xspear run runs it, in full or for chosen stages.

#### Pipeline

The stages of the pipeline (Fig. 3) pass files to each other. Every selector writes the panel’s gene list, which is all that stages 08–11 read, and DART also writes its rules. Each stage records its outcome, with the package version and commit, in a run manifest, and a stage does not start if an earlier one failed. Stage 00 loads counts (an AnnData h5ad file or a Matrix Market (MTX) matrix) and can attach Ensembl gene identifiers from a gene transfer format (GTF) file or mygene.info (needed for the export bundle), merge per-cell metadata and assign a group label such as the sample of origin. Stage 01 computes quality metrics and keeps cells with 500–8,000 detected genes and genes detected in at least three cells; it can also filter on mitochondrial fraction and call doublets with Scrublet (Wolock et al., 2019). Stage 02 caps the cells per cell type (3,000, seeded). Stage 03 normalises counts per 10,000, or per median library size, applies log(1 + *x*) and can flag highly variable genes. Stage 04, off by default, computes principal components, a neighbour graph, a uniform manifold approximation and projection (UMAP) (McInnes et al., 2018) and Harmony integration (Korsunsky et al., 2019). Stage 05 loads the target Xenium and its labels, which are used only for scoring. Stage 06 resolves a Xenium prior and thins a copy of the reference to it (Supplementary S6); every data-driven selector of stage 07 uses the thinned copy, and the classifier of stage 08 uses the original. Stage 07 runs DART as four substages, 07a rule learning, 07b rule scoring, 07c ranking and 07d orthogonal selection (Sections 4.3 and 4.4; defaults *K* = 4, *O*_max_ = 0 and 1,000 estimators per ensemble, as in this study). It can instead run the Wilcoxon test, COSG or NS-Forest (10 genes per cell type by default), read a curated marker list, run Spapros or its reference probe sets (principal components, differential expression, highly variable genes and random genes), or adopt an existing panel. Stage 10 writes the panel and, for the Xenium Panel Designer, a reference bundle of counts, gene and cell tables and cell types. Stage 11, off by default, is the off-target check (Section 4.9). This study ran stage 00 to load the references of the three Xenium targets and called the functions behind stages 06, 07 and 11 from its analysis scripts; the capping, splitting and scoring described below are in the paper repository.

#### Classification and evaluation

Stage 08 trains a classifier on the reference restricted to the panel and labels the target’s cells. The default is a one-versus-rest logistic regression (*C* = 1, at most 1,000 iterations, unweighted), as in this study’s scoring; a random forest (500 trees, depth 8) and balanced class weights are options. Reference and target are first normalised over the genes they share, as in this study’s scoring, and so does the library’s classify; normalising over the panel’s genes alone, which is what a Xenium run measuring only the panel provides, is an option. Cells can also be labelled by voting with DART’s rules instead of a classifier (Supplementary S5). Stage 09 reports macro-*F*_1_ and weighted *F*_1_ over the reference’s cell types, per-type precision, recall and *F*_1_, the confusion matrix and the fraction of cells assigned a type. An alternative evaluation built on Spapros’ forest classifier is described in Supplementary S7.

#### Environments

The analyses ran in Python 3.11.15 with numpy 2.4.6, pandas 2.3.3, SciPy 1.17.1, scikit-learn 1.9.0 (Pedregosa et al., 2011), scanpy 1.10.3 (Wolf et al., 2018), anndata 0.12.19, imodels 3.0.0 (Singh et al., 2021), Cython 3.3.0 (for the compiled ATPF criterion), COSG 1.2.0 and NS-Forest 4.1. Spapros 0.1.6 cannot run on numpy 2 or pandas 3, so X-SPEAR runs it in a second environment (numpy 1.26.4, pandas 2.3.3) as a subprocess that returns only the panel file. The package’s installer builds both environments, pinned to lock files of exactly these versions.

### 4.2 Datasets and preprocessing

Table 1 lists every dataset. For each target the candidate gene pool is the set of genes measured in both the single-cell reference and the Xenium panel: 4,766 (ovarian cancer), 4,548 (cervical cancer) and 4,791 (mouse brain). Expression is log(1 + *x*) of counts per 10,000 (CP10k), computed over this shared universe for both modalities before subsetting to a panel. For the benchmark, the reference was capped at 40,000 cells by stratified sampling, keeping small cell types whole, and the selectors were run on a stratified 80% of the capped reference.

The target Xenium cells were divided into design and evaluation halves by a spatial checker-board, and only the evaluation half was scored: about 183,000 cells for ovarian cancer, 181,000 for cervical cancer and 32,000 for mouse brain. The design half was not used.

#### Evaluation labels

For ovarian and cervical cancer, the labels are 10x Genomics’ cell-type annotation, mapped to the reference vocabulary. For cervical cancer, 10x left 56% of cells unassigned, and those cells were excluded. For mouse brain, the labels come from our label transfer, a scaled multinomial logistic regression trained on the reference over the shared genes. The original transfer normalised the reference over its full transcriptome but the Xenium over its panel; rebuilding it with consistent normalisation changed the labels of 18.3% of evaluation cells, and mouse brain results use the rebuilt labels except where noted (Supplementary S3). Labels are used only to score.

### 4.3 Rule generation

#### ATPF

ATPF grows scikit-learn (Pedregosa et al., 2011) decision trees with a custom precision– recall impurity, *I*(*n*) = 1 *− P × R*, where *P* is precision and *R* recall. This criterion targets isolation of one cell type rather than global class separation. Root-to-node paths are extracted as candidate rules.

#### RULEFIT

RULEFIT (Friedman and Popescu, 2008) (imodels 3.0.0) is applied to random-forest and gradient-boosted ensembles. Rules with nonzero coefficients in the *L*_1_-penalised logistic model are kept.

Each ATPF tree is grown on a bootstrap sample of the thinned reference, to depth 4, considering the square root of the number of genes at each split. Every root-to-node path, not only those that end in a leaf, is a candidate rule, and rules with the same conditions are merged. The ATPF criterion is a compiled extension of scikit-learn, built for the installed Python and scikit-learn versions. RULEFIT uses imodels’ RuleFitClassifier on a random forest with balanced class weights or on a gradient-boosted ensemble with 80% subsampling, both of depth 4, with up to 4,000 rules at 1,000 trees. The benchmarks used 1,000 estimators per ensemble. All candidate rules are evaluated on the full thinned design slice to give *P*, *R* and *F*_1_ (and *F_β_* for *β* = 0.5 and 2, which are not used). A cell type with fewer than five reference cells is skipped, and a cell type that receives no rule is reported as an error, after which the pipeline stops. Cell types are learned independently and can be spread over worker processes; each learner is seeded per cell type, so the candidate pool does not depend on the number of workers.

Three scoring options, not used in this study, change the candidate pool (stage 07b): discarding rules with a “*≤*” condition; recording each rule’s *F*_1_ against each other cell type alone, computed on the cells of the two types, whose minimum can replace one-versus-rest *F*_1_ in ranking; and rewriting each threshold as the fraction of reference cells at or below it, so that a rule fires at the same rank of any data it is applied to (Supplementary S5).

### 4.4 Rule ranking and orthogonal selection

ATPF candidates pass a quality gate: *P ≥* 0.75 and *R ≥* 0.50; if none pass, *P ≥* 0.45 and *R ≥* 0.30; if still none, no gate. The 20 best by *F*_1_ are then kept for each rule length. The top 40 RULEFIT rules per cell type by penalised *F*_1_ are kept. Pooled candidates are ranked by a penalised score,

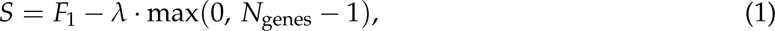

in two tiers. Rules with *F*_1_ *≥* 0.7 are ranked first, with *λ* = 0.10, and the rest follow with *λ* = 0.04. Ties are broken by *|P − R|* and then by rule length. For each cell type, rules are added greedily in rank order until *K* have been kept. A candidate is skipped if it shares more than *O*_max_ genes with the rules already chosen for that type, or if all its genes are already among them. Genes may be shared across cell types. The panel is the union of the selected rules’ genes. Defaults are *K* = 4 and *O*_max_ = 0; the selection constants sit on a plateau (Supplementary S1).

The tier threshold, both penalties and the order of tie-breaking can be changed, and the candidate pool can be kept before the quality gate, so that the gate, the number of rules kept per rule length and the RULEFIT top-*N* can be varied without relearning. Supplementary S1 used these, together with an option that keeps every gene to one cell type, with the types choosing rules in turn. Two further selection modes, not used here, are available from the library: a candidate’s score can be reduced by *λ* times the largest absolute Pearson correlation between its genes and those already on the panel, and a total gene budget can replace *K*, spent first on each type’s best rule that fits and then repeatedly on the rule that most raises its type’s best *F*_1_ per gene it adds. Selection can also be restricted to the genes the target Xenium measures, or to a list such as the genes that can be ordered on a panel.

### 4.5 Detection-aware thinning with a Xenium prior

For each gene *g*, a Xenium prior gives

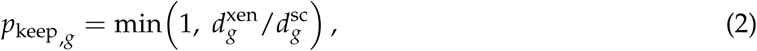

where 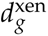 and 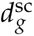 are the fractions of all cells with a nonzero count in the prior’s Xenium run and single-cell reference. Before learning, each nonzero entry of the reference count matrix is kept intact with probability *p*_keep,*g*_ and set to zero otherwise. This per-entry thinning reproduces the prior’s per-gene detection rate; binomial thinning of individual molecules would not, because it barely changes detection for well-expressed genes. The thinned counts are then normalised as above. The prior is used at its own magnitude and its labels are not used. Genes are matched between prior and reference by gene symbol, and a gene the prior does not cover is not thinned. This study’s ovarian cancer prior ships with X-SPEAR (Supplementary S6).

### 4.6 Xenium prior comparison

For each target, DART panels were designed with each Xenium prior, with the target’s own detection profile (an upper bound) and with no prior, exactly as in the benchmark: the same capped reference and design slice, 1,000 estimators per ensemble, *K* = 1–10 and *O*_max_ = 0, scored with the benchmark classifier on the held-out target Xenium. The panels designed with each target’s benchmark prior are therefore the benchmark’s DART panels. Here gene symbols were matched ignoring case, so that a prior from the other species reaches the genes whose human and mouse symbols agree (for example human *GFAP* and mouse *Gfap*): 2,139–2,348 of the target’s 4,548–4,791 genes (45–49%). X-SPEAR itself matches symbols exactly, which is the same for priors of the target’s species. Because panel size at a given *K* differs between priors, macro-*F*_1_ was interpolated along each prior’s curve at the realised sizes of the target’s benchmark panels with *K* = 3–6 (45–83 genes for ovarian cancer, 41–79 for mouse brain and 17–32 for cervical cancer) and averaged.

### 4.7 Benchmark protocol

#### Methods compared

We used the Wilcoxon rank-sum test (scanpy 1.10.3 (Wolf et al., 2018); top-*k* genes per cell type), COSG 1.2.0 and NS-Forest 4.1, at *k ∈ {*1, 2, 3, 4, 6, 8, 10, 12*}* genes per cell type for the Wilcoxon test and COSG and *k ∈ {*2, 4, 8*}* for NS-Forest. Spapros 0.1.6 ran through X- SPEAR’s Spapros interface, in its own environment (Section 4.1), at its default size; as X-SPEAR prepares it, it saw log(1 + *x*) CP10k expression with 4,000 highly variable genes flagged. Every method selected from the same cells, the stratified 80% slice of the capped reference (Section 4.2), and from the single-cell reference as measured. DART used the Xenium prior described in Section 2.6.

#### Scoring

Every panel in this paper, on Xenium and out of fold, was scored by one procedure. The training cells and the test cells are normalised over the genes they share and then restricted to the panel’s genes; a one-versus-rest logistic regression (*C* = 1, unweighted) is trained on the training cells and predicts every test cell. For the Xenium targets the training cells are the full single-cell reference without thinning and the test cells the held-out target Xenium half. macro- *F*_1_ averages per-cell-type *F*_1_ over the reference’s cell types; weighted *F*_1_ weights each type by its number of held-out cells. Comparisons at matched size interpolate each method’s macro-*F*_1_ linearly in realised panel size and average the difference over DART’s sizes that lie within the other method’s range. NS-Forest and Spapros give no curve, so DART’s curve was interpolated at their panel sizes; NS-Forest’s margin is the mean over its three budgets.

#### Seed variability

The reference thinning and every learner were repeated at four seeds (1, 2, 3 and 42; the benchmark uses 42), and each seed’s DART curve (*K* = 1–6 and 8) was compared with the other methods’ panels exactly as in the benchmark. Across seeds, the standard deviation of DART’s margin over a method was 0.005–0.017 macro-*F*_1_ (0.006–0.008 over the Wilcoxon test), and that of DART’s macro-*F*_1_ at a single *K* was up to 0.024. A DART panel’s genes partly change between seeds (mean pairwise Jaccard index 0.62–0.74 at *K* = 4).

#### Compute

Compute was measured on this study’s own runs, one job per target and method on the same cluster. For DART, the Wilcoxon test, COSG and NS-Forest, the job loaded the benchmark’s design slice and timed the method’s selection call exactly as the benchmark makes it (DART with the target’s Xenium prior, *K* = 4, *O*_max_ = 0 and 1,000 estimators; DART and NS-Forest with eight worker processes), recording the wall time of the call, the CPU time of the process and its workers, and the peak memory of the largest process. For Spapros, the scheduler summary of its job gave the same quantities, which also cover loading and preparing the data.

### 4.8 Single-cell out-of-fold evaluation on RGC subtypes

The RGC atlas (34,029 cells, 45 subtypes) was restricted to the 4,911 genes of the mouse Xenium 5K panel and split into five folds, stratified by subtype (scikit-learn StratifiedKFold, shuffled, seed 42). For each fold, every method selected its panel from the other four folds only, so the held-out cells never influenced their own panel. Each panel was scored as on Xenium (Benchmark protocol), with the four training folds as the training cells and the held-out fold as the test cells. The predictions of the five folds were pooled, and macro-*F*_1_ and per-subtype *F*_1_ were computed over all 34,029 cells. No Xenium prior was used, because there is no Xenium measurement to anticipate. DART was run with *K* = 4 rules per subtype and *O*_max_ = 0 at 1,000 estimators, and smaller panels were obtained as the first *K* = 1–3 rules of each fold’s *K* = 4 selection; the Wilcoxon test and COSG were run at 4, 6, 8, 10 and 12 genes per subtype and NS-Forest at its default, and Spapros was run on each fold’s training cells exactly as for the Xenium targets, at its default size (170–182 genes; a request for 200 genes returned the same panel) and at 50, 100 and 150 genes. Per-subtype differences were compared at near-matched panel sizes (Fig. 11b), and subtypes were split into thirds by their number of cells. For the knockout analysis, each gene of a fold’s *K* = 4 panel was removed in turn, the classifier was retrained without it, and the drop in macro-*F*_1_ on the held-out fold was recorded.

### 4.9 Probe off-target check

After probe design, the Xenium Panel Designer returns a probe report listing, for each probe, its predicted off-target genes. X-SPEAR separates probes with and without predicted off-targets, attaches the cell type each marker was selected for, and, for each marker with off-targets, draws a dotplot of the marker and its off-target genes across the cell types of the annotated reference (scanpy; expression as CP10k then log(1 + *x*)). The user replaces probes whose off-target genes are expressed in cell types the marker does not mark. Fig. 12 was drawn this way by X-SPEAR’s off-target stage, with the human retina atlas’s major cell classes (up to 3,000 cells each) as the reference.

## Supporting information

Latex source

## Statements

### Statement of ethics

This study analysed only publicly available, de-identified human datasets and published mouse datasets (see the Data availability statement). No new human or animal data were generated for the results reported here, so institutional review was not required.

### Author contributions

RP developed DART, built most of X-SPEAR, performed the benchmarking and analyses, and drafted the manuscript. RP and BD packaged X-SPEAR as a Python package. VP prepared the RGC single-cell reference, verified it against the public GEO data, and identified the normalisation default that X-SPEAR’s rule voting now corrects. YC conceived and supervised the study. All authors reviewed and approved the manuscript.

### Data availability statement

All datasets are public:

- ovarian cancer: 10x Genomics https://www.10xgenomics.com/datasets/17k-human-ovarian-cancer-scFFPE and https://www.10xgenomics.com/datasets/xenium-prime-ffpe-human-ovarian-cancer;
- cervical cancer: 10x Genomics https://www.10xgenomics.com/datasets/17k-human-cervical-cancer-scFFPE and https://www.10xgenomics.com/datasets/xenium-prime-ffpe-human-cervical-cancer;
- mouse brain: 10x Genomics https://www.10xgenomics.com/datasets/xenium-prime-fresh-frozen-mouse-brain, with the single-nucleus reference from CZ CELLxGENE https://cellxgene.cziscience.com/collections/31937775-0602-4e52-a799-b6acdd2bac2e;
- mouse retina: Gene Expression Omnibus (GEO) GSE325478 (reference; sample GSM9604550) and GSE307031 (Xenium; sample GSM9214539);
- breast cancer: GEO GSM7782698 and Zenodo 10076046 (reference) and 10x Genomics https://www.10xgenomics.com/datasets/xenium-prime-ffpe-human-breast-cancer (Xenium);
- RGC atlas: GEO GSE133382 (Tran et al., 2019);
- STHELAR annotations (Supplementary S3): BioStudies S-BIAD2146;
- human retina atlas (off-target example): CZ CELLxGENE https://cellxgene.cziscience.com/collections/4c6eaf5c-6d57-4c76-b1e9-60df8c655f1e.

The one input that is not public is the probe report of the human retina panel in Fig. 12, which belongs to an unpublished study by the authors’ laboratory.

### Code availability statement

X-SPEAR is an MIT-licensed Python package at https://github.com/yuyanchenglab/xspear; the version described here is commit bb86b4a. The Methods and Supplementary S5–S7 describe its interfaces, pipeline stages and options. The analysis code, result tables and figure scripts are at https://github.com/yuyanchenglab/xspear-paper.

## Acknowledgements

Support for this work was provided by the unrestricted Research to Prevent Blindness (RPB) departmental fund to the Scheie Eye Institute. The funder had no role in the design, data collection, data analysis, and reporting of this study.

Computation resources were generously provided by Penn Medicine Academic Computing Services (PMACS), whose High-Performance Computing (HPC) and Archive systems were funded by National Institutes of Health (NIH) grant 1S10OD012312.

