## Supplemental for "X-SPEAR: An Extensible Framework for Spatial Panel Evaluation and Refinement"

References cited here are listed in the main manuscript. Figure, table and section numbers without an "S" refer to the main manuscript.

#### S1. DART's selection constants are on a plateau

We swept every inherited selection constant one at a time (29 configurations), each with and without cross-type gene sharing, over $K = 1$–8:

- the ATPF quality gate;
- the number of rules kept per rule length;
- the RULEFIT top-$N$;
- the tier-1 $F_1$ threshold;
- both length penalties;
- the tie-break rule;
- $O_{\max} \in \{0, 1\}$.

Configurations were judged at fixed realised panel sizes, with each target's Xenium prior. Only the length penalty and $O_{\max}$ were load-bearing. Removing the length penalty roughly doubled the share of multi-gene rules, added about 55 genes and cost 0.045 on ovarian cancer (0.014 on mouse brain) even at matched size; Fig. 10 shows the effect of $O_{\max}$. Every other variation was within $\pm 0.01$ of the default on ovarian cancer and mouse brain. A design-time proxy, a held-out slice of the reference thinned by the Xenium prior, could not resolve differences inside this plateau (its ranking of configurations correlated with the target Xenium at Spearman $-0.23$ on ovarian cancer and $+0.12$ on mouse brain). Choosing the constants on the *other* datasets' Xenium, which the deployment rule permits, changed accuracy by only $+0.004$ macro-$F_1$. We therefore ship fixed defaults. The mouse brain numbers in this section use the original labels.

#### S2. Detection profiles: sampling noise, composition and normalisation

$p_{\text{keep}}$ is estimated from a finite number of cells, so it carries sampling noise. To measure it, we resampled with replacement the cells of the single-cell reference and of the Xenium cells the prior was computed from (the design half of each run), recomputed $p_{\text{keep}}$, and compared it with the estimate from all cells, over the genes detected in at least 1% of reference cells (200 resamples). One resample differs from the full estimate by a root-mean-square of 0.018 for ovarian cancer (16,187 reference cells, 3,497 genes) and 0.024 for cervical cancer (9,475 cells, 3,276 genes), against a standard deviation of $p_{\text{keep}}$ across genes of 0.29 and 0.26 (Fig. S1). The noise is small next to the differences between genes on which thinning acts.

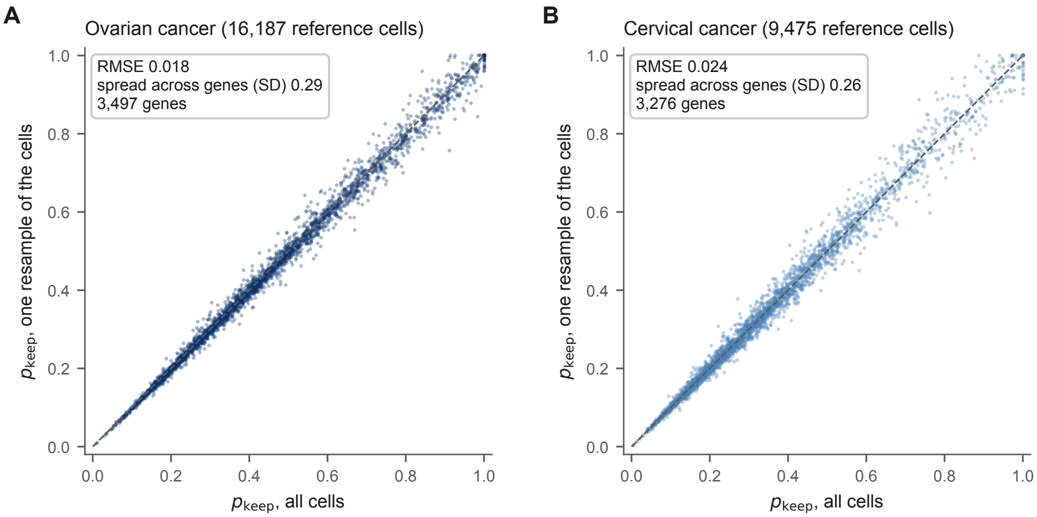


**Figure S1: Sampling noise of the detection-rate ratio $p_{\text{keep}}$.** $p_{\text{keep}}$ from one resample of the reference and Xenium cells against the estimate from all cells, for the genes detected in at least 1% of reference cells, for ovarian (A) and cervical cancer (B); dashed line, $y = x$. Each panel gives the root-mean-square error (RMSE) of one resample, averaged over 200 resamples, and the standard deviation (SD) of $p_{\text{keep}}$ across genes.

Because the profile is computed from all cells, it partly reflects cell-type composition. Reweighting within-type detection to the reference's composition moves $p_{\text{keep}}$ by a median of 0.017 on ovarian and 0.019 on cervical cancer (Spearman 0.977 and 0.985 with the all-cell profile).

The reference's stored expression is normalised over its full transcriptome, while our benchmark normalises over the shared universe. The Wilcoxon test is insensitive to the difference (mean macro-$F_1$ within $\pm 0.003$). COSG is not: the full-transcriptome basis raises it by 0.015 on ovarian and 0.036 on cervical cancer, and lowers it by 0.003 on mouse brain (original labels). Comparisons with COSG should be read with this in mind.

### S3. Evaluation labels: provenance, validation and an independent annotation

**Provenance.**

- *Ovarian and cervical cancer.* The labels are 10x Genomics' annotation, mapped to the reference vocabulary. Cell types absent from the reference were removed, and for cervical cancer the 56% of cells 10x left unassigned were excluded.
- *Mouse brain.* The labels are our own transfer from its reference. The original transfer normalised the reference over 20,409 genes and the Xenium over its 5,006-gene panel. Rebuilding it with both sides normalised over the same genes changed 18.3% of evaluation-cell labels, all in one direction. Implausible immune and meningeal calls in a healthy coronal section fell sharply (T cells 770 to 36; vascular leptomeningeal cells 3,515 to 1,118), and the cells became neurons. Re-scoring all 138 mouse brain runs of our pipeline on the rebuilt labels kept their order (Spearman 0.981) and raised macro-$F_1$ by a median of 0.062. Mouse brain numbers in this paper use the rebuilt labels except in S1 and S2, as noted.

**Validation.** Fig. S2 checks the *true* labels of the ovarian and cervical Xenium cells, that is 10x's annotation used for scoring, not any panel's predictions. For each labelled Xenium cell type (evaluation half) we compared its mean expression profile with that of every single-cell reference cell type over the shared genes, both normalised as for scoring. The matrices are diagonal-dominant: the mean same-type Spearman correlation is 0.83 for ovarian and 0.82 for cervical cancer, and each Xenium type's best match is its own reference type for all 6 cervical and 11 of 14 ovarian types. The three exceptions are tumour states (tumour, proliferative tumour and inflammatory tumour cells) that match the closely related VEGFA-positive tumour cells as well as or slightly better than their own type (by at most 0.016).

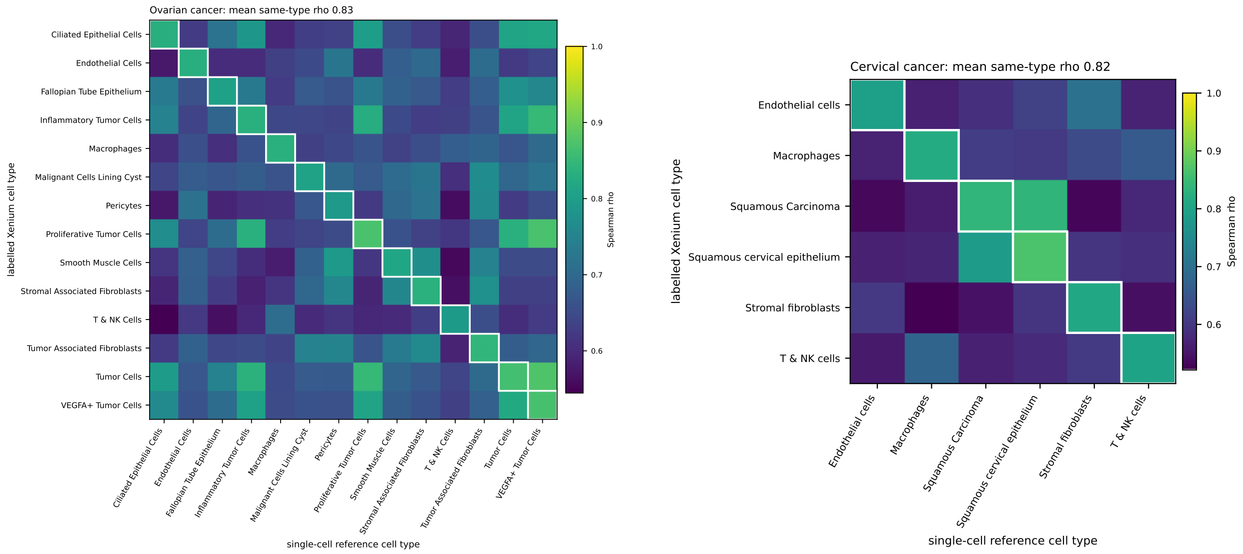


**Figure S2: Validation of the true Xenium labels used for scoring.** Spearman correlation between the mean log-normalised profiles of single-cell reference cell types and of the labelled Xenium cell types (10x's annotation on the evaluation half, not panel predictions) over shared genes, for ovarian (left) and cervical cancer (right). Boxes mark each Xenium type against its own reference type. Mean diagonal $\rho$ is 0.83 and 0.82. The best match is the type's own reference type for 11 of 14 ovarian and 6 of 6 cervical types.

**Independent annotation.** The STHELAR resource provides an annotation of the same ovarian Xenium section, at the level of five lineages, that was produced independently of 10x's (BioStudies S-BIAD2146). We matched its cells to our evaluation cells by mutual nearest cell centroid (98.6% of cells matched). With the evaluation labels coarsened to the same five lineages, the two annotations agree on 82.8% of cells.

#### S4. Compute cost

Table S1 lists the values plotted in Fig. S3.

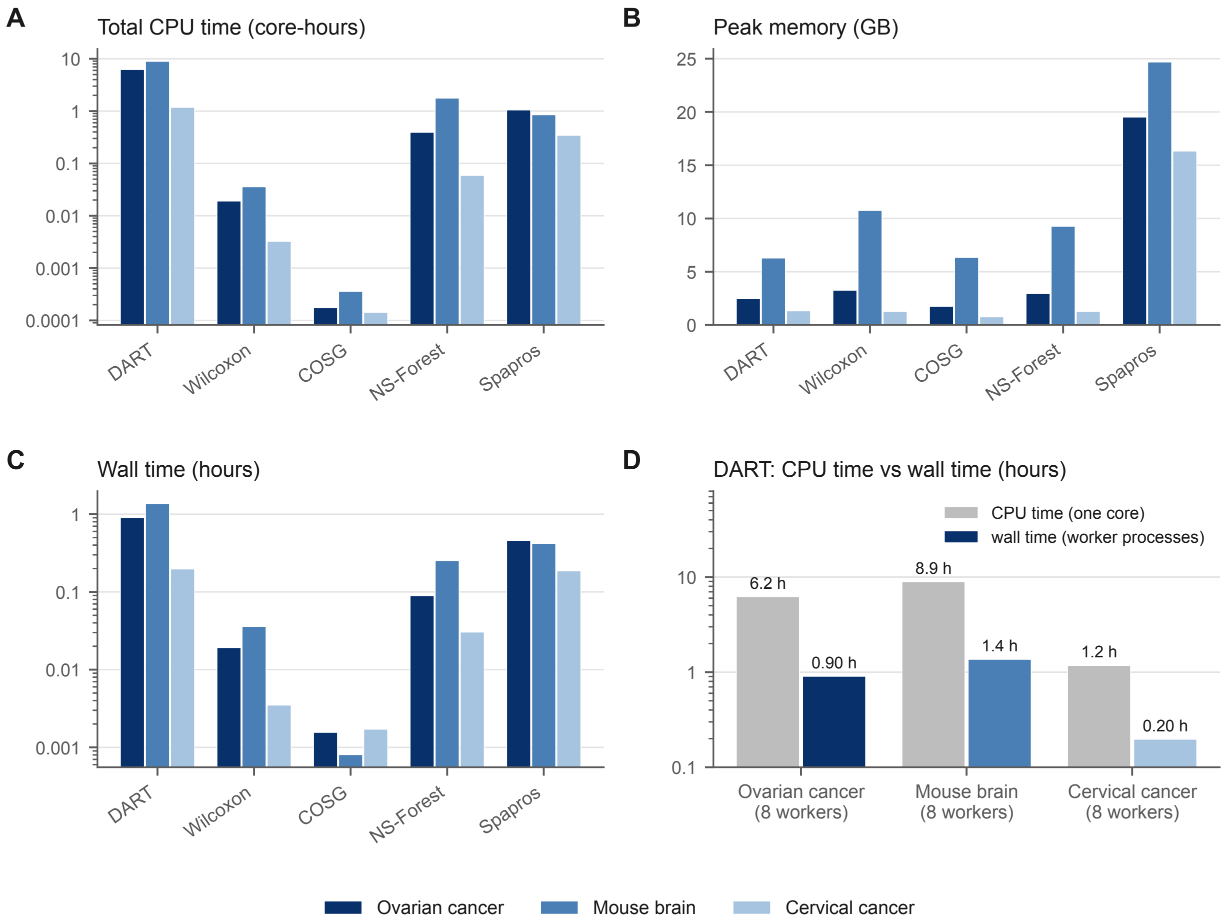


**Figure S3: Compute cost per panel-design method.** From this study's runs on the three targets (Methods). **(A)** Total CPU time, including worker processes. **(B)** Peak memory of the largest process. **(C)** Wall time. **(D)** DART's CPU time against its wall time with eight worker processes. (A), (C) and (D) use log scales. Exact values are in Table S1.

**Table S1: Compute cost per panel-design method on each target.** Worker processes, CPU time (core-hours), peak memory of the largest process (GB) and wall time (hours), from this study's runs (Methods). Spapros's values also cover loading and preparing the data.

| | Ovarian cancer | | | | Mouse brain | | | | Cervical cancer | | | |
|---|---|---|---|---|---|---|---|---|---|---|---|---|
| Method | workers | CPU (h) | memory (GB) | time (h) | workers | CPU (h) | memory (GB) | time (h) | workers | CPU (h) | memory (GB) | time (h) |
| DART | 8 | 6.21 | 2.5 | 0.90 | 8 | 8.89 | 6.3 | 1.36 | 8 | 1.17 | 1.3 | 0.20 |
| Wilcoxon | 1 | 0.02 | 3.3 | 0.02 | 1 | 0.04 | 10.7 | 0.04 | 1 | <0.01 | 1.3 | <0.01 |
| COSG | 1 | <0.01 | 1.7 | <0.01 | 1 | <0.01 | 6.3 | <0.01 | 1 | <0.01 | 0.8 | <0.01 |
| NS-Forest | 8 | 0.39 | 2.9 | 0.09 | 8 | 1.77 | 9.3 | 0.25 | 8 | 0.06 | 1.3 | 0.03 |
| Spapros | 1 | 1.05 | 19.5 | 0.46 | 1 | 0.85 | 24.7 | 0.42 | 1 | 0.34 | 16.3 | 0.19 |

### S5. Labelling cells with DART's rules (rule voting)

Rule voting labels a target's cells with a DART panel's own rules instead of a classifier: the `rules` method of `classify`, `mode = "rules"` in stage 08, or `xspear classify --method rules`. This study did not use it: every prediction reported here comes from the classifier, and we make no claim about rule voting's accuracy.

A rule fires in a cell when all its conditions hold on the cell's log-normalised expression. The reference and the target are normalised over the genes they share, the scale on which the thresholds were learned. Each cell is assigned the cell type with the most firing rules. With up to $K$ mostly single-gene rules per type, ties are common, and a tie goes to the tied type with the highest probability under the classifier trained on the reference; that is why a classifier is trained in this mode too (without one, the first tied type is taken). A cell in which no rule fires is left unassigned. Besides macro-$F_1$ over the real cell types, with an unassigned cell counted as missed for its true type, the evaluation then reports the fraction of cells assigned and macro-$F_1$ among the assigned cells.

Two options change the thresholds before voting. Quantile thresholds (rule scoring, Methods) express each threshold as a quantile of the reference, so that a rule fires at the same rank of the target's distribution as of the reference's. Otsu calibration replaces each threshold with Otsu's threshold (Otsu, 1979) for that gene in the target, which needs no labels.

### S6. Xenium priors in X-SPEAR

**The shipped ovarian cancer prior.** X-SPEAR ships this study's ovarian cancer prior, measured on 10x Genomics' FFPE Flex reference against the Xenium Prime 5K run of the same tissue block: 4,766 genes, median $p_{\mathrm{keep}}$ 0.590, with each gene's two detection rates. It is the prior used here, unchanged. `xspear.xenium_prior("ovarian")` returns it, `xspear prior ovarian -o FILE` writes a copy, `--p-keep ovarian` uses it wherever the command line takes a prior, and it is stage 06's default. It suits a planned human FFPE Xenium Prime 5K experiment (Section 2.5).

**Other sources and gene matching.** A prior can also be a table of genes and $p_{\mathrm{keep}}$ (tables of $1 - p_{\mathrm{keep}}$ from earlier versions are converted on reading), or it can be measured from a paired single-cell reference and Xenium run by Eq. 2, over the genes they share. X-SPEAR matches a prior's genes to the reference's by exact symbol and leaves a gene the prior does not cover unthinned. Stage 06 stops when a prior covers none of the reference's genes and warns when it covers fewer than half. After thinning it recomputes stage 03's normalisation on the thinned copy.

**Depth normalisation** (`xspear profile`; stage 06) is meant for a prior whose single-cell reference was sequenced much more or less deeply than usual, which shifts its single-cell detection rates, and therefore its $p_{\text{keep}}$, as a whole. It keeps the prior's ranking of genes and replaces each value with the value at the same percentile of an anchor profile, by default the shipped ovarian prior. Several priors are combined through the mean, median or 20% trimmed mean of their percentile ranks; a prior with a median $p_{\text{keep}}$ above 0.95 is left out as showing no detection loss; and a gene can be required to be covered by a minimum number of priors. It is off by default: this study used every prior at its own magnitude and did not evaluate depth normalisation.

**Choosing $K$** (`xspear kcurve`). A stratified 25% of the reference is held out and thinned with the prior. Panels are designed on the remaining cells for $K = 1, \ldots, K_{\text{max}}$ (one candidate pool, reselected), a classifier trained on the unthinned remaining cells predicts the held-out cells, and macro-$F_1$ is reported against $K$ and panel size, to help choose $K$ within a gene budget. Thinned single-cell data are cleaner than Xenium, so the values are optimistic. This study did not use it, and Supplementary S1 found that a held-out thinned slice of the reference could not rank selection constants within their plateau.

#### S7. Other package functions

**Off-target dominance factor.** Stage 11 can also score each marker that has predicted off-target genes: the highest mean expression of any of its off-target genes in a cell type other than the one the marker marks, divided by the marker's mean expression in the type it marks. A value of 1 or more is flagged, and the dotplots are then ordered by the score. It is off by default and was not used for Fig. 12. Panel genes chosen for a reason other than marking a cell type can be labelled through named gene sets, and the probe report can be split into specific and unspecific probes and annotated without loading any expression data.

**Forest-based evaluation.** Instead of scoring stage 08's labels, stage 09 can run Spapros' forest-based classification on the panel's genes in the reference. It reports each gene's importance for each cell type and whether that type is where the gene is most expressed, and can keep only the genes above an importance threshold, with dotplots of the genes that do and do not match.

**Supporting code.** A synthetic reference and target, with marker genes for each cell type on a shared background and a target thinned by a known per-gene keep probability, test an installation end to end in seconds. The Spapros worker records the versions it ran with, and its selection, which also yields the reference probe sets, is cached by panel size. Two presets record whether a run used a Xenium prior and map the case names of earlier runs onto them, and a function working in the units of earlier versions ($1 - p_{\text{keep}}$) reproduces their depth-normalised files exactly.
