## Supplementary material for "X-SPEAR: An Extensible Framework for Spatial Panel Evaluation and Refinement": Latex source: compute_table.tex

\begin{tabular}{@{}lcccccccccccc@{}}
\toprule
 & \multicolumn{4}{c}{Ovarian cancer} & \multicolumn{4}{c}{Mouse brain} & \multicolumn{4}{c}{Cervical cancer} \\
\cmidrule(lr){2-5}\cmidrule(lr){6-9}\cmidrule(lr){10-13}
Method & workers & CPU (h) & memory (GB) & time (h) & workers & CPU (h) & memory (GB) & time (h) & workers & CPU (h) & memory (GB) & time (h) \\
\midrule
\dart{} & 8 & 6.21 & 2.5 & 0.90 & 8 & 8.89 & 6.3 & 1.36 & 8 & 1.17 & 1.3 & 0.20 \\
Wilcoxon & 1 & 0.02 & 3.3 & 0.02 & 1 & 0.04 & 10.7 & 0.04 & 1 & $<$0.01 & 1.3 & $<$0.01 \\
COSG & 1 & $<$0.01 & 1.7 & $<$0.01 & 1 & $<$0.01 & 6.3 & $<$0.01 & 1 & $<$0.01 & 0.8 & $<$0.01 \\
NS-Forest & 8 & 0.39 & 2.9 & 0.09 & 8 & 1.77 & 9.3 & 0.25 & 8 & 0.06 & 1.3 & 0.03 \\
\spapros{} & 1 & 1.05 & 19.5 & 0.46 & 1 & 0.85 & 24.7 & 0.42 & 1 & 0.34 & 16.3 & 0.19 \\
\bottomrule
\end{tabular}
