## Supplementary material for "X-SPEAR: An Extensible Framework for Spatial Panel Evaluation and Refinement": Latex source: fig_benchmark.pdf

**A** One panel per method near DART's default size on each target, ranked by mean macro-F1

| # | Method | Genes (per target) | Macro-F1 ovarian | Macro-F1 mouse brain | Macro-F1 cervical | Macro-F1 mean | DART − method | CPU (core-h) | Memory (GB) | Time (h) |
|---|---|---|---|---|---|---|---|---|---|---|
| 1 | DART | 58 / 53 / 23 | **0.555** | **0.680** | **0.595** | **0.610** | — | 5.42 | **6.3** | 0.82 |
| 2 | Spapros | 78 / 62 / 29 | 0.537 | 0.615 | 0.544 | 0.565 | +0.060 | 0.75 | 24.7 | 0.36 |
| 3 | COSG | 56 / 48 / 24 | 0.503 | 0.628 | 0.542 | 0.558 | +0.052 | **<0.01** | 6.3 | **<0.01** |
| 4 | Wilcoxon | 51 / 48 / 24 | 0.464 | 0.502 | 0.560 | 0.509 | +0.099 | 0.02 | 10.7 | 0.02 |
| 5 | NS-Forest | 24 / 22 / 7 | 0.374 | 0.456 | 0.338 | 0.389 | +0.150 | 0.74 | 9.3 | 0.12 |

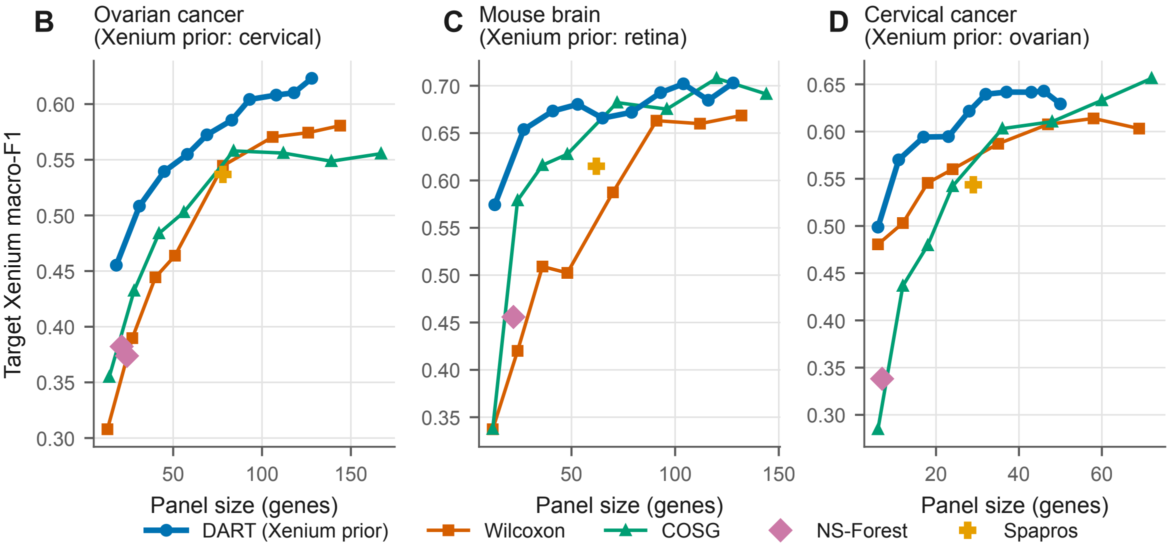
