## Supplementary material for "X-SPEAR: An Extensible Framework for Spatial Panel Evaluation and Refinement": Latex source: fig_pertype.pdf

**A**

Ovarian cancer (Xenium prior for DART: cervical)

| | Tumor Cells (52,609) | Smooth Muscle Cells (31,262) | Proliferative Tumor Cells (24,853) | Tumor Associated Fibroblasts (19,609) | Macrophages (14,017) | Stromal Associated Fibroblasts (12,208) | Endothelial Cells (11,064) | T & NK Cells (5,187) | VEGFA+ Tumor Cells (4,546) | Pericytes (2,478) | Fallopian Tube Epithelium (1,528) | Malignant Cells Lining Cyst (1,347) | Inflammatory Tumor Cells (1,119) | Ciliated Epithelial Cells (936) |
|---|---|---|---|---|---|---|---|---|---|---|---|---|---|---|
| DART (58 genes) | **0.55** | 0.88 | 0.51 | 0.75 | **0.72** | 0.73 | **0.72** | **0.50** | 0.17 | 0.23 | 0.26 | **0.69** | **0.59** | 0.47 |
| Wilcoxon (51 genes) | 0.41 | 0.45 | 0.66 | 0.72 | 0.58 | 0.71 | 0.59 | 0.21 | 0.28 | 0.17 | 0.31 | 0.53 | 0.41 | 0.46 |
| COSG (56 genes) | 0.23 | **0.89** | 0.41 | 0.76 | 0.62 | 0.73 | 0.61 | 0.23 | 0.14 | **0.27** | 0.38 | 0.68 | 0.53 | **0.57** |
| NS-Forest (24 genes) | 0.28 | 0.75 | 0.51 | 0.56 | 0.26 | 0.48 | 0.35 | 0.14 | 0.18 | 0.25 | 0.12 | 0.39 | 0.57 | 0.39 |
| Spapros (78 genes) | 0.49 | 0.77 | **0.68** | **0.81** | 0.66 | **0.77** | 0.68 | 0.27 | **0.31** | 0.16 | **0.39** | 0.55 | 0.49 | 0.47 |

**B**

Mouse brain (Xenium prior for DART: retina)

| | neuron (9,814) | inhibitory interneuron (5,954) | astrocyte (4,626) | oligodendrocyte (3,535) | endothelial cell (2,311) | medium spiny neuron (1,600) | vascular leptomeningeal cell (1,118) | pericyte (1,104) | microglial cell (929) | oligodendrocyte precursor cell (718) | macrophage (48) | T cell (36) |
|---|---|---|---|---|---|---|---|---|---|---|---|---|
| DART (53 genes) | 0.86 | **0.58** | **0.84** | **0.86** | **0.81** | **0.87** | 0.72 | 0.61 | 0.82 | 0.81 | 0.26 | **0.13** |
| Wilcoxon (48 genes) | 0.24 | 0.53 | 0.77 | 0.50 | 0.78 | 0.66 | 0.58 | 0.58 | 0.45 | **0.82** | 0.10 | 0.03 |
| COSG (48 genes) | 0.79 | 0.45 | 0.47 | 0.80 | 0.80 | 0.75 | **0.76** | **0.66** | **0.88** | 0.74 | **0.38** | 0.04 |
| NS-Forest (22 genes) | 0.63 | 0.10 | 0.15 | 0.34 | 0.58 | 0.76 | 0.58 | 0.55 | 0.78 | 0.61 | 0.37 | 0.00 |
| Spapros (62 genes) | **0.87** | 0.35 | 0.82 | 0.79 | 0.77 | 0.83 | 0.65 | 0.47 | 0.85 | 0.80 | 0.12 | 0.05 |

**C**

Cervical cancer (Xenium prior for DART: ovarian)

| | T & NK cells (51,942) | Stromal fibroblasts (37,774) | Squamous cervical epithelium (26,616) | Macrophages (26,107) | Endothelial cells (21,479) | Squamous Carcinoma (17,382) |
|---|---|---|---|---|---|---|
| DART (23 genes) | 0.69 | 0.62 | 0.66 | **0.64** | 0.68 | 0.28 |
| Wilcoxon (24 genes) | 0.68 | **0.64** | **0.69** | 0.29 | 0.68 | 0.38 |
| COSG (24 genes) | 0.63 | 0.64 | 0.58 | 0.30 | **0.72** | 0.38 |
| NS-Forest (7 genes) | 0.53 | 0.56 | 0.19 | 0.09 | 0.24 | **0.43** |
| Spapros (29 genes) | **0.69** | 0.64 | 0.69 | 0.27 | 0.69 | 0.29 |
