## Supplementary material for "X-SPEAR: An Extensible Framework for Spatial Panel Evaluation and Refinement": Latex source: fig_transfer.pdf

| | target's own (upper bound) | ovarian (FFPE, human) | cervical (FFPE, human) | breast (FFPE, human) | retina (FF, mouse) | mouse brain (FF, mouse) | no prior |
|---|---|---|---|---|---|---|---|
| Ovarian cancer (FFPE, human; 45–83 genes) | 0.600<br>+0.129 | same as own | **0.563**<br>+0.093 | 0.520<br>+0.049 | 0.495<br>+0.025 | 0.482<br>+0.011 | 0.470 |
| Mouse brain (FF, mouse; 41–79 genes) | 0.700<br>+0.057 | 0.596<br>-0.047 | 0.529<br>-0.114 | 0.509<br>-0.134 | **0.673**<br>+0.030 | same as own | 0.643 |
| Cervical cancer (FFPE, human; 17–32 genes) | 0.666<br>+0.149 | **0.612**<br>+0.096 | same as own | 0.538<br>+0.022 | 0.500<br>-0.016 | 0.483<br>-0.033 | 0.517 |

Xenium prior used to thin the single-cell reference
