## Supplementary material for "X-SPEAR: An Extensible Framework for Spatial Panel Evaluation and Refinement": Latex source: references.bib

@article{wilcoxon1945,
  author  = {Wilcoxon, Frank},
  title   = {Individual Comparisons by Ranking Methods},
  journal = {Biometrics Bulletin},
  year    = {1945},
  volume  = {1},
  number  = {6},
  pages   = {80--83},
  doi     = {10.2307/3001968}
}

@article{dai2022cosg,
  author  = {Dai, Min and Pei, Xiaobing and Wang, Xiu-Jie},
  title   = {Accurate and Fast Cell Marker Gene Identification with {COSG}},
  journal = {Briefings in Bioinformatics},
  year    = {2022},
  volume  = {23},
  number  = {2},
  pages   = {bbab579},
  doi     = {10.1093/bib/bbab579}
}

@article{aevermann2021nsforest,
  author  = {Aevermann, Brian and Zhang, Yun and Novotny, Mark and Keshk, Mohamed and Bakken, Trygve and Miller, Jeremy and Hodge, Rebecca and Lelieveldt, Boudewijn and Lein, Ed and Scheuermann, Richard H.},
  title   = {A Machine Learning Method for the Discovery of Minimum Marker Gene Combinations for Cell Type Identification from Single-Cell {RNA} Sequencing},
  journal = {Genome Research},
  year    = {2021},
  volume  = {31},
  number  = {10},
  pages   = {1767--1780},
  doi     = {10.1101/gr.275569.121}
}

@article{kuemmerle2024spapros,
  author  = {Kuemmerle, Louis B. and Luecken, Malte D. and Firsova, Alexandra B. and Barros de Andrade e Sousa, Lisa and Stra{\ss}er, Lena and Mekki, Ilhem Isra and Campi, Francesco and Heumos, Lukas and Shulman, Maiia and Beliaeva, Valentina and Hediyeh-Zadeh, Soroor and Schaar, Anna C. and Mahbubani, Krishnaa T. and Sountoulidis, Alexandros and Balassa, Tam{\'a}s and Kovacs, Ferenc and Horvath, Peter and Piraud, Marie and Ert{\"u}rk, Ali and Samakovlis, Christos and Theis, Fabian J.},
  title   = {Probe Set Selection for Targeted Spatial Transcriptomics},
  journal = {Nature Methods},
  year    = {2024},
  volume  = {21},
  number  = {12},
  pages   = {2260--2270},
  doi     = {10.1038/s41592-024-02496-z}
}

@article{janesick2023xenium,
  author  = {Janesick, Amanda and Shelansky, Robert and Gottscho, Andrew D. and others},
  title   = {High Resolution Mapping of the Tumor Microenvironment Using Integrated Single-Cell, Spatial and In Situ Analysis},
  journal = {Nature Communications},
  year    = {2023},
  volume  = {14},
  pages   = {8353},
  doi     = {10.1038/s41467-023-43458-x}
}

@article{wolf2018scanpy,
  author  = {Wolf, F. Alexander and Angerer, Philipp and Theis, Fabian J.},
  title   = {{SCANPY}: Large-Scale Single-Cell Gene Expression Data Analysis},
  journal = {Genome Biology},
  year    = {2018},
  volume  = {19},
  pages   = {15},
  doi     = {10.1186/s13059-017-1382-0}
}

@article{friedman2008rulefit,
  author  = {Friedman, Jerome H. and Popescu, Bogdan E.},
  title   = {Predictive Learning via Rule Ensembles},
  journal = {The Annals of Applied Statistics},
  year    = {2008},
  volume  = {2},
  number  = {3},
  pages   = {916--954},
  doi     = {10.1214/07-AOAS148}
}

@article{pedregosa2011sklearn,
  author  = {Pedregosa, Fabian and Varoquaux, Ga{\"e}l and Gramfort, Alexandre and others},
  title   = {Scikit-learn: Machine Learning in {Python}},
  journal = {Journal of Machine Learning Research},
  year    = {2011},
  volume  = {12},
  pages   = {2825--2830}
}

@article{allen2023brainaging,
  author  = {Allen, William E. and Blosser, Timothy R. and Sullivan, Zuri A. and Dulac, Catherine and Zhuang, Xiaowei},
  title   = {Molecular and Spatial Signatures of Mouse Brain Aging at Single-Cell Resolution},
  journal = {Cell},
  year    = {2023},
  volume  = {186},
  number  = {1},
  pages   = {194--208},
  doi     = {10.1016/j.cell.2022.12.010}
}

@article{tran2019rgc,
  author  = {Tran, Nicholas M. and Shekhar, Karthik and Whitney, Irene E. and others},
  title   = {Single-Cell Profiles of Retinal Ganglion Cells Differing in Resilience to Injury Reveal Neuroprotective Genes},
  journal = {Neuron},
  year    = {2019},
  volume  = {104},
  number  = {6},
  pages   = {1039--1055},
  doi     = {10.1016/j.neuron.2019.11.006}
}

@article{sanes2015rgc,
  author  = {Sanes, Joshua R. and Masland, Richard H.},
  title   = {The Types of Retinal Ganglion Cells: Current Status and Implications for Neuronal Classification},
  journal = {Annual Review of Neuroscience},
  year    = {2015},
  volume  = {38},
  pages   = {221--246},
  doi     = {10.1146/annurev-neuro-071714-034120}
}

@article{xu2021scanvi,
  author  = {Xu, Chenling and Lopez, Romain and Mehlman, Edouard and Regier, Jeffrey and Jordan, Michael I. and Yosef, Nir},
  title   = {Probabilistic Harmonization and Annotation of Single-Cell Transcriptomics Data with Deep Generative Models},
  journal = {Molecular Systems Biology},
  year    = {2021},
  volume  = {17},
  number  = {1},
  pages   = {e9620},
  doi     = {10.15252/msb.20209620}
}

@article{dominguezconde2022celltypist,
  author  = {Dom{\'i}nguez Conde, C. and Xu, C. and Jarvis, L. B. and others},
  title   = {Cross-Tissue Immune Cell Analysis Reveals Tissue-Specific Features in Humans},
  journal = {Science},
  year    = {2022},
  volume  = {376},
  number  = {6594},
  pages   = {eabl5197},
  doi     = {10.1126/science.abl5197}
}

@article{aran2019singler,
  author  = {Aran, Dvir and Looney, Agnieszka P. and Liu, Leqian and others},
  title   = {Reference-Based Analysis of Lung Single-Cell Sequencing Reveals a Transitional Profibrotic Macrophage},
  journal = {Nature Immunology},
  year    = {2019},
  volume  = {20},
  pages   = {163--172},
  doi     = {10.1038/s41590-018-0276-y}
}

@article{zhang2019cellassign,
  author  = {Zhang, Allen W. and O'Flanagan, Ciara and Chavez, Elizabeth A. and others},
  title   = {Probabilistic Cell-Type Assignment of Single-Cell {RNA}-seq for Tumor Microenvironment Profiling},
  journal = {Nature Methods},
  year    = {2019},
  volume  = {16},
  pages   = {1007--1015},
  doi     = {10.1038/s41592-019-0529-1}
}

@article{yuan2006grouplasso,
  author  = {Yuan, Ming and Lin, Yi},
  title   = {Model Selection and Estimation in Regression with Grouped Variables},
  journal = {Journal of the Royal Statistical Society: Series B},
  year    = {2006},
  volume  = {68},
  number  = {1},
  pages   = {49--67},
  doi     = {10.1111/j.1467-9868.2005.00532.x}
}

@misc{li2026retinaatlas,
  author       = {Li, J. and Wang, J. and Ibarra, I. L. and Cheng, X. and Luecken, M. D. and others},
  title        = {Single Cell Atlas of the Human Retina},
  howpublished = {CZ CELLxGENE collection 4c6eaf5c-6d57-4c76-b1e9-60df8c655f1e; Nature Genetics},
  year         = {2026},
  doi          = {10.1038/s41588-025-02454-1}
}

@article{lyu2025retinaaging,
  author  = {Lyu, P. and Palazzo, I. and Jin, Y. and Campbell, L. J. and Santiago, C. P. and others},
  title   = {Comparative Single-Cell Multiomic Analysis Reveals Evolutionarily Conserved and Species-Specific Cellular Mechanisms Mediating Natural Retinal Aging},
  journal = {bioRxiv},
  year    = {2025},
  doi     = {10.1101/2025.09.09.673285}
}

@article{wolock2019scrublet,
  author  = {Wolock, Samuel L. and Lopez, Romain and Klein, Allon M.},
  title   = {Scrublet: computational identification of cell doublets in single-cell transcriptomic data},
  journal = {Cell Systems},
  volume  = {8},
  number  = {4},
  pages   = {281--291.e9},
  year    = {2019},
  doi     = {10.1016/j.cels.2018.11.005}
}

@article{korsunsky2019harmony,
  author  = {Korsunsky, Ilya and Millard, Nghia and Fan, Jean and Slowikowski, Kamil and Zhang, Fan and Wei, Kevin and Baglaenko, Yuriy and Brenner, Michael and Loh, Po-Ru and Raychaudhuri, Soumya},
  title   = {Fast, sensitive and accurate integration of single-cell data with {Harmony}},
  journal = {Nature Methods},
  volume  = {16},
  pages   = {1289--1296},
  year    = {2019},
  doi     = {10.1038/s41592-019-0619-0}
}

@article{mcinnes2018umap,
  author  = {McInnes, Leland and Healy, John and Melville, James},
  title   = {{UMAP}: Uniform Manifold Approximation and Projection for Dimension Reduction},
  journal = {arXiv preprint arXiv:1802.03426},
  year    = {2018}
}

@article{otsu1979threshold,
  author  = {Otsu, Nobuyuki},
  title   = {A threshold selection method from gray-level histograms},
  journal = {IEEE Transactions on Systems, Man, and Cybernetics},
  volume  = {9},
  number  = {1},
  pages   = {62--66},
  year    = {1979},
  doi     = {10.1109/TSMC.1979.4310076}
}

@article{singh2021imodels,
  author  = {Singh, Chandan and Nasseri, Keyan and Tan, Yan Shuo and Tang, Tiffany and Yu, Bin},
  title   = {imodels: a python package for fitting interpretable models},
  journal = {Journal of Open Source Software},
  volume  = {6},
  number  = {61},
  pages   = {3192},
  year    = {2021},
  doi     = {10.21105/joss.03192}
}
