## Supplementary material for "X-SPEAR: An Extensible Framework for Spatial Panel Evaluation and Refinement": Latex source: XSPEAR.tex

% ==============================================================================
% X-SPEAR: eXtensible Spatial Panel Evaluation And Refinement
% bioRxiv preprint. Restructured 2026-10-08 (Rushil's list: datasets table in main, Xenium-prior
% comparison, benchmark with Spapros and a ranked summary table, per-cell-type and K/O figures,
% single-cell RGC section, off-target dotplot example; no ablation, no RGC application).
% Every number traces to analyses/staged_pipeline/RESULTS.md or a committed result file; figure
% scripts are in figures/. Build: ./build_biorxiv.sh
% ==============================================================================
\documentclass[11pt,letterpaper]{article}

% --- Core packages ---
\usepackage[utf8]{inputenc}
\usepackage[T1]{fontenc}
\usepackage{mathpazo}
\usepackage{microtype}
\usepackage[margin=1in]{geometry}

% --- Math and symbols ---
\usepackage{amsmath,amssymb,amsfonts}

% --- Graphics and tables ---
\usepackage{graphicx}
\usepackage{booktabs}
\usepackage{multirow}
\usepackage{array}
\usepackage{caption}
\captionsetup{font=small,labelfont=bf}
\usepackage[section]{placeins}   % floats stay inside the section that cites them

% --- Colors and hyperlinks ---
\usepackage[dvipsnames]{xcolor}
\usepackage[colorlinks=true,linkcolor=MidnightBlue,citecolor=MidnightBlue,urlcolor=MidnightBlue]{hyperref}
\usepackage{xurl}                 % long data URLs may break at any character

% --- Bibliography ---
\usepackage[round,sort&compress]{natbib}

% --- Spacing ---
\usepackage{setspace}
\onehalfspacing
\raggedbottom

% --- Formatting helpers ---
\usepackage[normalem]{ulem}
\usepackage{enumitem}
\setlist{itemsep=1pt,topsep=3pt,parsep=0pt}
\emergencystretch=1.5em

% --- Schematic diagrams ---
\usepackage{tikz}
\usetikzlibrary{positioning, arrows.meta, calc, fit, backgrounds}

% --- Section formatting ---
\usepackage{titlesec}
\titleformat{\section}{\large\bfseries}{\thesection}{1em}{}
\titleformat{\subsection}{\normalsize\bfseries}{\thesubsection}{1em}{}
\titleformat{\subsubsection}{\normalsize\itshape}{\thesubsubsection}{1em}{}

% --- Custom commands ---
\newcommand{\xspear}{\textsc{X-SPEAR}}
\newcommand{\atpf}{\textsc{ATPF}}
\newcommand{\rulefit}{\textsc{RuleFit}}
\newcommand{\spapros}{Spapros}
\newcommand{\dart}{\textsc{DART}}
\newcommand{\Fone}{macro-$F_1$}
\newcommand{\pkeep}{{p_{\mathrm{keep}}}}
% Items only the authors can supply. The build script labels the PDF _DRAFT while any remain.
\newcommand{\PENDING}[1]{\textcolor{red!80!black}{\textbf{[#1]}}}
% Numbers still being computed while the manuscript is restructured (the build refuses any left).
\newcommand{\TBD}[1]{\textcolor{orange!80!black}{\textbf{[TBD: #1]}}}
% Author names in order; used by the supplement's title block and the build's PDF metadata.
\newcommand{\authornames}{Rushil Patel, Bryant Duong, Veer Patel, Rachel Lyn-Sue, Kyungbin Oh, Yuyan Cheng}

% ==============================================================================
\begin{document}

\title{\xspear{}: An Extensible Framework for Spatial Panel Evaluation and Refinement\\[0.4em]
  \large Detection-aware combinatorial marker selection for targeted spatial transcriptomics}

\author{%
  Rushil Patel\textsuperscript{1}, Bryant Duong\textsuperscript{1}, Veer Patel\textsuperscript{1},
  Rachel Lyn-Sue\textsuperscript{1},\\ Kyungbin Oh\textsuperscript{1},
  Yuyan Cheng\textsuperscript{1,*}\\[0.6em]
  \small\textsuperscript{1}Department of Ophthalmology, Scheie Eye Institute, Perelman School of Medicine,\\
  \small University of Pennsylvania, Philadelphia, PA, USA\\

\date{}
\maketitle

% ==============================================================================
\begin{abstract}
\noindent
Targeted spatial transcriptomics platforms such as 10x Genomics Xenium measure a predefined panel of
genes. A gene is usually detected in fewer cells in the spatial assay than in the single-cell RNA
sequencing (scRNA-seq) reference used to choose it, by an amount that differs from gene to gene. We
present eXtensible Spatial Panel Evaluation And Refinement (\xspear{}), a framework for designing and
checking Xenium gene panels. Its marker-selection method, Decision-rule Aggregation for Rare-cell
Typing (\dart{}), learns short Boolean gene rules and keeps several gene-disjoint rules per cell type.
\dart{} first thins the reference to per-gene detection rates measured in a previous experiment, which
we call a Xenium prior, so no Xenium data from the target sample are needed before the panel is
ordered. A Xenium prior helped most when it matched the target's assay chemistry and species, even
when it came from a different tissue, and a mismatched prior could lower accuracy. With such a prior,
\dart{} outperformed four marker-selection methods on held-out Xenium data from ovarian cancer, mouse
brain and cervical cancer at matched panel size: by 0.04--0.10 macro-averaged $F_1$ score (\Fone{})
over the Wilcoxon rank-sum test, 0.04--0.08 over COSine similarity-based marker Gene identification
(COSG), 0.10--0.17 over Necessary and Sufficient Forest (NS-Forest) and 0.04--0.08 over the
probe-set selection method \spapros{}. It reached the Wilcoxon test's best accuracy with 42--69\%
fewer genes. \dart{}'s rules agree with the
classifier that uses the panel and name recognisable marker programs for each cell type. \xspear{}
also checks the probes of any panel for predicted off-target transcripts that are expressed in other
cell types. \xspear{} is available as an open-source Python package.
\end{abstract}

\vspace{0.5em}
\noindent\textbf{Keywords:} spatial transcriptomics, Xenium, gene panel design, marker selection,
Boolean rules, detection efficiency, single-cell RNA sequencing

% ==============================================================================
\section{Introduction}
\label{sec:introduction}

Targeted in situ transcriptomics platforms such as 10x Genomics Xenium measure hundreds to thousands
of genes at subcellular resolution in intact tissue, so that cell types, subtypes and states can be
identified in their spatial context~\citep{janesick2023xenium}. They require the user to choose the
gene panel before the experiment. The panel is usually chosen from an annotated single-cell RNA
sequencing (scRNA-seq) or single-nucleus RNA sequencing (snRNA-seq) reference, and the choice
determines which cell identities the experiment can resolve. A probe that hybridises to unintended
transcripts can also produce misleading spatial signal, so the designed probes need checking.

Several computational methods select markers or probe sets for this purpose: differential-expression
rankings such as the Wilcoxon rank-sum test~\citep{wilcoxon1945}; specificity scores such as COSine
similarity-based marker Gene identification (COSG)~\citep{dai2022cosg}; minimal marker combinations
such as Necessary and Sufficient Forest (NS-Forest)~\citep{aevermann2021nsforest}; and probe-set
optimisation such as \spapros{}~\citep{kuemmerle2024spapros}. Two problems remain. First, single-gene
markers are often insufficient for rare populations, closely related subtypes and transient states,
and methods that find gene combinations usually reduce each cell type to one combination, which fails
if any of its genes is poorly detected. Second, a gene's detection in a targeted spatial assay
differs from its detection in the single-cell reference. Across the ovarian cancer panel studied
here, the median gene is detected in Xenium in only about half as many cells as in the matched
reference, and the loss varies strongly from gene to gene. A marker that looks excellent in the
reference may be nearly invisible in situ. The methods we compare choose genes from the reference as
measured and do not model this loss.

Here we present eXtensible Spatial Panel Evaluation And Refinement (\xspear{}), a framework with two
modules. The first is Decision-rule Aggregation for Rare-cell Typing (\dart{}), which learns short
Boolean gene rules and selects several gene-disjoint rules for each cell type. Before learning,
\dart{} thins the reference to each gene's measured single-cell-to-Xenium detection rate, so that
rules relying on poorly detected genes score poorly and are not chosen. The detection rates come from
a \emph{Xenium prior}: a previous experiment in which a single-cell reference and a Xenium run were
both measured, usually on a different tissue, because a designer never has Xenium data from the
target sample before ordering its panel. The second module checks the probes of any panel, whichever
method chose it, for predicted off-target transcripts that are expressed in other cell types. The
current version targets cell-type marker selection for Xenium. The ``X'' denotes extensibility,
toward other targeted platforms and other selection objectives.

We first ask which Xenium priors help \dart{}, using every combination of prior and target among five
paired single-cell and Xenium datasets. We then benchmark \dart{} against the Wilcoxon test, COSG,
NS-Forest and \spapros{} on three targets (ovarian cancer, mouse brain and cervical cancer), with a
deployment rule throughout: nothing measured on the target's Xenium informs the panel, and its labels
are used only for scoring. With a Xenium prior from a different tissue, \dart{} outperformed all four
methods at matched panel size on every target. Its rules agree with the downstream classifier and name
interpretable marker programs. On 45 mouse retinal ganglion cell (RGC) subtypes, held out within a
single-cell atlas, \dart{} matched the best methods and led COSG and NS-Forest at every subtype
abundance.

% ==============================================================================
\section{Results}
\label{sec:results}

% ------------------------------------------------------------------------------
\subsection{\xspear{} designs a detection-aware panel and checks its probes}
\label{sec:results-framework}

\xspear{} takes an annotated single-cell reference of the tissue to be profiled, together with a
Xenium prior, and returns a Xenium gene panel with evidence for classification performance and
probe-level off-target risk (Fig.~\ref{fig:fig1}). It is a Python package that can be used as a library, from the
command line or as a pipeline of twelve stages (below). Panel design with \dart{} proceeds in three steps
(Fig.~\ref{fig:dart-schematic}).

\begin{figure}[!htbp]
  \centering
  \resizebox{\textwidth}{!}{%
  \begin{tikzpicture}[
    font=\footnotesize,
    every node/.append style={execute at begin node={\hyphenpenalty=10000\exhyphenpenalty=10000}},
    arr/.style={-{Stealth[length=2.6mm,width=2mm]}, thick, draw=gray!70},
    io/.style={draw, rounded corners=2pt, fill=blue!8, align=center, text width=30mm, minimum height=9mm},
    pri/.style={draw=orange!75, dashed, rounded corners=2pt, fill=orange!10, align=center, text width=30mm, minimum height=9mm},
    mod/.style={draw=orange!70, rounded corners=2pt, fill=orange!15, align=center, text width=27mm, minimum height=11mm},
    outbox/.style={draw, rounded corners=2pt, fill=green!11, align=center, text width=24mm, minimum height=11mm},
    rule/.style={draw, rounded corners=2pt, fill=gray!7, align=center, inner sep=3pt, font=\scriptsize},
    frame/.style={draw, rounded corners=3pt},
    hdr/.style={font=\bfseries, anchor=base}
  ]
    \draw[frame] (0,0) rectangle (19.6,-3.2);
    \node[hdr] at (9.8,0.25) {A\quad \xspear{} workflow};
    \node[io]  (ref)  at (2.0,-0.95) {annotated single-cell\\reference};
    \node[pri] (pri)  at (2.0,-2.3)  {Xenium prior\\(previous experiment)};
    \node[mod] (dart) at (6.7,-1.6)  {\textbf{\dart{}}\\marker selection};
    \node[mod] (probe)at (10.4,-1.6) {\textbf{Xenium Panel}\\\textbf{Designer}\\probe design};
    \node[mod] (offt) at (14.1,-1.6) {\textbf{off-target}\\check};
    \node[outbox] (panel) at (17.6,-1.6) {Xenium\\gene panel};
    \draw[arr] (ref.east)--([yshift=2.5mm]dart.west);
    \draw[arr, draw=orange!75] (pri.east)--([yshift=-2.5mm]dart.west);
    \foreach \a/\b in {dart/probe,probe/offt,offt/panel} \draw[arr] (\a.east)--(\b.west);
    \draw[frame] (0,-4.05) rectangle (9.4,-9.1);
    \node[hdr] at (4.7,-3.8) {B\quad \dart{} rule assembly};
    \node[draw=orange!70, rounded corners=2pt, fill=orange!13, align=center, text width=72mm] (thin) at (4.7,-4.75)
      {detection-aware thinning: the reference is thinned\\to the Xenium prior's per-gene detection rates};
    \node[align=center, font=\scriptsize] (ct) at (4.7,-6.05) {per cell type: up to $K$ gene-disjoint rules};
    \node[rule] (r1) at (4.7,-6.55) {Rule 1:~$\mathit{GeneA}{>}t_1 \wedge \mathit{GeneB}{\le}t_2$};
    \node[rule] (r2) at (4.7,-7.15) {Rule 2:~$\mathit{GeneC}{>}t_3$};
    \node[draw, dashed, rounded corners=2pt, inner sep=4pt, fit=(ct)(r1)(r2)] (rules) {};
    \node[draw, rounded corners=2pt, fill=green!11, align=center, text width=52mm, minimum height=7mm] (union) at (4.7,-8.5)
      {panel $=\bigcup$ selected-rule genes};
    \draw[arr] (thin.south)--(rules.north);
    \draw[arr] (rules.south)--(union.north);
    \draw[frame] (10.2,-4.05) rectangle (19.6,-9.1);
    \node[hdr] at (14.9,-3.8) {C\quad off-target check (any method's panel)};
    \node[draw, rounded corners=2pt, fill=blue!8, align=center, text width=34mm, minimum height=11mm] (rep) at (12.5,-5.2)
      {probe report from the\\Xenium Panel Designer};
    \node[draw, rounded corners=2pt, fill=gray!8, align=center, text width=34mm, minimum height=11mm] (dot) at (12.5,-7.6)
      {dotplot: marker and its\\off-target genes by cell type};
    \node[draw, rounded corners=2pt, fill=green!12, align=center, text width=30mm, minimum height=11mm] (ok) at (17.3,-6.4)
      {off-targets silent\\$\Rightarrow$ keep probe};
    \node[draw, rounded corners=2pt, fill=red!9, align=center, text width=30mm, minimum height=11mm] (bad) at (17.3,-8.25)
      {off-targets expressed\\elsewhere $\Rightarrow$ flag probe};
    \draw[arr] (rep.south)--(dot.north);
    \draw[arr, draw=green!50!black] (dot.east)--(ok.west);
    \draw[arr, draw=red!65, dashed] (dot.east)--(bad.west);
  \end{tikzpicture}}
  \caption{\textbf{\xspear{} designs a detection-aware Xenium panel and checks its probes.}
  \textbf{(A)}~Workflow: \dart{} marker selection from an annotated single-cell reference and a
  Xenium prior, probe design in the Xenium Panel Designer, and the off-target check.
  \textbf{(B)}~\dart{} keeps up to $K$ gene-disjoint Boolean rules per cell type, learned on a
  reference thinned to the Xenium prior's per-gene detection rates. A rule can require a gene to be
  high ($>$) or low ($\le$). \textbf{(C)}~The off-target check reads the probe report of any panel,
  draws each marker beside its predicted off-target genes across the reference's cell types, and
  flags probes whose off-target genes are expressed in cell types the marker does not mark.}
  \label{fig:fig1}
\end{figure}

The first step is detection-aware thinning. For each gene, a keep probability $\pkeep$ is the ratio of
its detection rate (the fraction of cells with a nonzero count) in the Xenium prior's Xenium run to its
detection rate in that experiment's single-cell reference (Methods). Each nonzero count of the
single-cell reference is kept with probability $\pkeep$ and zeroed otherwise, so that the thinned
reference reproduces each gene's expected in situ detection. The prior's cells need no labels, and the
target's Xenium is never used.

The second step is rule learning. Three learners propose candidate Boolean rules for each cell type:
thresholded conjunctions such as $\textit{GeneA}>t_1 \wedge \textit{GeneB}\le t_2$, which require one
gene to be high and another to be low. The Asymmetric Tree Panel Finder (\atpf{}), developed here,
grows decision trees whose splits isolate one target cell type rather than separating all classes.
\rulefit{}~\citep{friedman2008rulefit}, a rule-ensemble method, is applied to random-forest and
gradient-boosted ensembles; it extracts rules from the trees and keeps those with nonzero
$L_1$-penalised coefficients.

The third step is orthogonal selection. Candidates are ranked by $F_1$ minus a penalty per additional
gene, and up to $K$ rules are kept per cell type, sharing at most $O_{\max}$ genes. The default is
gene-disjoint rules ($O_{\max}=0$). The panel is the union of the selected rules' genes.

\begin{figure}[!htbp]
  \centering
  \resizebox{\textwidth}{!}{%
  \begin{tikzpicture}[
    font=\small,
    every node/.append style={execute at begin node={\hyphenpenalty=10000\exhyphenpenalty=10000}},
    box/.style    ={draw, rounded corners, align=center, minimum height=12mm, inner sep=5pt, fill=MidnightBlue!7},
    learner/.style={draw, rounded corners, align=center, minimum height=9mm, minimum width=34mm, fill=ForestGreen!10},
    opt/.style    ={draw, rounded corners, align=center, inner sep=5pt, fill=BurntOrange!12},
    outbox/.style ={draw, rounded corners, align=center, minimum height=12mm, inner sep=5pt, fill=MidnightBlue!20},
    ar/.style     ={-{Stealth[length=2.6mm,width=2mm]}, thick},
  ]
    \node[box] (ref) {Annotated single-cell\\ reference\\[1pt] \footnotesize(cells $\times$ genes, cell-type labels)};
    \node[opt, above=8mm of ref, xshift=30mm] (xen) {Xenium prior\\[1pt] \footnotesize(per-gene detection rates of a previous\\ \footnotesize single-cell and Xenium experiment; no labels)};
    \node[box, right=16mm of ref] (thin) {Detection-aware\\ thinning\\[1pt] \footnotesize($\pkeep$, Eq.~\ref{eq:pkeep})};
    \draw[ar] (xen.south) -- (thin.north) node[midway, right, font=\footnotesize] {$\pkeep$};
    \draw[ar] (ref) -- (thin);
    \node[learner, right=18mm of thin, yshift=11mm] (atpf) {\textsc{ATPF}\\[-1pt] \footnotesize target-isolating trees};
    \node[learner, below=3mm of atpf] (rf) {\textsc{RuleFit}--RF};
    \node[learner, below=3mm of rf] (gb) {\textsc{RuleFit}--GB};
    \draw[ar] (thin.east) -- (atpf.west);
    \draw[ar] (thin.east) -- (rf.west);
    \draw[ar] (thin.east) -- (gb.west);
    \node[box, right=18mm of rf] (pool) {Pooled candidate\\ Boolean rules\\[1pt] \footnotesize(per cell type)};
    \draw[ar] (atpf.east) -- ([yshift=2.5mm]pool.west);
    \draw[ar] (rf.east) -- (pool.west);
    \draw[ar] (gb.east) -- ([yshift=-2.5mm]pool.west);
    \node[box, right=14mm of pool] (sel) {Size-penalised ranking\\ $+$ orthogonal\\ multi-rule selection\\[1pt] \footnotesize($K$ rules, $\le O_{\max}$ shared genes)};
    \draw[ar] (pool) -- (sel);
    \node[outbox, right=14mm of sel, minimum width=18mm] (panel) {Gene\\ panel};
    \draw[ar] (sel) -- (panel);
    \begin{scope}[every node/.style={font=\footnotesize, text=gray!45!black}]
      \node[below=1.5mm of thin] {stage 06};
      \node[below=1.5mm of gb] {stage 07a: rule learning};
      \node[below=1.5mm of pool] {stage 07b: rule scoring};
      \node[below=1.5mm of sel] {stages 07c--07d};
    \end{scope}
  \end{tikzpicture}%
  }
  \caption{\textbf{The \dart{} algorithm.} The single-cell reference is first thinned so that each gene
  is detected as often as in the Xenium prior: each nonzero count is kept with probability $\pkeep$
  (Eq.~\ref{eq:pkeep}). The prior is a previous experiment with both a single-cell reference and a
  Xenium run, usually of another tissue, and its labels are not used. Three learners then propose
  candidate Boolean rules per cell type: \atpf{}, whose trees isolate one cell type, and \rulefit{} on
  random-forest (RF) and gradient-boosted (GB) ensembles. Candidates are ranked by $F_1$ with a penalty
  per additional gene, and up to $K$ rules sharing at most $O_{\max}$ genes are kept per type. The
  panel is the union of the selected rules' genes. Grey labels give the pipeline stage that runs each
  step (Fig.~\ref{fig:pipeline}).}
  \label{fig:dart-schematic}
\end{figure}

The off-target module reads the probe report that the Xenium Panel Designer returns for a submitted
panel, which lists each probe's predicted off-target genes. For each marker it draws a dotplot of the
marker beside its off-target genes across the reference's cell types. A probe whose off-target genes
are expressed in cell types that the marker does not mark would add false signal there and should be
replaced; the dotplot shows which probes these are. Because the check needs only the probe report and
the reference, it applies to a panel from any selection method (Section~\ref{sec:results-offtarget}).

\xspear{}'s pipeline runs these steps as twelve numbered stages, each configured by its own file and
each optional (Fig.~\ref{fig:pipeline}). Stages 00--04 load and prepare the single-cell reference:
ingest, quality control, a cap on the cells per cell type, normalisation and, if wanted, an embedding.
Stage 06 thins a copy of the reference to the Xenium prior's detection rates, and stage 07 selects the
panel, with \dart{} (as four substages: rule learning, rule scoring, ranking and orthogonal selection)
or with one of the other selectors compared here. Stage 10 exports the panel together with the
reference bundle that the Xenium Panel Designer takes, and stage 11 checks the probe report it returns.
Where a labelled Xenium run of the target exists, stages 05, 08 and 09 load it, label its cells with a
classifier trained on the reference and score the labels, as the evaluations in this study do. Every
selector writes the same panel file, which is all the later stages read, so the classification,
export and off-target stages serve a panel from any method. The interfaces and each stage are
described in the Methods (Section~\ref{sec:methods-software}).

\begin{figure}[!htbp]
  \centering
  \resizebox{\textwidth}{!}{%
  \begin{tikzpicture}[
    font=\footnotesize,
    every node/.append style={execute at begin node={\hyphenpenalty=10000\exhyphenpenalty=10000}},
    arr/.style={-{Stealth[length=2.4mm,width=1.9mm]}, thick, draw=gray!70},
    io/.style={-{Stealth[length=2.2mm,width=1.8mm]}, semithick, draw=gray!55},
    st/.style={draw, rounded corners=2pt, align=center, text width=25.5mm, minimum height=13.5mm,
               inner sep=2pt},
    prep/.style={st, fill=MidnightBlue!8},
    des/.style={st, fill=orange!17, draw=orange!75},
    ev/.style={st, fill=violet!11},
    ord/.style={st, fill=green!12},
    off/.style={dashed},
    note/.style={font=\scriptsize, align=center, text width=27mm, text=gray!25!black},
    key/.style={draw, rounded corners=1pt, minimum width=3.2mm, minimum height=3.2mm, inner sep=0pt}
  ]
    % ---- the twelve stages, in the order `xspear run` executes them ---------------------------
    \node[prep] (s00) at (0,0)    {\textbf{00 Ingest}\\counts, gene identifiers, metadata};
    \node[prep] (s01) at (3.05,0)  {\textbf{01 Quality control}\\cell and gene filters, doublets};
    \node[prep] (s02) at (6.1,0)  {\textbf{02 Sampling}\\cap cells per cell type};
    \node[prep] (s03) at (9.15,0)  {\textbf{03 Normalisation}\\CP10k, then $\log(1+x)$};
    \node[prep, off] (s04) at (12.2,0) {\textbf{04 Embedding}\\PCA, UMAP, Harmony};
    \node[ev] (s05) at (15.25,0)   {\textbf{05 Spatial ingest}\\target cells and labels};
    \node[des] (s06) at (0,-3.1)  {\textbf{06 Detection profile}\\thin a copy of the reference};
    \node[des] (s07) at (3.05,-3.1) {\textbf{07 Selection}\\\dart{} (07a--d) or another selector};
    \node[ev] (s08) at (6.1,-3.1) {\textbf{08 Classification}\\label the target's cells};
    \node[ev] (s09) at (9.15,-3.1) {\textbf{09 Evaluation}\\macro-$F_1$, per-type $F_1$};
    \node[ord] (s10) at (12.2,-3.1) {\textbf{10 Export}\\panel, upload bundle};
    \node[ord, off] (s11) at (15.25,-3.1) {\textbf{11 Off-target}\\probe report $\to$ dotplots};
    \foreach \a/\b in {s00/s01,s01/s02,s02/s03,s03/s04,s04/s05,s06/s07,s07/s08,s08/s09,s09/s10,s10/s11}
      \draw[arr] (\a.east)--(\b.west);
    \draw[arr] (s05.south) -- ++(0,-0.78) -| (s06.north);

    % ---- what enters and leaves ---------------------------------------------------------------
    \node[note] (iref) at (0,1.75)  {\textbf{annotated single-cell reference}};
    \node[note] (itgt) at (15.25,1.75) {\textbf{target Xenium}, if labelled (optional)};
    \draw[io] (iref)--(s00.north);
    \draw[io] (itgt)--(s05.north);
    \node[note] (ipri) at (0,-4.95)  {\textbf{Xenium prior}\\(per-gene $\pkeep$)};
    \node[note] (opan) at (3.05,-4.95) {panel genes (and \dart{}'s rules)};
    \node[note] (ometr) at (9.15,-4.95) {predictions and scores};
    \node[note] (obun) at (12.2,-4.95) {bundle for the Xenium Panel Designer};
    \node[note] (iprb) at (15.25,-4.95) {its probe report};
    \draw[io, draw=orange!75] (ipri)--(s06.south);
    \draw[io] (s07.south)--(opan);
    \draw[io] (s09.south)--(ometr);
    \draw[io] (s10.south)--(obun);
    \draw[io] (iprb)--(s11.south);

    % ---- key -----------------------------------------------------------------------------------
    \node[key, fill=MidnightBlue!8] (k1) at (-1.15,-6.15) {};
    \node[anchor=west, font=\scriptsize] at (k1.east) {reference preparation};
    \node[key, fill=orange!17, draw=orange!75] (k2) at (2.55,-6.15) {};
    \node[anchor=west, font=\scriptsize] at (k2.east) {panel design};
    \node[key, fill=violet!11] (k3) at (5.15,-6.15) {};
    \node[anchor=west, font=\scriptsize] at (k3.east) {evaluation (labelled target only)};
    \node[key, fill=green!12] (k4) at (9.85,-6.15) {};
    \node[anchor=west, font=\scriptsize] at (k4.east) {export and probe check};
    \node[key, dashed, fill=white] (k5) at (13.75,-6.15) {};
    \node[anchor=west, font=\scriptsize] at (k5.east) {off by default};
  \end{tikzpicture}}
  \caption{\textbf{The \xspear{} pipeline.} Twelve numbered stages run in this order, each configured
  by its own file and each optional. Stages 00--04 prepare the annotated single-cell reference. Stage
  06 thins a copy of it to the Xenium prior's per-gene detection rates, and stage 07 selects the
  panel: \dart{} as four substages (07a rule learning, 07b rule scoring, 07c ranking, 07d orthogonal
  selection) or another selector (the Wilcoxon test, COSG, NS-Forest, \spapros{} or a curated list).
  Stage 10 exports the panel and the reference bundle for the Xenium Panel Designer, and stage 11
  checks the probe report the Designer returns. Stages 05, 08 and 09 evaluate a panel on a labelled
  Xenium run of the target, when one exists. CP10k, counts per 10{,}000; PCA, principal component
  analysis; UMAP, uniform manifold approximation and projection.}
  \label{fig:pipeline}
\end{figure}

% ------------------------------------------------------------------------------
\subsection{Datasets}
\label{sec:results-datasets}

We used five paired datasets, each with a single-cell reference and a Xenium run, and one
single-cell atlas (Table~\ref{tab:datasets}). Three pairs serve as \emph{targets}, which need
cell-type labels on the Xenium cells to score a panel: ovarian cancer, cervical cancer and mouse brain.
All five pairs serve as Xenium priors. Ovarian and cervical cancer are the reference cases: formalin-fixed
paraffin-embedded (FFPE) tissue profiled with the 10x Genomics Chromium Flex assay (Flex) and Xenium
Prime 5K on the same tissue block, with 10x's cell-type annotation of the Xenium cells. Mouse brain
adds a fresh-frozen (FF) target of another species. The mouse retina pair supplies a same-species
prior for mouse brain; it has no cell-type labels, so it cannot be a target. The breast cancer pair is
used only as a prior: its Flex reference and its Xenium run come from different tumours, so it is not
suitable as a target. The mouse RGC atlas, with 45 annotated subtypes, has no Xenium
counterpart with subtype labels, so it is used for a held-out single-cell benchmark of fine subtypes.

\begin{table}[!htbp]
  \centering\footnotesize
  \caption{\textbf{Datasets used in this study.} Targets are scored on the held-out half of their
  Xenium run; every pair also serves as a Xenium prior. Cell counts are those used here (for the
  targets' Xenium runs, after quality control). Genes: the genes measured in both the single-cell
  reference and the Xenium panel (for the RGC atlas, its genes on the mouse Xenium 5K panel). FFPE, formalin-fixed paraffin-embedded; FF, fresh-frozen; Flex, 10x
  Genomics Chromium Flex; snRNA, single-nucleus RNA sequencing; scRNA, single-cell RNA sequencing.
  CZ, Chan Zuckerberg; GEO, Gene Expression Omnibus. Full links are in the Data availability
  statement.}
  \label{tab:datasets}
  \resizebox{0.995\textwidth}{!}{%
  \begin{tabular}{@{}>{\raggedright}p{2.2cm}>{\raggedright}p{1.1cm}>{\raggedright}p{3.9cm}>{\raggedright}p{3.0cm}c>{\raggedright}p{2.6cm}>{\raggedright\arraybackslash}p{3.5cm}@{}}
    \toprule
    Dataset & Species & Single-cell reference & Xenium & Genes & Role & Source \\
    \midrule
    Ovarian cancer & human & FFPE Flex; 16{,}187 cells; 14 cell types & Prime 5K; FFPE; same block; 363{,}311 cells & 4{,}766 & target; prior &
      \href{https://www.10xgenomics.com/datasets/17k-human-ovarian-cancer-scFFPE}{10x Flex},
      \href{https://www.10xgenomics.com/datasets/xenium-prime-ffpe-human-ovarian-cancer}{10x Xenium} \\
    Cervical cancer & human & FFPE Flex; 9{,}475 cells; 6 cell types & Prime 5K; FFPE; same block; 366{,}273 cells & 4{,}548 & target; prior &
      \href{https://www.10xgenomics.com/datasets/17k-human-cervical-cancer-scFFPE}{10x Flex},
      \href{https://www.10xgenomics.com/datasets/xenium-prime-ffpe-human-cervical-cancer}{10x Xenium} \\
    Mouse brain & mouse & snRNA (10x 3$'$ v3), aging atlas~\citep{allen2023brainaging}; 79{,}667 nuclei; 12 cell types & Prime 5K; FF coronal section; 63{,}173 cells & 4{,}791 & target; prior &
      \href{https://cellxgene.cziscience.com/collections/31937775-0602-4e52-a799-b6acdd2bac2e}{CZ CELLxGENE},
      \href{https://www.10xgenomics.com/datasets/xenium-prime-fresh-frozen-mouse-brain}{10x Xenium} \\
    Mouse retina & mouse & snRNA multiome, retina and pigment epithelium, 12-week mice~\citep{lyu2025retinaaging}; 10{,}118 nuclei & Xenium 5K mouse panel; FF; 12-week mice; 70{,}558 cells & 5{,}006 & prior &
      \href{https://www.ncbi.nlm.nih.gov/geo/query/acc.cgi?acc=GSE325478}{GEO GSE325478},
      \href{https://www.ncbi.nlm.nih.gov/geo/query/acc.cgi?acc=GSE307031}{GEO GSE307031} \\
    Breast cancer & human & FFPE Flex~\citep{janesick2023xenium}; 44{,}863 cells with at least 200 genes; a different tumour from the Xenium & Prime 5K; FFPE; 699{,}110 cells & 5{,}086 & prior &
      \href{https://www.ncbi.nlm.nih.gov/geo/query/acc.cgi?acc=GSM7782698}{GEO GSM7782698},
      \href{https://www.10xgenomics.com/datasets/xenium-prime-ffpe-human-breast-cancer}{10x Xenium} \\
    RGC atlas & mouse & scRNA (10x)~\citep{tran2019rgc}; 34{,}029 cells; 45 subtypes & --- & 4{,}911 & single-cell benchmark &
      \href{https://www.ncbi.nlm.nih.gov/geo/query/acc.cgi?acc=GSE133382}{GEO GSE133382} \\
    \bottomrule
  \end{tabular}}
\end{table}

Several other datasets were considered and set aside. A skin melanoma pair was removed because its
single-cell reference turned out to be from mouse lemur while its Xenium run was human, so its
detection rates measured a species mismatch. A dermatomyositis skin pair had a nearly flat detection
profile from a shallow reference and was not validated. A second ovarian cancer Xenium dataset was
measured with an earlier 5K panel version lacking about 30\% of the genes, and in situ
whole-transcriptome Atera data show no detection loss to correct, so neither tests detection-aware
design. A human spleen pair was set aside because the source of its Xenium run could not be
documented.

% ------------------------------------------------------------------------------
\subsection{Detection loss is large, gene-specific and reproducible across experiments}
\label{sec:results-detection}

A Xenium prior is only useful if a gene's detection loss measured in one experiment predicts its loss
in another. On the ovarian cancer Xenium, the median gene is detected in 52\% as many cells as in the
matched single-cell reference (4{,}259 genes detected in at least 0.1\% of reference cells), and the
loss varies widely between genes (Fig.~\ref{fig:detrepro}a). The loss is reproducible. Per-gene
$\pkeep$ in ovarian and cervical cancer, two different tissues profiled on the same Xenium panel,
correlate at Spearman $\rho=0.88$ over 3{,}056 shared genes. That is more strongly than the genes'
single-cell detection breadth correlates between the two tissues ($\rho=0.62$), so $\pkeep$ captures
a property of the gene and its probes rather than shared biology (Fig.~\ref{fig:detrepro}b).

\begin{figure}[!htbp]
  \centering
  \includegraphics[width=0.86\textwidth]{figures/fig_detection_reproducibility.pdf}
  \caption{\textbf{The single-cell-to-Xenium detection loss is large, gene-specific and reproducible
  across tissues.} \textbf{(A)}~Per-gene detection rate (fraction of cells with a nonzero count) in
  the ovarian single-cell reference versus ovarian Xenium, for the 4{,}259 genes detected in at
  least 0.1\% of reference cells. Colour is $\pkeep$. The median gene keeps 52\% of its detection.
  \textbf{(B)}~Per-gene $\pkeep$ in ovarian versus cervical cancer, two tissues profiled on the same
  Xenium 5K panel, for the 3{,}056 genes expressed in at least 1\% of reference cells in both
  (Spearman $\rho=0.88$; dashed line $y=x$, red line least-squares fit). The loss reproduces more
  faithfully than single-cell detection breadth ($\rho=0.62$).}
  \label{fig:detrepro}
\end{figure}

% ------------------------------------------------------------------------------
\subsection{Designing a panel without the target's Xenium}
\label{sec:results-design}

Every panel in this study is designed and evaluated in the same way (Fig.~\ref{fig:design}). Three
datasets are involved. The \emph{single-cell reference} is annotated single-cell data of the target
tissue; the panel is chosen from its genes. The \emph{Xenium prior} supplies per-gene detection rates
from a previous experiment that measured both a single-cell reference and a Xenium run; only its
detection rates are used. The \emph{target Xenium} is the Xenium run of the target tissue. Its
Xenium cells were split into two spatial halves by a checkerboard, and only the held-out half was
scored. Nothing from the target Xenium informs the panel. To score a panel, a classifier is trained on
the single-cell reference restricted to the panel's genes, it predicts the cell type of every
held-out target Xenium cell, and the predictions are compared with the target's labels by \Fone{}, the
unweighted mean of the per-cell-type $F_1$ scores.

\begin{figure}[!htbp]
  \centering
  \resizebox{\textwidth}{!}{%
  \begin{tikzpicture}[
    font=\footnotesize,
    every node/.append style={execute at begin node={\hyphenpenalty=10000\exhyphenpenalty=10000}},
    arr/.style={-{Stealth[length=2.6mm,width=2mm]}, thick, draw=gray!70},
    frame/.style={draw, rounded corners=3pt},
    hdr/.style={font=\bfseries, anchor=base},
    box/.style={draw, rounded corners=2pt, align=center, minimum height=11mm}
  ]
    \draw[frame] (0,0) rectangle (6.8,-4.6);
    \draw[frame] (7.8,0) rectangle (12.6,-4.6);
    \draw[frame] (13.6,0) rectangle (19.6,-4.6);
    \node[hdr] at (3.4,0.25)  {Inputs available before ordering};
    \node[hdr] at (10.2,0.25) {Panel design};
    \node[hdr] at (16.6,0.25) {Evaluation};
    \node[box, fill=orange!10, draw=orange!75, dashed, text width=58mm] (pri) at (3.4,-1.25)
      {\textbf{Xenium prior}\\per-gene detection rates from a previous\\single-cell and Xenium experiment};
    \node[box, fill=blue!8, text width=58mm] (ref) at (3.4,-3.35)
      {\textbf{single-cell reference}\\annotated cells of the target tissue};
    \node[box, fill=orange!15, draw=orange!70, text width=40mm] (des) at (10.2,-1.6)
      {thin the reference to the\\prior's detection rates,\\then select genes};
    \node[box, fill=green!11, text width=40mm] (pan) at (10.2,-3.45) {gene panel};
    \draw[arr, draw=orange!75] (pri.east)--([yshift=2.5mm]des.west);
    \draw[arr] (ref.east)--([yshift=-2.5mm]des.west);
    \draw[arr] (des.south)--(pan.north);
    \node[box, fill=violet!12, text width=54mm] (tx) at (16.6,-1.25)
      {\textbf{target Xenium}\\held-out half of the target's run};
    \node[box, fill=gray!8, text width=54mm] (ev) at (16.6,-3.35)
      {classifier trained on the reference\\predicts each target cell;\\\Fone{} against the target's labels};
    \draw[arr] (pan.east)--([yshift=-2.5mm]ev.west);
    \draw[arr] (tx.south)--(ev.north);
  \end{tikzpicture}}
  \caption{\textbf{Designing and evaluating a panel without the target's Xenium.} The Xenium prior
  and the single-cell reference are available before the panel is ordered. The reference is thinned
  to the prior's per-gene detection rates (\dart{}) or used as measured (the other methods), and a
  panel is selected. The panel is evaluated on the held-out half of the target Xenium: a classifier
  trained on the reference, restricted to the panel's genes, predicts each cell, and the predictions
  are scored by \Fone{} against the target's labels, which are used only here.}
  \label{fig:design}
\end{figure}

% ------------------------------------------------------------------------------
\subsection{A Xenium prior helps when it matches the target's chemistry and species}
\label{sec:results-priors}

We first asked which Xenium priors help \dart{}. For each target we designed \dart{} panels with every
available prior, and with none, and compared them at the target's default panel size
(Fig.~\ref{fig:transfer}; Methods). The target's own Xenium is shown as an upper bound that no
designer has before ordering.

The prior that helped most, other than the target's own, always matched the target's chemistry and
species. For the two human FFPE tumours it was the other tumour, profiled with the same Flex reference
and Xenium chemistry: cervical cancer raised ovarian cancer from 0.470 \Fone{} without a prior to
0.563, against 0.600 with the target's own Xenium, and ovarian cancer raised cervical cancer from
0.517 to 0.612 (own 0.666). The breast pair, of the same chemistry but with its reference and Xenium
run from different tumours, helped less (0.520 and 0.538). For the fresh-frozen mouse brain, the
fresh-frozen mouse retina prior helped (0.643 to 0.673; own 0.700), whereas every human FFPE prior
lowered accuracy, by 0.05 to 0.13. Conversely, the two fresh-frozen mouse priors helped ovarian cancer
little ($+0.011$ and $+0.025$) and lowered cervical cancer ($-0.016$ and $-0.033$). A prior from the
other species reaches only the genes whose human and mouse symbols agree, about half of the target's
genes (Methods). The best results therefore need a prior of the same chemistry and species as the
planned experiment, which can come from a different tissue.

\begin{figure}[!htbp]
  \centering
  \includegraphics[width=\textwidth]{figures/fig_transfer.pdf}
  \caption{\textbf{Which Xenium priors help \dart{}.} \Fone{} on the held-out target Xenium of
  \dart{} panels designed with each Xenium prior (columns) for each target (rows). Each prior's curve
  ($K=1$--10) is interpolated at the sizes of the target's benchmark panels with $K=3$--6 (range in the
  row label) and averaged. The upper number is \Fone{}; the lower number and the colour give the
  difference from designing with no prior. ``Target's own'' uses the target's own Xenium, which is
  unavailable before ordering, and is shown as an upper bound. A prior from the other species reaches
  only the genes whose symbols agree (Methods). FFPE, formalin-fixed paraffin-embedded; FF,
  fresh-frozen.}
  \label{fig:transfer}
\end{figure}

% ------------------------------------------------------------------------------
\subsection{Benchmark against existing marker-selection methods}
\label{sec:results-benchmark-design}

We then compared \dart{} with four existing methods: the Wilcoxon rank-sum test, COSG, NS-Forest and
\spapros{}. Each target used a Xenium prior from a different tissue, chosen by how closely its samples
match the target's and not by any score on the target: for ovarian and cervical cancer, each other
(FFPE tissue with a Flex reference and Xenium Prime 5K on the same block), and for mouse brain, the
mouse retina pair, the only other mouse pair, which like the target is fresh-frozen tissue with a
single-nucleus reference and the mouse 5K panel (Fig.~\ref{fig:bench-example}). The other methods
select genes from the single-cell reference as measured, as they are normally used. Because panel size
drives accuracy, \dart{} was run with $K=1$--10 rules per cell type and the Wilcoxon test and COSG with
1--12 genes per cell type, and methods were compared at matched realised panel size by interpolating
along each accuracy curve. NS-Forest's panel barely changes with its budget (21--24 genes on ovarian
cancer and one identical panel on each of the other targets), and \spapros{} was run at its default
size, so neither gives an accuracy curve and \dart{} was compared with them at their own panel
sizes.

\begin{figure}[!htbp]
  \centering
  \resizebox{\textwidth}{!}{%
  \begin{tikzpicture}[
    font=\footnotesize,
    every node/.append style={execute at begin node={\hyphenpenalty=10000\exhyphenpenalty=10000}},
    arr/.style={-{Stealth[length=2.6mm,width=2mm]}, thick, draw=gray!70},
    box/.style={draw, rounded corners=2pt, align=center, minimum height=14mm}
  ]
    \node[box, fill=orange!10, draw=orange!75, dashed, text width=30mm] (pri) at (0,0.9)
      {\textbf{Xenium prior}\\cervical cancer\\(Flex + Xenium 5K)};
    \node[box, fill=blue!8, text width=30mm] (ref) at (0,-1.0)
      {\textbf{single-cell reference}\\ovarian cancer Flex\\(14 cell types)};
    \node[box, fill=orange!15, draw=orange!70, text width=38mm] (sel) at (4.95,-0.05)
      {\textbf{select a panel}\\\dart{} (with the prior);\\Wilcoxon, COSG, NS-Forest,\\\spapros{} (reference only)};
    \node[box, fill=gray!8, text width=34mm] (cls) at (9.8,-0.05)
      {\textbf{classify ovarian cell types}\\classifier trained on the\\ovarian reference, panel genes};
    \node[box, fill=violet!12, text width=34mm] (ev) at (14.5,-0.05)
      {\textbf{evaluate on ovarian Xenium}\\held-out half; \Fone{}\\at matched panel size};
    \draw[arr, draw=orange!75] (pri.east)--([yshift=2.5mm]sel.west);
    \draw[arr] (ref.east)--([yshift=-2.5mm]sel.west);
    \draw[arr] (sel.east)--(cls.west);
    \draw[arr] (cls.east)--(ev.west);
  \end{tikzpicture}}
  \caption{\textbf{Benchmark design, shown for the ovarian cancer target.} The cervical cancer Xenium
  prior and the ovarian cancer single-cell reference are used to select a panel; a classifier trained
  on the ovarian reference with the panel's genes classifies the cells of the held-out half of the
  ovarian Xenium, and \Fone{} is compared at matched panel size. The same design is used for the
  cervical cancer target (ovarian prior) and the mouse brain target (mouse retina prior). Only \dart{}
  uses the prior; the other methods select from the reference as measured.}
  \label{fig:bench-example}
\end{figure}

% ------------------------------------------------------------------------------
\subsection{\dart{} outperforms existing marker-selection methods}
\label{sec:results-main}

At matched panel size, \dart{} outperformed the Wilcoxon test, COSG, NS-Forest and \spapros{} on every
target (Fig.~\ref{fig:benchmark}; Table~\ref{tab:headline}). On ovarian cancer, the mean advantage over
the Wilcoxon test was $+0.064$ \Fone{}, ranging from $+0.036$ to $+0.118$ across panel sizes, and
\dart{} reached the Wilcoxon test's best accuracy (0.581 with 144 genes) with 83 genes. On mouse brain
the advantage was $+0.105$, and \dart{} needed 41 genes to match the Wilcoxon test's best, against
its 132. On cervical cancer it was $+0.044$, with 28 genes against 58. Across the three targets the
margins were $+0.038$ to $+0.079$ over COSG, $+0.099$ to $+0.175$ over NS-Forest and $+0.044$ to
$+0.082$ over \spapros{}. When \dart{}'s thinning and rule learning were repeated at four seeds, every
margin stayed positive (the smallest was $+0.039$, over \spapros{} on ovarian cancer), and the
seed-to-seed standard deviation of a margin was 0.005--0.017 (Methods).

\begin{figure}[!htbp]
  \centering
  \includegraphics[width=\textwidth]{figures/fig_benchmark.pdf}
  \caption{\textbf{\dart{} outperforms existing marker-selection methods.} \textbf{(A)}~Ranked
  summary over the three targets. On each target, each method is represented by one panel near
  \dart{}'s default size (\dart{} with $K=4$; the Wilcoxon test and COSG at the per-type budget closest
  in size; NS-Forest at its largest panel; \spapros{} at its default), and methods are ranked by their
  mean \Fone{}. Columns give the panel sizes (ovarian / mouse brain / cervical), \Fone{} on each target
  and its mean, the difference between \dart{} and the method at the method's panel size averaged over
  the targets, and compute measured on this study's runs: central processing unit (CPU) time and wall
  time averaged over the targets, and the largest peak memory (Supplementary Table~\ref{tab:compute}).
  Bold marks the best value in each column. \textbf{(B--D)}~\Fone{} on the held-out target Xenium against realised panel size for \dart{}
  with a Xenium prior ($K=1$--10) and for the Wilcoxon test and COSG (1--12 genes per cell type),
  NS-Forest and \spapros{}. Mouse brain is scored on rebuilt labels (Supplementary~S3).}
  \label{fig:benchmark}
\end{figure}

\begin{table}[!htbp]
  \centering\small
  \caption{\textbf{\dart{} (Xenium prior) versus each method at matched panel size.} Mean \Fone{}
  difference over \dart{}'s realised sizes, with the range across sizes in brackets. NS-Forest (mean
  over its three budgets) and \spapros{} are compared at their own panel sizes. The last column gives
  the number of genes \dart{} needs to match the Wilcoxon test's best accuracy and, in brackets, the
  number the Wilcoxon test itself uses. Seed 42; across four seeds the standard deviation of each
  margin is 0.005--0.017 (Methods). Mouse brain is scored on rebuilt labels (Supplementary~S3).}
  \label{tab:headline}
  \resizebox{\textwidth}{!}{%
  \begin{tabular}{@{}lccccc@{}}
    \toprule
     & \multicolumn{4}{c}{\dart{} minus each method} & \dart{} genes to reach \\
    \cmidrule(lr){2-5}
    Target (Xenium prior) & Wilcoxon & COSG & NS-Forest & \spapros{} & Wilcoxon's best \\
    \midrule
    Ovarian (cervical) & $+0.064$ [$+0.036$, $+0.118$] & $+0.054$ [$+0.030$, $+0.078$] & $+0.099$ & $+0.044$ & 83 (144) \\
    Mouse brain (retina) & $+0.105$ [$+0.023$, $+0.230$] & $+0.038$ [$-0.018$, $+0.217$] & $+0.170$ & $+0.054$ & 41 (132) \\
    Cervical (ovarian) & $+0.044$ [$+0.018$, $+0.071$] & $+0.079$ [$+0.015$, $+0.214$] & $+0.175$ & $+0.082$ & 28 (58) \\
    \bottomrule
  \end{tabular}}
\end{table}

Two qualifications apply to COSG. First, on mouse brain \dart{}'s lead over COSG is concentrated in
small panels: it is $+0.041$ to $+0.217$ below about 55 genes, but from about 65 genes upward the two
are level ($-0.018$ to $+0.017$), and COSG's best point (0.708 with 120 genes) is slightly above
\dart{}'s best (0.703). On cervical, COSG's best point (0.656 with 72 genes) lies beyond the largest
panel on \dart{}'s curve (50 genes). Second, COSG's accuracy depends on how the reference is
normalised; normalising over the full transcriptome rather than the shared gene universe would raise
COSG by about 0.015 on ovarian and 0.036 on cervical (Supplementary~S2).

Fig.~\ref{fig:benchmark}a ranks one panel per method near \dart{}'s default size on each target.
\dart{} ranked first on every target and by mean \Fone{} (0.610), ahead of \spapros{} (0.565), COSG
(0.558), the Wilcoxon test (0.509) and NS-Forest (0.389). At each method's own panel size, \dart{} led
\spapros{} by 0.060, COSG by 0.052, the Wilcoxon test by 0.099 and NS-Forest by 0.150 on average over
the three targets.

The methods also differ in their compute (Fig.~\ref{fig:benchmark}a; Supplementary
Fig.~\ref{fig:sfig_compute}). \dart{} splits its rule learning into independent tasks, one per cell
type and learner, and runs them on parallel worker processes. It used the most CPU time, 1.2--8.9
core-hours per target against 0.3--1.1 for \spapros{} and at most 1.8 for the marker selectors, but with
eight workers it finished in 12--82 minutes, six to seven times faster than on one core, and its largest
process needed at most 6.3~GB of memory. \spapros{} took 11--28 minutes and needed 16--25~GB, and the
Wilcoxon test and COSG finished in under three minutes.

% ------------------------------------------------------------------------------
\subsection{Per-cell-type accuracy on the target Xenium}
\label{sec:results-pertype}

The differences in \Fone{} come from particular cell types (Fig.~\ref{fig:pertype}). On ovarian
cancer, \dart{}'s panel gave the highest $F_1$ of the five methods for 6 of the 14 cell types and
\spapros{}'s larger panel for 5; \dart{} scored above the Wilcoxon test on 11 types and above
NS-Forest on 12. Its largest gain was on T and natural killer (NK) cells (0.50, against 0.14--0.27 for
the other methods). It fell furthest behind the best method, by 0.14--0.17, on two tumour states,
proliferative and VEGFA-positive tumour cells, which the Wilcoxon test and \spapros{} separated
better. On mouse brain, \dart{} gave the highest $F_1$ for 6 of 12 cell types and scored above the
Wilcoxon test on 11, most of all on neurons (0.86 against 0.24). The two rarest mouse brain types, macrophages (48 held-out cells) and T cells (36),
were poorly classified by every method ($F_1$ at most 0.38). On cervical cancer, with six broad cell
types, \dart{}'s advantage came from macrophages (0.64, against 0.09--0.30), and it was weakest on
squamous carcinoma (0.28, against 0.29--0.43).

\begin{figure}[!htbp]
  \centering
  \includegraphics[width=\textwidth]{figures/fig_pertype.pdf}
  \caption{\textbf{Per-cell-type $F_1$ on each target Xenium.} $F_1$ of each cell type (columns,
  ordered by the number of held-out Xenium cells, in brackets) for one panel per method near
  \dart{}'s default size (rows; panel size in brackets), as in Fig.~\ref{fig:benchmark}a. Bold marks
  the best method for each cell type. \textbf{(A)}~Ovarian cancer (cervical prior). \textbf{(B)}~Mouse
  brain (retina prior; rebuilt labels). \textbf{(C)}~Cervical cancer (ovarian prior).}
  \label{fig:pertype}
\end{figure}

% ------------------------------------------------------------------------------
\subsection{Number of rules per cell type and rule overlap}
\label{sec:results-ko}

Two settings control a \dart{} panel: $K$, the number of rules kept per cell type, and $O_{\max}$, the
number of genes that two rules of the same cell type may share (Fig.~\ref{fig:ko}). Each additional
rule adds genes, so $K$ also sets the panel size. \Fone{} rose with $K$ on every target, steeply from
$K=1$ to $K=3$ and more slowly afterwards. On ovarian cancer it rose from 0.455 with one rule per type (18
genes) to 0.555 at the default $K=4$ (58 genes) and 0.623 at $K=10$ (128 genes); on mouse brain from
0.574 to 0.680 and 0.703; and on cervical cancer from 0.499 to 0.595, reaching 0.643 at $K=9$ before
falling to 0.629 at $K=10$. The effect of letting rules share one gene ($O_{\max}=1$) depended on the
target. On ovarian cancer it lowered \Fone{} at every $K$ above 1, by 0.085 at $K=4$. On mouse brain
the two settings differed by at most 0.024, with no consistent sign. On cervical cancer sharing was
better from $K=4$ upward, by 0.014 at $K=4$ and 0.043 at $K=10$.

\begin{figure}[!htbp]
  \centering
  \includegraphics[width=\textwidth]{figures/fig_ko.pdf}
  \caption{\textbf{\dart{} accuracy as a function of the number of rules per cell type and their
  overlap.} \Fone{} on the held-out target Xenium against $K$, the number of rules kept per cell type,
  for gene-disjoint rules ($O_{\max}=0$, the default) and rules that may share one gene
  ($O_{\max}=1$), with each target's Xenium prior. Numbers give the panel size at $K=4$ and $K=10$.
  The dotted line marks the default $K=4$.}
  \label{fig:ko}
\end{figure}

% ------------------------------------------------------------------------------
\subsection{\dart{}'s rules are interpretable and agree with the classifier}
\label{sec:results-rules}

Each \dart{} panel comes with explicit rules, such as a gene above a threshold, alone or combined with
others. These rules name recognisable marker programs (Table~\ref{tab:rules-example}). For the
interferon-high inflammatory tumour cells of the ovarian cancer reference, \dart{}'s four gene-disjoint
rules use \emph{MX1} (in a conjunction that also requires \emph{ESR1} high and \emph{NAMPT} and
\emph{CD200} low), \emph{IFIT1}, \emph{IFIT3} and \emph{DDX58}, all interferon-stimulated genes. The
rules for ciliated epithelial cells use cilia genes (\emph{CFAP100}, \emph{CDHR4}, \emph{TEKT1},
\emph{DNAH9}), those for proliferative tumour cells use cell-cycle genes (\emph{CIP2A}, \emph{RRM2},
\emph{TYMS}, \emph{TPX2}), and those for T and natural killer cells use \emph{TRAC}, \emph{CD8A},
\emph{CXCR4} and \emph{IL7R}. A ranked gene list carries no threshold and does not say which genes act
together.

\begin{table}[!htbp]
  \centering\small
  \caption{\textbf{Example \dart{} rules.} The four gene-disjoint rules kept for selected cell types
  of the ovarian cancer reference (cervical Xenium prior, $K=4$). Thresholds are on the log-normalised
  scale of the thinned reference. $P$ and $R$, precision and recall of the rule in the thinned
  reference.}
  \label{tab:rules-example}
  \begin{tabular}{@{}>{\raggedright}p{3.4cm}>{\raggedright}p{7.6cm}cc@{}}
    \toprule
    Cell type & Rule & $P$ & $R$ \\
    \midrule
    \multirow{4}{3.4cm}{Inflammatory tumour cells} & $\mathit{MX1}>3.95 \wedge \mathit{ESR1}>2.40 \wedge \mathit{NAMPT}\le4.19 \wedge \mathit{CD200}\le4.41$ & 0.56 & 0.33 \\
     & $\mathit{IFIT1}>1.17$ & 0.25 & 0.19 \\
     & $\mathit{IFIT3}>2.52$ & 0.15 & 0.27 \\
     & $\mathit{DDX58}>4.68$ & 0.23 & 0.17 \\
    \midrule
    \multirow{4}{3.4cm}{Ciliated epithelial cells} & $\mathit{CFAP100}>4.17 \wedge \mathit{SLC2A3}\le1.21$ & 0.78 & 0.73 \\
     & $\mathit{CDHR4}>3.65$ & 0.82 & 0.56 \\
     & $\mathit{TEKT1}>2.59$ & 0.58 & 0.46 \\
     & $\mathit{DNAH9}>2.77$ & 0.72 & 0.32 \\
    \midrule
    \multirow{4}{3.4cm}{Proliferative tumour cells} & $\mathit{CIP2A}>2.79$ & 0.62 & 0.59 \\
     & $\mathit{RRM2}>2.41$ & 0.71 & 0.52 \\
     & $\mathit{TYMS}>2.45$ & 0.70 & 0.49 \\
     & $\mathit{TPX2}>2.75$ & 0.58 & 0.52 \\
    \bottomrule
  \end{tabular}
\end{table}

Panels are deployed with a classifier that never sees the rules, yet the classifier's coefficients
recover them (Table~\ref{tab:rules}). A ``$\textit{gene}>t$'' condition in a cell type's rule carries a
positive coefficient for that type in 98--100\% of cases on the three Xenium targets and in 96.5\%
across the 45 RGC subtypes. Rule genes carry 2.0--3.0 times the weight of the panel's other genes, and
a cell type's largest coefficient belongs to one of its rule genes for every cell type of the three
targets and for 92\% of the RGC subtype panels. The rules are therefore an accurate, human-readable account of why the classifier calls a
cell type.

\begin{table}[!htbp]
  \centering\small
  \caption{\textbf{\dart{}'s rules agree with the deployed classifier.} The classifier that scores each
  panel (Methods), for the benchmark's $K=4$ panel of each Xenium target and, for the RGC atlas, each
  fold's $K=4$ panel with that fold's classifier, pooled over the five folds. ``$>$'' and ``$\le$''
  are the fractions of rule conditions whose gene has a positive or negative coefficient,
  respectively, for the rule's cell type (number of conditions in brackets). Weight ratio is the mean
  $|$coefficient$|$ of a type's rule genes over that of the panel's other genes.}
  \label{tab:rules}
  \resizebox{\textwidth}{!}{%
  \begin{tabular}{@{}lcccc@{}}
    \toprule
    Panel (cell types) & ``$>$'' $\to$ positive & ``$\le$'' $\to$ negative & weight ratio & top coefficient is a rule gene \\
    \midrule
    Ovarian (14) & 100\% (57) & 83\% (6) & 2.3$\times$ & 100\% \\
    Mouse brain (12) & 98\% (49) & 86\% (7) & 2.2$\times$ & 100\% \\
    Cervical (6) & 100\% (20) & 100\% (6) & 2.0$\times$ & 100\% \\
    RGC subtypes (45; 5 folds) & 96.5\% (1{,}522) & 83.8\% (597) & 3.0$\times$ & 92\% \\
    \bottomrule
  \end{tabular}}
\end{table}

% ------------------------------------------------------------------------------
\subsection{Fine retinal ganglion cell subtypes}
\label{sec:results-rgc}

The Xenium targets have 6--14 broad cell types, which a single marker usually separates; on them most
\dart{} rules use one gene. RGCs, the projection neurons of the retina, comprise more than 40
transcriptionally related subtypes~\citep{sanes2015rgc,tran2019rgc}, many without a unique marker. No
Xenium dataset has RGC subtype labels, so we evaluated the 45 subtypes of a published atlas by
five-fold out-of-fold prediction within the single-cell data (Methods): each panel is designed on four
folds and scored on the held-out fold. Here 75\% of \dart{}'s rules are multi-gene conjunctions, with a
mean of 2.4 genes per rule ($K=4$ panels of the five folds), against 7--15\% on the three Xenium
targets.

\dart{}, the Wilcoxon test and \spapros{} reached the same accuracy at matched size, about 0.955 to
0.96 \Fone{} from 150 genes upward (Fig.~\ref{fig:rgc}a): \dart{} scored 0.959 with 184 genes and 0.960
with 223, the Wilcoxon test 0.957 with 191 and 0.958 with 224, and \spapros{} 0.956 with 175. COSG
was 0.04--0.06 lower at matched size and reached 0.945 only with 393 genes, and NS-Forest's single
panel (71 genes) scored 0.879. \dart{} led COSG ($+0.028$ to $+0.037$) and NS-Forest ($+0.052$ to
$+0.076$) in every third of subtype abundance, from the rarest to the commonest, whereas against the
Wilcoxon test and \spapros{} the per-subtype differences stayed within $\pm0.007$ at every abundance
(Fig.~\ref{fig:rgc}b). No single probe was load-bearing, consistent with each subtype being covered
by several gene-disjoint rules: among the 230 single-gene knockouts of one fold's $K=4$ panel, the
largest drop in \Fone{} was 0.003.

\begin{figure}[!htbp]
  \centering
  \includegraphics[width=\textwidth]{figures/fig_rgc.pdf}
  \caption{\textbf{Forty-five RGC subtypes, held out within the single-cell atlas.} \textbf{(A)}~Out-of-fold
  \Fone{} against panel size for \dart{} ($K=1$--4), the Wilcoxon test and COSG (4--12 genes per
  subtype), \spapros{} (50, 100 and 150 genes requested, and its default of 170--182 genes) and
  NS-Forest. \textbf{(B)}~Mean per-subtype $F_1$ difference between \dart{} and each method, by subtype
  abundance (thirds of the 45 subtypes), each at the closest panel sizes: \dart{} with 223 genes against
  the Wilcoxon test with 224 and COSG with 238, \dart{} with 184 against \spapros{} with 175, and
  \dart{} with 84 against NS-Forest with 71.}
  \label{fig:rgc}
\end{figure}

% ------------------------------------------------------------------------------
\subsection{Checking a panel's probes for off-target expression}
\label{sec:results-offtarget}

The off-target check applies to a panel from any method once its probes are designed.
Fig.~\ref{fig:offtarget} shows its output for a 176-gene human retina panel designed in a separate
study, using the probe report returned by the Xenium Panel Designer and a single-nucleus atlas of the
human retina~\citep{li2026retinaatlas}. The probe for the cone marker \emph{CRX} has one predicted
off-target, \emph{RPS6KL1}, which is expressed weakly and evenly across the retinal cell classes, so
it cannot mimic the cone pattern and the probe can be kept. Three of the probes for the amacrine-cell marker \emph{PAX6} have predicted off-targets. Of
those measured in the atlas, \emph{ALX1} is barely expressed, but \emph{PAX2}, \emph{PAX5} and
\emph{PAX8} are strongly expressed in astrocytes, where \emph{PAX6} itself is absent. Those probes
could report amacrine-like signal in astrocytes, so they are flagged for replacement.

\begin{figure}[!htbp]
  \centering
  \includegraphics[width=0.32\textwidth]{figures/fig_offtarget_CRX.png}\hspace{2em}
  \includegraphics[width=0.36\textwidth]{figures/fig_offtarget_PAX6.png}
  \caption{\textbf{Example output of the off-target check.} Dotplots of a marker (left column of each
  plot) beside its predicted off-target genes across the major cell classes of a human retina
  single-nucleus atlas~\citep{li2026retinaatlas}; dot size is the fraction of cells expressing the
  gene and colour its mean expression. \textbf{(A, left)}~The cone marker \emph{CRX}: its off-target
  \emph{RPS6KL1} is expressed weakly and evenly across classes, so the probe is kept. \textbf{(B, right)}~The
  amacrine-cell marker \emph{PAX6}: of its off-targets, \emph{PAX2}, \emph{PAX5} and \emph{PAX8} are
  expressed in astrocytes, so its probes are flagged. Panel from a 176-gene human retina design. AC,
  amacrine cells; BC, bipolar cells; HC, horizontal cells; MG, M\"uller glia; RGC, retinal ganglion
  cells; RPE, retinal pigment epithelium.}
  \label{fig:offtarget}
\end{figure}

% ==============================================================================
\section{Discussion}
\label{sec:discussion}

\xspear{} turns an annotated single-cell reference into a Xenium panel without any data from the
target sample. It makes three contributions:
\begin{enumerate}[leftmargin=*,label=\textbf{\arabic*.}]
  \item \textbf{Detection-aware design from a Xenium prior.} The gap between a gene's detection in a
    single-cell reference and in Xenium is large, gene-specific and reproducible across tissues.
    Thinning the reference to a previous experiment's detection rates anticipates that gap before
    ordering. The prior should match the planned experiment's chemistry and species, and it can come
    from another tissue; a mismatched prior can lower accuracy.
  \item \textbf{\dart{}.} With a Xenium prior from a different tissue, \dart{} outperforms the
    Wilcoxon test, COSG, NS-Forest
    and \spapros{} at matched panel size on every target we tested, and reaches the Wilcoxon test's
    best accuracy with 42--69\% fewer genes. Its rules name interpretable marker programs and agree
    with the classifier that uses the panel. On fine RGC subtypes it matches the best methods, leads
    COSG and NS-Forest at every subtype abundance, and no single probe is load-bearing.
  \item \textbf{A method-independent off-target check.} Dotplots of each marker beside its predicted
    off-target genes show which probes would add signal in the wrong cell types, for a panel chosen
    by any method.
\end{enumerate}

Two limitations matter most. First, the benefit depends on a suitable Xenium prior: for the best
results, the prior should come from the same assay chemistry and the same species as the target.
Second, the evaluation labels of the Xenium targets derive from reference-based annotation: 10x's for
ovarian and cervical cancer and our own label transfer for mouse brain. The comparisons are therefore
within the reference's annotation scheme, although an independent annotation of the ovarian section
agreed with the evaluation labels on 82.8\% of cells at the level of five lineages
(Supplementary~S3).

The thinning, selection and off-target modules are separable. Other targeted platforms can be
supported once a detection profile and a probe-design output exist for them, and the same
architecture can host objectives beyond cell-type identification, such as panels that preserve
differential expression across conditions or spatial niches.

% ==============================================================================
\section{Methods}
\label{sec:methods}

\subsection{The \xspear{} software}
\label{sec:methods-software}

\paragraph{Interfaces.} \xspear{} is a Python package (Code availability statement) with three
interfaces. As a library, \texttt{design\_panel} takes an annotated single-cell reference, a dataset
configuration and, optionally, a Xenium prior, and returns the panel's genes, the selected rules (each
with its cell type, thresholds, precision and recall) and the full pool of candidate rules.
\texttt{Panel.reselect} selects again from that pool at another $K$ or $O_{\max}$ without relearning;
this study used it for the RGC panels with $K=1$--3. \texttt{classify} labels a target's cells with a
panel, and \texttt{metrics.evaluate} scores the labels. A dataset configuration, one JavaScript Object
Notation (JSON) file, names the
reference's cell-type column, the cell types to leave out, and the renames and exclusions that map the
target's labels onto the reference's. On the command line, \texttt{xspear design}, \texttt{xspear
classify} and \texttt{xspear evaluate} wrap these calls. \texttt{xspear prior} lists and exports the
Xenium priors shipped with the package, \texttt{xspear profile} depth-normalises priors and
\texttt{xspear kcurve} plots accuracy against $K$ (Supplementary~S6). \texttt{xspear init} writes
annotated configuration files for the pipeline, and \texttt{xspear run} runs it, in full or for chosen
stages.

\paragraph{Pipeline.} The stages of the pipeline (Fig.~\ref{fig:pipeline}) pass files to each other.
Every selector writes the panel's gene list, which is all that stages 08--11 read, and \dart{} also
writes its rules. Each stage records its outcome, with the package version and commit, in a run
manifest, and a stage does not start if an earlier one failed. Stage 00 loads counts (an AnnData h5ad
file or a Matrix Market (MTX) matrix) and can attach Ensembl gene identifiers from a gene transfer
format (GTF) file or mygene.info (needed for the export bundle), merge per-cell metadata and assign a
group label such as the sample of origin. Stage 01 computes quality metrics and keeps
cells with 500--8{,}000 detected genes and genes detected in at least three cells; it can also filter on
mitochondrial fraction and call doublets with Scrublet~\citep{wolock2019scrublet}. Stage 02 caps the cells
per cell type (3{,}000, seeded). Stage 03 normalises counts per 10{,}000, or per median library size,
applies $\log(1+x)$ and can flag highly variable genes. Stage 04, off by default, computes principal
components, a neighbour graph, a uniform manifold approximation and projection
(UMAP)~\citep{mcinnes2018umap} and Harmony
integration~\citep{korsunsky2019harmony}. Stage 05 loads the target Xenium and its labels, which are used
only for scoring. Stage 06 resolves a Xenium prior and thins a copy of the reference to it
(Supplementary~S6); every data-driven selector of stage 07 uses the thinned copy, and the classifier of
stage 08 uses the original. Stage 07 runs \dart{} as four substages, 07a rule learning, 07b rule scoring,
07c ranking and 07d orthogonal selection (Sections~\ref{sec:methods-boolean}
and~\ref{sec:methods-rule-selection}; defaults $K=4$, $O_{\max}=0$ and 1{,}000 estimators per ensemble, as in
this study).
It can instead run the Wilcoxon test, COSG or NS-Forest (10 genes per cell type by default), read a
curated marker list, run \spapros{} or its reference probe sets (principal components, differential
expression, highly variable genes and random genes), or adopt an existing panel. Stage 10 writes the panel
and, for the Xenium Panel Designer, a reference bundle of counts, gene and cell tables and cell types.
Stage 11, off by default, is the off-target check (Section~\ref{sec:methods-off-target}). This study ran
stage 00 to load the references of the three Xenium targets and called the functions behind stages 06, 07
and 11 from its analysis scripts; the capping, splitting and scoring described below are in the paper
repository.

\paragraph{Classification and evaluation.} Stage 08 trains a classifier on the reference restricted to
the panel and labels the target's cells. The default is a one-versus-rest logistic regression ($C=1$, at
most 1{,}000 iterations, unweighted), as in this study's scoring; a random forest (500 trees, depth 8)
and balanced class weights are options. Reference and target are first normalised over the genes they
share, as in this study's scoring, and so does the library's \texttt{classify}; normalising over the
panel's genes alone, which is what a Xenium run measuring only the panel provides, is an option. Cells can also be labelled by voting with \dart{}'s rules instead of a classifier
(Supplementary~S5). Stage 09 reports \Fone{} and weighted $F_1$ over the reference's cell types,
per-type precision, recall and $F_1$, the confusion matrix and the fraction of cells assigned a type. An
alternative evaluation built on \spapros{}' forest classifier is described in Supplementary~S7.

\paragraph{Environments.} The analyses ran in Python 3.11.15 with numpy 2.4.6, pandas 2.3.3, SciPy
1.17.1, scikit-learn 1.9.0~\citep{pedregosa2011sklearn}, scanpy 1.10.3~\citep{wolf2018scanpy}, anndata
0.12.19, imodels 3.0.0~\citep{singh2021imodels}, Cython 3.3.0 (for the compiled \atpf{} criterion), COSG
1.2.0 and NS-Forest 4.1. \spapros{} 0.1.6 cannot run on numpy 2 or pandas 3, so \xspear{} runs it in a
second environment (numpy 1.26.4, pandas 2.3.3) as a subprocess that returns only the panel file. The
package's installer builds both environments, pinned to lock files of exactly these versions.

\subsection{Datasets and preprocessing}
\label{sec:methods-data}

Table~\ref{tab:datasets} lists every dataset. For each target the candidate gene pool is the set of
genes measured in both the single-cell reference and the Xenium panel: 4{,}766 (ovarian cancer),
4{,}548 (cervical cancer) and 4{,}791 (mouse brain). Expression is $\log(1+x)$ of counts per 10{,}000
(CP10k), computed over this shared universe for both modalities before subsetting to a panel. For the
benchmark, the reference was capped at 40{,}000 cells by stratified sampling, keeping small cell types
whole, and the selectors were run on a stratified 80\% of the capped reference.

The target Xenium cells were divided into design and evaluation halves by a spatial checkerboard, and
only the evaluation half was scored: about 183{,}000 cells for ovarian cancer, 181{,}000 for cervical
cancer and 32{,}000 for mouse brain. The design half was not used.

\paragraph{Evaluation labels.} For ovarian and cervical cancer, the labels are 10x Genomics' cell-type
annotation, mapped to the reference vocabulary. For cervical cancer, 10x left 56\% of cells
unassigned, and those cells were excluded. For mouse brain, the labels come from our label transfer,
a scaled multinomial logistic regression trained on the reference over the shared genes. The original
transfer normalised the reference over its full transcriptome but the Xenium over its panel; rebuilding
it with consistent normalisation changed the labels of 18.3\% of evaluation cells, and mouse brain
results use the rebuilt labels except where noted (Supplementary~S3). Labels are used only to score.

\subsection{Rule generation}
\label{sec:methods-boolean}

\paragraph{\atpf{}.} \atpf{} grows scikit-learn~\citep{pedregosa2011sklearn} decision trees with a
custom precision--recall impurity, $I(n)=1-P\times R$, where $P$ is precision and $R$ recall. This
criterion targets isolation of one cell type rather than global class separation. Root-to-node paths
are extracted as candidate rules.

\paragraph{\rulefit{}.} \rulefit{}~\citep{friedman2008rulefit} (imodels 3.0.0) is applied to
random-forest and gradient-boosted ensembles. Rules with nonzero coefficients in the
$L_1$-penalised logistic model are kept.

Each \atpf{} tree is grown on a bootstrap sample of the thinned reference, to depth 4, considering the
square root of the number of genes at each split. Every root-to-node path, not only those that end in a
leaf, is a candidate rule, and rules with the same conditions are merged. The \atpf{} criterion is a
compiled extension of scikit-learn, built for the installed Python and scikit-learn versions.
\rulefit{} uses imodels' \texttt{RuleFitClassifier} on a random forest with balanced class weights or on a
gradient-boosted ensemble with 80\% subsampling, both of depth 4, with up to 4{,}000 rules at 1{,}000
trees. The benchmarks used 1{,}000 estimators per ensemble. All candidate rules are evaluated on the full
thinned design slice to give $P$, $R$ and $F_1$ (and $F_\beta$ for $\beta=0.5$ and 2, which are not
used). A cell type with fewer than five reference cells is skipped, and a cell type that receives no rule
is reported as an error, after which the pipeline stops. Cell types are learned independently and can be
spread over worker processes; each learner is seeded per cell type, so the candidate pool does not depend
on the number of workers.

Three scoring options, not used in this study, change the candidate pool (stage 07b): discarding rules
with a ``$\le$'' condition; recording each rule's $F_1$ against each other cell type alone, computed on
the cells of the two types, whose minimum can replace one-versus-rest $F_1$ in ranking; and rewriting each
threshold as the fraction of reference cells at or below it, so that a rule fires at the same rank of any
data it is applied to (Supplementary~S5).

\subsection{Rule ranking and orthogonal selection}
\label{sec:methods-rule-selection}

\atpf{} candidates pass a quality gate: $P\ge0.75$ and $R\ge0.50$; if none pass, $P\ge0.45$ and
$R\ge0.30$; if still none, no gate. The 20 best by $F_1$ are then kept for each rule length. The top 40
\rulefit{} rules per cell type by penalised $F_1$ are kept. Pooled candidates are ranked by a penalised
score,
\begin{equation}
  S = F_1 - \lambda\cdot\max(0,\,N_{\text{genes}}-1),
  \label{eq:penalized-fbeta}
\end{equation}
in two tiers. Rules with $F_1\ge0.7$ are ranked first, with $\lambda=0.10$, and the rest follow with
$\lambda=0.04$. Ties are broken by $|P-R|$ and then by rule length. For each cell type, rules are added
greedily in rank order until $K$ have been kept. A candidate is skipped if it shares more than
$O_{\max}$ genes with the rules already chosen for that type, or if all its genes are already among
them. Genes may be shared across cell types. The panel is the union of the selected rules' genes.
Defaults are $K=4$ and $O_{\max}=0$; the selection constants sit on a plateau (Supplementary~S1).

The tier threshold, both penalties and the order of tie-breaking can be changed, and the candidate pool
can be kept before the quality gate, so that the gate, the number of rules kept per rule length and the
\rulefit{} top-$N$ can be varied without relearning. Supplementary~S1 used these, together with an option
that keeps every gene to one cell type, with the types choosing rules in turn. Two further selection
modes, not used here, are available from the library: a candidate's score can be reduced by $\lambda$
times the largest absolute Pearson correlation between its genes and those already on the panel, and a
total gene budget can replace $K$, spent first on each type's best rule that fits and then repeatedly on
the rule that most raises its type's best $F_1$ per gene it adds. Selection can also be restricted to the
genes the target Xenium measures, or to a list such as the genes that can be ordered on a panel.

\subsection{Detection-aware thinning with a Xenium prior}
\label{sec:methods-dart-transfer}

For each gene $g$, a Xenium prior gives
\begin{equation}
  \pkeep_{,g} = \min\!\left(1,\; d^{\mathrm{xen}}_g / d^{\mathrm{sc}}_g\right),
  \label{eq:pkeep}
\end{equation}
where $d^{\mathrm{xen}}_g$ and $d^{\mathrm{sc}}_g$ are the fractions of all cells with a nonzero count
in the prior's Xenium run and single-cell reference. Before learning, each nonzero entry of the
reference count matrix is kept intact with probability $\pkeep_{,g}$ and set to zero otherwise. This
per-entry thinning reproduces the prior's per-gene detection rate; binomial thinning of individual
molecules would not, because it barely changes detection for well-expressed genes. The thinned counts
are then normalised as above. The prior is used at its own magnitude and its labels are not used.
Genes are matched between prior and reference by gene symbol, and a gene the prior does not cover is not
thinned. This study's ovarian cancer prior ships with \xspear{} (Supplementary~S6).

\subsection{Xenium prior comparison}
\label{sec:methods-priors}

For each target, \dart{} panels were designed with each Xenium prior, with the target's own detection
profile (an upper bound) and with no prior, exactly as in the benchmark: the same capped reference and
design slice, 1{,}000 estimators per ensemble, $K=1$--10 and $O_{\max}=0$, scored with the benchmark
classifier on the held-out target Xenium. The panels designed with each target's benchmark prior are
therefore the benchmark's \dart{} panels. Here gene symbols were matched ignoring case, so that a prior
from the other species reaches the genes whose human and mouse symbols agree (for example human
\emph{GFAP} and mouse \emph{Gfap}): 2{,}139--2{,}348 of the target's 4{,}548--4{,}791 genes (45--49\%).
\xspear{} itself matches symbols exactly, which is the same for priors of the target's species. Because panel size at a given $K$ differs between priors,
\Fone{} was interpolated along each prior's curve at the realised sizes of the target's benchmark
panels with $K=3$--6 (45--83 genes for ovarian cancer, 41--79 for mouse brain and 17--32 for cervical
cancer) and averaged.

\subsection{Benchmark protocol}
\label{sec:methods-benchmarking}

\paragraph{Methods compared.} We used the Wilcoxon rank-sum test (scanpy
1.10.3~\citep{wolf2018scanpy}; top-$k$ genes per cell type), COSG 1.2.0 and NS-Forest 4.1, at
$k\in\{1,2,3,4,6,8,10,12\}$ genes per cell type for the Wilcoxon test and COSG and $k\in\{2,4,8\}$ for
NS-Forest. \spapros{} 0.1.6 ran through \xspear{}'s \spapros{} interface, in its own environment
(Section~\ref{sec:methods-software}), at its default size; as \xspear{} prepares it, it saw
$\log(1+x)$ CP10k expression with 4{,}000 highly variable genes flagged. Every method selected from the
same cells, the stratified 80\% slice of the capped reference (Section~\ref{sec:methods-data}), and
from the single-cell reference as measured. \dart{}
used the Xenium prior described in Section~\ref{sec:results-benchmark-design}.

\paragraph{Scoring.} Every panel in this paper, on Xenium and out of fold, was scored by one
procedure. The training cells and the test cells are normalised over the genes they share and then
restricted to the panel's genes; a one-versus-rest logistic regression ($C=1$, unweighted) is trained
on the training cells and predicts every test cell. For the Xenium targets the training cells are the
full single-cell reference without thinning and the test cells the held-out target Xenium half. \Fone{} averages per-cell-type $F_1$
over the reference's cell types; weighted $F_1$ weights each type by its number of held-out cells.
Comparisons at matched size interpolate each method's \Fone{} linearly in realised panel size and
average the difference over \dart{}'s sizes that lie within the other method's range. NS-Forest and
\spapros{} give no curve, so \dart{}'s curve was interpolated at their panel sizes; NS-Forest's margin
is the mean over its three budgets.

\paragraph{Seed variability.} The reference thinning and every learner were repeated at four seeds (1,
2, 3 and 42; the benchmark uses 42), and each seed's \dart{} curve ($K=1$--6 and 8) was compared with
the other methods' panels exactly as in the benchmark. Across seeds, the standard deviation of
\dart{}'s margin over a method was 0.005--0.017 \Fone{} (0.006--0.008 over the Wilcoxon test), and that
of \dart{}'s \Fone{} at a single $K$ was up to 0.024. A \dart{} panel's genes partly change between
seeds (mean pairwise Jaccard index 0.62--0.74 at $K=4$).

\paragraph{Compute.} Compute was measured on this study's own runs, one job per target and method
on the same cluster. For \dart{}, the Wilcoxon test, COSG and NS-Forest, the job loaded the
benchmark's design slice and timed the method's selection call exactly as the benchmark makes it
(\dart{} with the target's Xenium prior, $K=4$, $O_{\max}=0$ and 1{,}000 estimators; \dart{} and NS-Forest
with eight worker processes), recording the wall time of the call, the CPU time of the process and its
workers, and the peak memory of the largest process. For \spapros{}, the scheduler summary of its job
gave the same quantities, which also cover loading and preparing the data.

\subsection{Single-cell out-of-fold evaluation on RGC subtypes}
\label{sec:methods-oof}

The RGC atlas (34{,}029 cells, 45 subtypes) was restricted to the 4{,}911 genes of the mouse Xenium
5K panel and split into five folds, stratified by subtype (scikit-learn \texttt{StratifiedKFold}, shuffled,
seed 42). For each fold, every method selected its panel from the other four folds only, so the
held-out cells never influenced their own panel. Each panel was scored as on Xenium (Benchmark
protocol), with the four training folds as the training cells and the held-out fold as the test cells.
The predictions of the five folds were pooled, and \Fone{} and per-subtype $F_1$ were computed over all
34{,}029 cells.
No Xenium prior was used, because there is no Xenium measurement to anticipate. \dart{} was run with
$K=4$ rules per subtype and $O_{\max}=0$ at 1{,}000 estimators, and smaller panels were obtained as
the first $K=1$--3 rules of each fold's $K=4$ selection; the Wilcoxon test and COSG were run at 4, 6, 8,
10 and 12 genes per subtype and NS-Forest at its default, and \spapros{} was run on each fold's
training cells exactly as for the Xenium targets, at its default size (170--182 genes; a request for
200 genes returned the same panel) and at 50, 100 and 150 genes. Per-subtype differences were compared at near-matched panel sizes (Fig.~\ref{fig:rgc}b),
and subtypes were split into thirds by their number of cells. For the knockout analysis, each gene
of a fold's $K=4$ panel was removed in turn, the classifier was retrained without it, and the drop in
\Fone{} on the held-out fold was recorded.

\subsection{Probe off-target check}
\label{sec:methods-off-target}

After probe design, the Xenium Panel Designer returns a probe report listing, for each probe, its
predicted off-target genes. \xspear{} separates probes with and without predicted off-targets,
attaches the cell type each marker was selected for, and, for each marker with off-targets, draws a
dotplot of the marker and its off-target genes across the cell types of the annotated reference
(scanpy; expression as CP10k then $\log(1+x)$). The user replaces probes whose off-target genes are
expressed in cell types the marker does not mark. Fig.~\ref{fig:offtarget} was drawn this way by
\xspear{}'s off-target stage, with the human retina atlas's major cell classes (up to 3{,}000 cells
each) as the reference.

% ==============================================================================
\section*{Statements}

\subsection*{Statement of ethics}
This study analysed only publicly available, de-identified human datasets and published mouse
datasets (see the Data availability statement). No new human or animal data were generated for the
results reported here, so institutional review was not required.

\subsection*{Conflict of interest statement}
The authors have no conflicts of interest to declare.

\subsection*{Author contributions}
RP developed \dart{}, built most of \xspear{}, performed the benchmarking and analyses, and drafted the
manuscript. RP and BD packaged \xspear{} as a Python package. VP prepared the RGC single-cell reference,
verified it against the public GEO data, and identified the normalisation default that \xspear{}'s rule
voting now corrects. YC conceived and supervised the study. All authors reviewed and approved the
manuscript.

\subsection*{Data availability statement}
All datasets are public:
\begin{itemize}
  \item ovarian cancer: 10x Genomics
    \url{https://www.10xgenomics.com/datasets/17k-human-ovarian-cancer-scFFPE} and
    \url{https://www.10xgenomics.com/datasets/xenium-prime-ffpe-human-ovarian-cancer};
  \item cervical cancer: 10x Genomics
    \url{https://www.10xgenomics.com/datasets/17k-human-cervical-cancer-scFFPE} and
    \url{https://www.10xgenomics.com/datasets/xenium-prime-ffpe-human-cervical-cancer};
  \item mouse brain: 10x Genomics
    \url{https://www.10xgenomics.com/datasets/xenium-prime-fresh-frozen-mouse-brain}, with the
    single-nucleus reference from CZ CELLxGENE
    \url{https://cellxgene.cziscience.com/collections/31937775-0602-4e52-a799-b6acdd2bac2e};
  \item mouse retina: Gene Expression Omnibus (GEO) GSE325478 (reference; sample GSM9604550) and
    GSE307031 (Xenium; sample GSM9214539);
  \item breast cancer: GEO GSM7782698 and Zenodo 10076046 (reference) and 10x Genomics
    \url{https://www.10xgenomics.com/datasets/xenium-prime-ffpe-human-breast-cancer} (Xenium);
  \item RGC atlas: GEO GSE133382~\citep{tran2019rgc};
  \item STHELAR annotations (Supplementary~S3): BioStudies S-BIAD2146;
  \item human retina atlas (off-target example): CZ CELLxGENE
    \url{https://cellxgene.cziscience.com/collections/4c6eaf5c-6d57-4c76-b1e9-60df8c655f1e}.
\end{itemize}
The one input that is not public is the probe report of the human retina panel in
Fig.~\ref{fig:offtarget}, which belongs to an unpublished study by the authors' laboratory.

\subsection*{Code availability statement}
\xspear{} is an MIT-licensed Python package at \url{https://github.com/yuyanchenglab/xspear}; the
version described here is commit \texttt{bb86b4a}. The Methods and Supplementary~S5--S7 describe
its interfaces, pipeline stages and options. The analysis code, result tables and figure scripts are at
\url{https://github.com/yuyanchenglab/xspear-paper}.

\subsection*{Acknowledgements}
Support for this work was provided by the unrestricted Research to Prevent Blindness (RPB)
departmental fund to the Scheie Eye Institute. The funder had no role in the design, data collection,
data analysis, and reporting of this study.

Computation resources were generously provided by Penn Medicine Academic Computing Services (PMACS),
whose High-Performance Computing (HPC) and Archive systems were funded by National Institutes of
Health (NIH) grant 1S10OD012312.

% ==============================================================================
\bibliographystyle{unsrtnat}
\bibliography{references}

% ==============================================================================
\clearpage
\setcounter{figure}{0}
\setcounter{table}{0}
\renewcommand{\thefigure}{S\arabic{figure}}
\renewcommand{\thetable}{S\arabic{table}}
\renewcommand{\theHfigure}{supp.\arabic{figure}}
\renewcommand{\theHtable}{supp.\arabic{table}}
\setcounter{page}{1}
\renewcommand{\thepage}{S\arabic{page}}
\begin{center}
  {\Large\bfseries Supplementary Information}\\[0.5em]
  {\large \xspear{}: An Extensible Framework for Spatial Panel Evaluation and Refinement}\\[0.4em]
  \authornames
\end{center}
\noindent References cited here are listed in the main manuscript. Figure, table and section numbers
without an ``S'' refer to the main manuscript.
\label{sec:supplementary}

\subsection*{S1. \dart{}'s selection constants are on a plateau}
We swept every inherited selection constant one at a time (29 configurations), each with and without
cross-type gene sharing, over $K=1$--8:
\begin{itemize}
  \item the \atpf{} quality gate;
  \item the number of rules kept per rule length;
  \item the \rulefit{} top-$N$;
  \item the tier-1 $F_1$ threshold;
  \item both length penalties;
  \item the tie-break rule;
  \item $O_{\max}\in\{0,1\}$.
\end{itemize}
Configurations were judged at fixed realised panel sizes, with each target's Xenium prior. Only the
length penalty and $O_{\max}$ were load-bearing. Removing the length penalty roughly doubled the share
of multi-gene rules, added about 55 genes and cost 0.045 on ovarian cancer (0.014 on mouse brain) even
at matched size; Fig.~\ref{fig:ko} shows the effect of $O_{\max}$. Every other variation was within
$\pm0.01$ of the default on ovarian cancer and mouse brain. A design-time proxy, a held-out slice of
the reference thinned by the Xenium prior, could not resolve differences inside this plateau (its
ranking of configurations correlated with the target Xenium at Spearman $-0.23$ on ovarian cancer
and $+0.12$ on mouse brain). Choosing the constants on the \emph{other} datasets' Xenium, which the
deployment rule permits, changed accuracy by only $+0.004$ \Fone{}. We therefore ship fixed defaults.
The mouse brain numbers in this section use the original labels.

\subsection*{S2. Detection profiles: sampling noise, composition and normalisation}
$\pkeep$ is estimated from a finite number of cells, so it carries sampling noise. To measure it, we
resampled with replacement the cells of the single-cell reference and of the Xenium cells the prior was
computed from (the design half of each run), recomputed $\pkeep$, and compared it with the estimate from
all cells, over the genes detected in at least 1\% of reference cells (200 resamples). One resample
differs from the full estimate by a root-mean-square of 0.018 for ovarian cancer (16{,}187 reference
cells, 3{,}497 genes) and 0.024 for cervical cancer (9{,}475 cells, 3{,}276 genes), against a standard
deviation of $\pkeep$ across genes of 0.29 and 0.26 (Fig.~\ref{fig:svar}). The noise is small next to
the differences between genes on which thinning acts.

\begin{figure}[!htbp]
  \centering
  \includegraphics[width=0.85\textwidth]{figures/fig_pkeep_noise.pdf}
  \caption{\textbf{Sampling noise of the detection-rate ratio $\pkeep$.} $\pkeep$ from one resample of
  the reference and Xenium cells against the estimate from all cells, for the genes detected in at least
  1\% of reference cells, for ovarian (A) and cervical cancer (B); dashed line, $y=x$. Each panel gives the
  root-mean-square error (RMSE) of one resample, averaged over 200 resamples, and the standard deviation
  (SD) of $\pkeep$ across genes.}
  \label{fig:svar}
\end{figure}

Because the profile is computed from all cells, it partly reflects cell-type composition.
Re-weighting within-type detection to the reference's composition moves $\pkeep$ by a median of 0.017
on ovarian and 0.019 on cervical cancer (Spearman 0.977 and 0.985 with the all-cell profile).

The reference's stored expression is normalised over its full transcriptome, while our benchmark
normalises over the shared universe. The Wilcoxon test is insensitive to the difference (mean \Fone{}
within $\pm0.003$). COSG is not: the full-transcriptome basis raises it by 0.015 on ovarian and 0.036
on cervical cancer, and lowers it by 0.003 on mouse brain (original labels). Comparisons with COSG
should be read with this in mind.

\subsection*{S3. Evaluation labels: provenance, validation and an independent annotation}
\paragraph{Provenance.}
\begin{itemize}
  \item \emph{Ovarian and cervical cancer.} The labels are 10x Genomics' annotation, mapped to the
    reference vocabulary. Cell types absent from the reference were removed, and for cervical cancer the
    56\% of cells 10x left unassigned were excluded.
  \item \emph{Mouse brain.} The labels are our own transfer from its reference. The original transfer
    normalised the reference over 20{,}409 genes and the Xenium over its 5{,}006-gene panel. Rebuilding
    it with both sides normalised over the same genes changed 18.3\% of evaluation-cell labels, all in
    one direction. Implausible immune and meningeal calls in a healthy coronal section fell sharply
    (T cells 770 to 36; vascular leptomeningeal cells 3{,}515 to 1{,}118), and the cells became
    neurons. Re-scoring all 138 mouse brain runs of our pipeline on the rebuilt labels kept their order
    (Spearman 0.981) and raised \Fone{} by a median of 0.062. Mouse brain numbers in this paper use the
    rebuilt labels except in S1 and S2, as noted.
\end{itemize}

\paragraph{Validation.} Fig.~\ref{fig:slabel} checks the \emph{true} labels of the ovarian and
cervical Xenium cells, that is 10x's annotation used for scoring, not any panel's predictions. For each
labelled Xenium cell type (evaluation half) we compared its mean expression profile with that of every
single-cell reference cell type over the shared genes, both normalised as for scoring. The matrices are
diagonal-dominant: the mean same-type Spearman correlation is 0.83 for ovarian and 0.82 for cervical
cancer, and each Xenium type's best match is its own reference type for all 6 cervical and 11 of 14
ovarian types. The three exceptions are tumour states (tumour, proliferative tumour and inflammatory
tumour cells) that match the closely related VEGFA-positive tumour cells as well as or slightly
better than their own type (by at most 0.016).

\begin{figure}[!htbp]
  \centering
  \includegraphics[width=0.5\textwidth]{figures/fig_label_validation_ovarian.pdf}\hfill
  \includegraphics[width=0.47\textwidth]{figures/fig_label_validation_cervical.pdf}
  \caption{\textbf{Validation of the true Xenium labels used for scoring.} Spearman correlation between
  the mean log-normalised profiles of single-cell reference cell types and of the labelled Xenium cell
  types (10x's annotation on the evaluation half, not panel predictions) over shared genes, for ovarian
  (left) and cervical cancer (right). Boxes mark each Xenium type against its own reference type. Mean
  diagonal $\rho$ is 0.83 and 0.82. The best match is the type's own reference type for 11 of 14
  ovarian and 6 of 6 cervical types.}
  \label{fig:slabel}
\end{figure}

\paragraph{Independent annotation.} The STHELAR resource provides an annotation of the same ovarian
Xenium section, at the level of five lineages, that was produced independently of 10x's (BioStudies
S-BIAD2146). We matched its cells to our evaluation cells by mutual nearest cell centroid (98.6\% of
cells matched). With the evaluation labels coarsened to the same five lineages, the two annotations
agree on 82.8\% of cells.

\subsection*{S4. Compute cost}
Table~\ref{tab:compute} lists the values plotted in Fig.~\ref{fig:sfig_compute}.

\begin{table}[!ht]
  \centering\small
  \caption{\textbf{Compute cost per panel-design method on each target.} Worker processes, CPU time
  (core-hours), peak memory of the largest process (GB) and wall time (hours), from this study's runs
  (Methods). \spapros{}'s values also cover loading and preparing the data.}
  \label{tab:compute}
  \resizebox{\textwidth}{!}{%
  \input{figures/compute_table.tex}}
\end{table}

\begin{figure}[!htbp]
  \centering
  \includegraphics[width=\textwidth]{figures/fig_compute.pdf}
  \caption{\textbf{Compute cost per panel-design method.} From this study's runs on the three targets
  (Methods). \textbf{(A)}~Total CPU time, including worker processes. \textbf{(B)}~Peak memory of the
  largest process. \textbf{(C)}~Wall time. \textbf{(D)}~\dart{}'s CPU time against its wall time with
  eight worker processes. (A), (C) and (D) use log scales. Exact values are in Table~\ref{tab:compute}.}
  \label{fig:sfig_compute}
\end{figure}

\FloatBarrier
\subsection*{S5. Labelling cells with \dart{}'s rules (rule voting)}
\label{sec:rule-voting}
Rule voting labels a target's cells with a \dart{} panel's own rules instead of a classifier: the
\texttt{rules} method of \texttt{classify}, \texttt{mode = "rules"} in stage 08, or
\texttt{xspear classify -{}-method rules}. This study did not use it: every prediction reported here comes from the classifier,
and we make no claim about rule voting's accuracy.

A rule fires in a cell when all its conditions hold on the cell's log-normalised expression. The
reference and the target are normalised over the genes they share, the scale on which the thresholds were
learned. Each cell is assigned the cell type with the most firing rules. With up to $K$ mostly
single-gene rules per type, ties are common, and a tie goes to the tied type with the highest probability
under the classifier trained on the reference; that is why a classifier is trained in this mode too
(without one, the first tied type is taken). A cell in which no rule fires is left unassigned. Besides
\Fone{} over the real cell types, with an unassigned cell counted as missed for its true type, the
evaluation then reports the fraction of cells assigned and \Fone{} among the assigned cells.

Two options change the thresholds before voting. Quantile thresholds (rule scoring, Methods) express each
threshold as a quantile of the reference, so that a rule fires at the same rank of the target's
distribution as of the reference's. Otsu calibration replaces each threshold with Otsu's
threshold~\citep{otsu1979threshold} for that gene in the target, which needs no labels.

\subsection*{S6. Xenium priors in \xspear{}}
\label{sec:package-priors}
\paragraph{The shipped ovarian cancer prior.} \xspear{} ships this study's ovarian cancer prior, measured on
10x Genomics' FFPE Flex reference against the Xenium Prime 5K run of the same tissue block: 4{,}766 genes,
median $\pkeep$ 0.590, with each gene's two detection rates. It is the prior used here, unchanged.
\texttt{xspear.xenium\_prior("ovarian")} returns it, \texttt{xspear prior ovarian -o FILE} writes a copy,
\texttt{-{}-p-keep ovarian} uses it wherever the command line takes a prior, and it is stage 06's default.
It suits a planned human FFPE Xenium Prime 5K experiment (Section~\ref{sec:results-priors}).

\paragraph{Other sources and gene matching.} A prior can also be a table of genes and $\pkeep$ (tables of
$1-\pkeep$ from earlier versions are converted on reading), or it can be measured from a paired
single-cell reference and Xenium run by Eq.~\ref{eq:pkeep}, over the genes they share. \xspear{} matches a
prior's genes to the reference's by exact symbol and leaves a gene the prior does not cover unthinned.
Stage 06 stops when a prior covers none of the reference's genes and warns when it covers fewer than half.
After thinning it recomputes stage 03's normalisation on the thinned copy.

\paragraph{Depth normalisation} (\texttt{xspear profile}; stage 06) is meant for a prior whose single-cell
reference was sequenced much more or less deeply than usual, which shifts its single-cell detection rates,
and therefore its $\pkeep$, as a whole. It keeps the prior's ranking of genes and replaces each value with
the value at the same percentile of an anchor profile, by default the shipped ovarian prior. Several
priors are combined through the mean, median or 20\% trimmed mean of their percentile ranks; a prior with
a median $\pkeep$ above 0.95 is left out as showing no detection loss; and a gene can be required to be
covered by a minimum number of priors. It is off by default: this study used every prior at its own
magnitude and did not evaluate depth normalisation.

\paragraph{Choosing $K$} (\texttt{xspear kcurve}). A stratified 25\% of the reference is held out and
thinned with the prior. Panels are designed on the remaining cells for $K=1,\dots,K_{\max}$ (one candidate
pool, reselected), a classifier trained on the unthinned remaining cells predicts the held-out cells, and
\Fone{} is reported against $K$ and panel size, to help choose $K$ within a gene budget. Thinned
single-cell data are cleaner than Xenium, so the values are optimistic. This study did not use it, and
Supplementary~S1 found that a held-out thinned slice of the reference could not rank selection constants
within their plateau.

\subsection*{S7. Other package functions}
\label{sec:package-other}
\paragraph{Off-target dominance factor.} Stage 11 can also score each marker that has predicted off-target
genes: the highest mean expression of any of its off-target genes in a cell type other than the one the
marker marks, divided by the marker's mean expression in the type it marks. A value of 1 or more is
flagged, and the dotplots are then ordered by the score. It is off by default and was not used for
Fig.~\ref{fig:offtarget}. Panel genes chosen for a reason other than marking a cell type can be labelled
through named gene sets, and the probe report can be split into specific and unspecific probes and
annotated without loading any expression data.

\paragraph{Forest-based evaluation.} Instead of scoring stage 08's labels, stage 09 can run \spapros{}'
forest-based classification on the panel's genes in the reference. It reports each gene's importance for
each cell type and whether that type is where the gene is most expressed, and can keep only the genes above
an importance threshold, with dotplots of the genes that do and do not match.

\paragraph{Supporting code.} A synthetic reference and target, with marker genes for each cell type on a
shared background and a target thinned by a known per-gene keep probability, test an installation end to
end in seconds. The \spapros{} worker records the versions it ran with, and its selection, which also yields
the reference probe sets, is cached by panel size. Two presets record whether a run used a Xenium prior and
map the case names of earlier runs onto them, and a function working in the units of earlier versions
($1-\pkeep$) reproduces their depth-normalised files exactly.

\end{document}
