## Supplementary figures and images for "X-SPEAR: An Extensible Framework for Spatial Panel Evaluation and Refinement"

### fig_compute.pdf

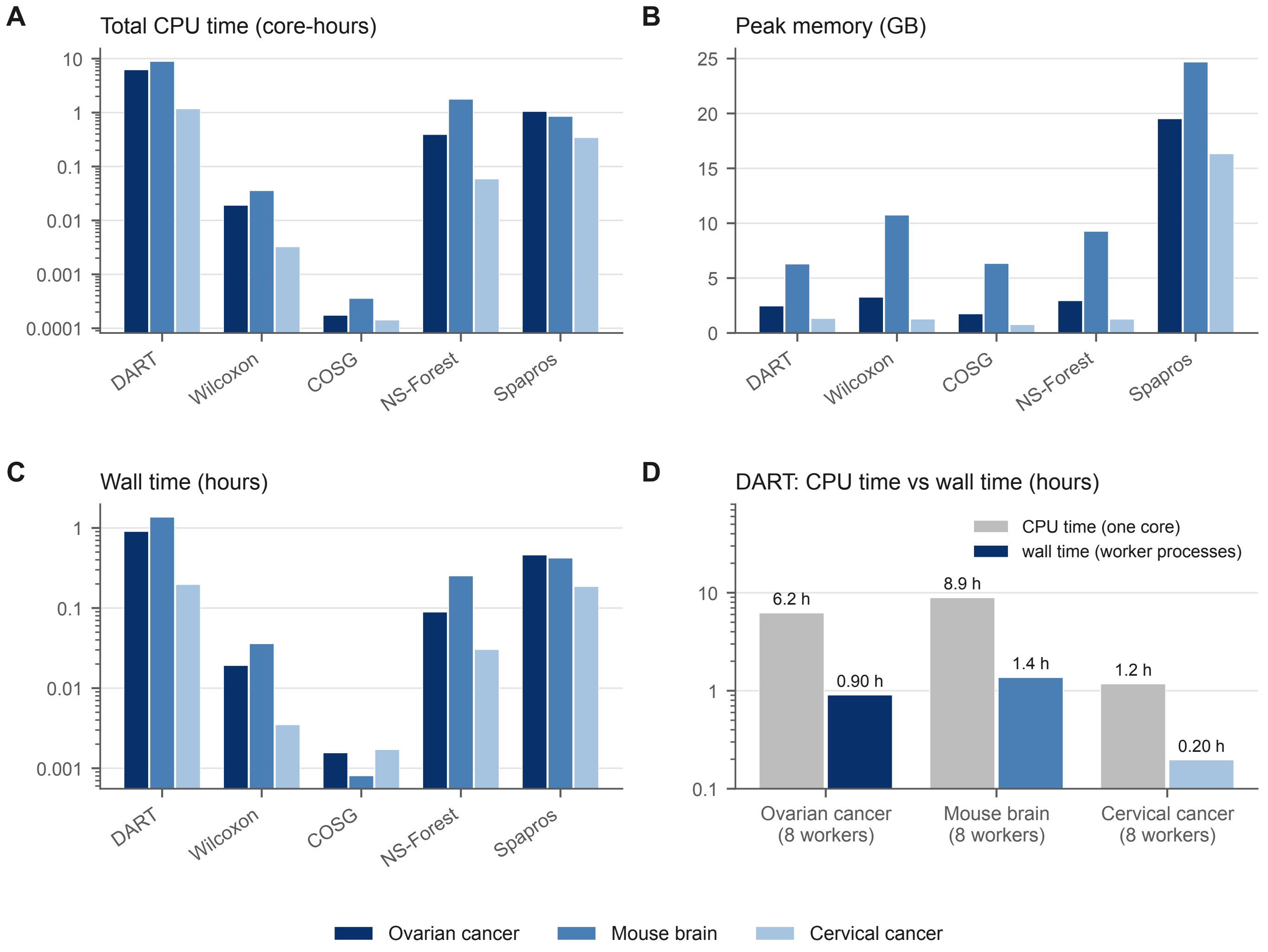

### fig_detection_reproducibility.pdf

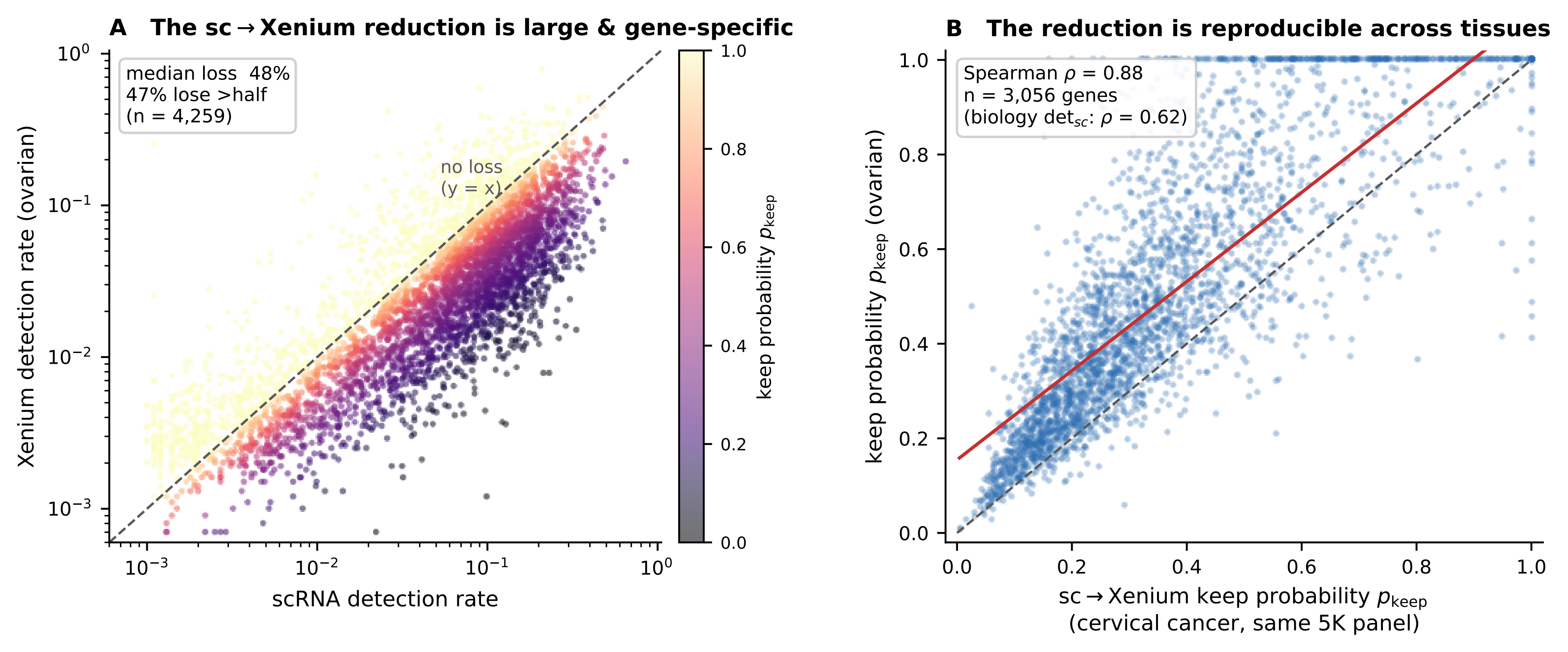

### fig_ko.pdf

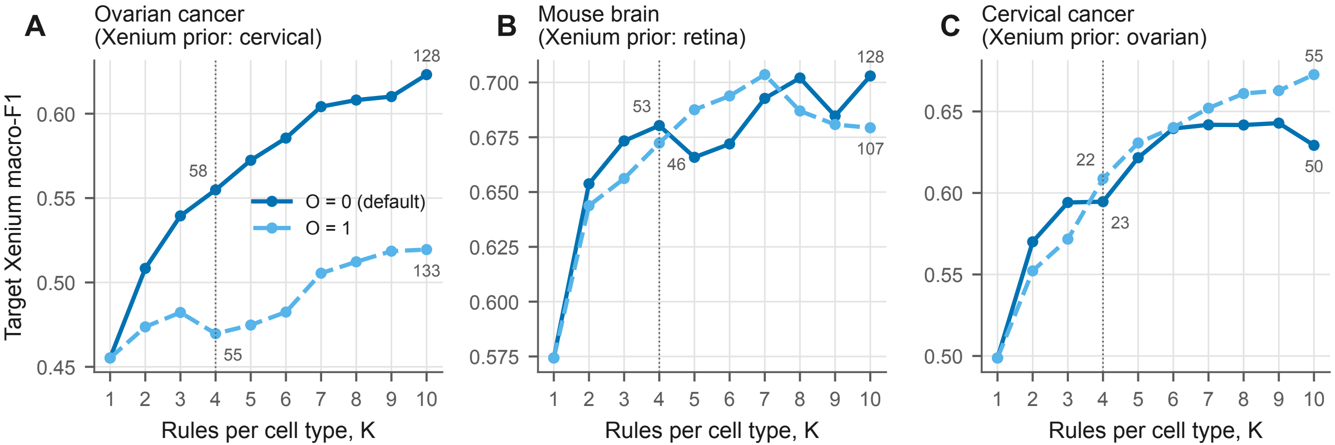

### fig_label_validation_cervical.pdf

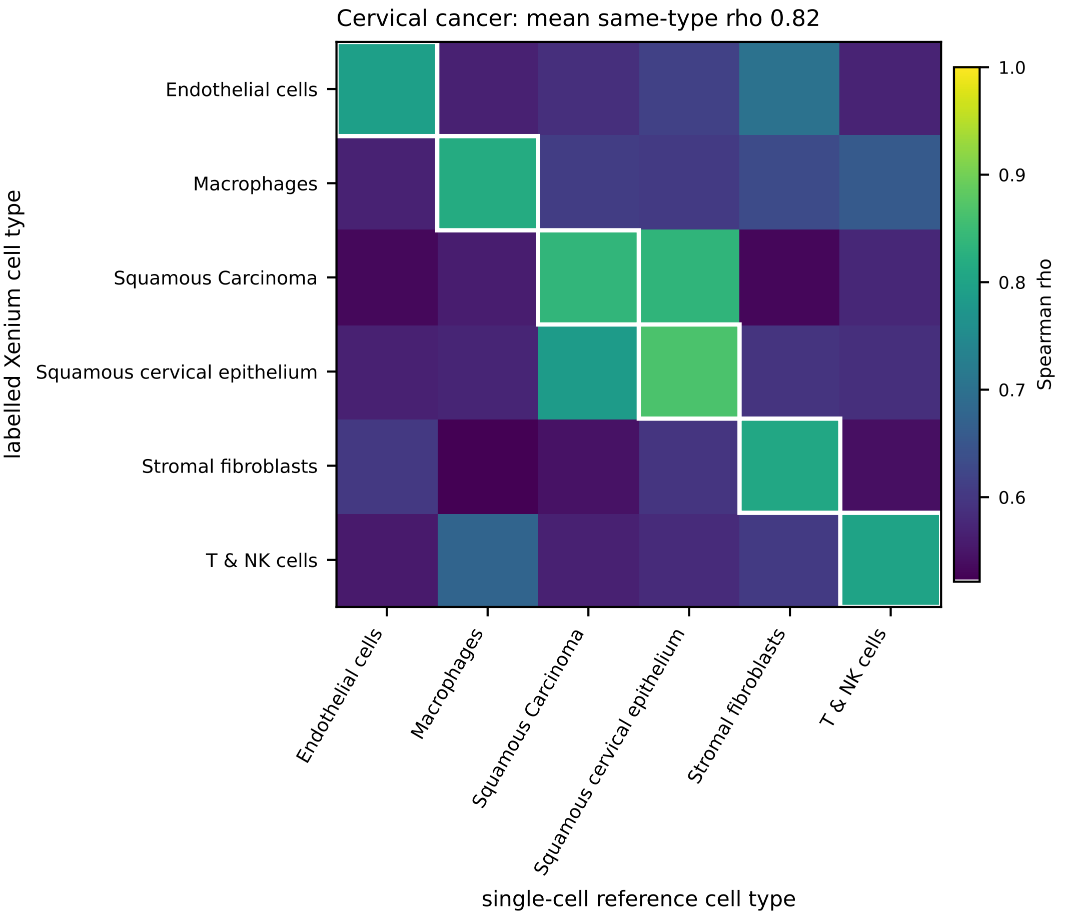

### fig_label_validation_ovarian.pdf

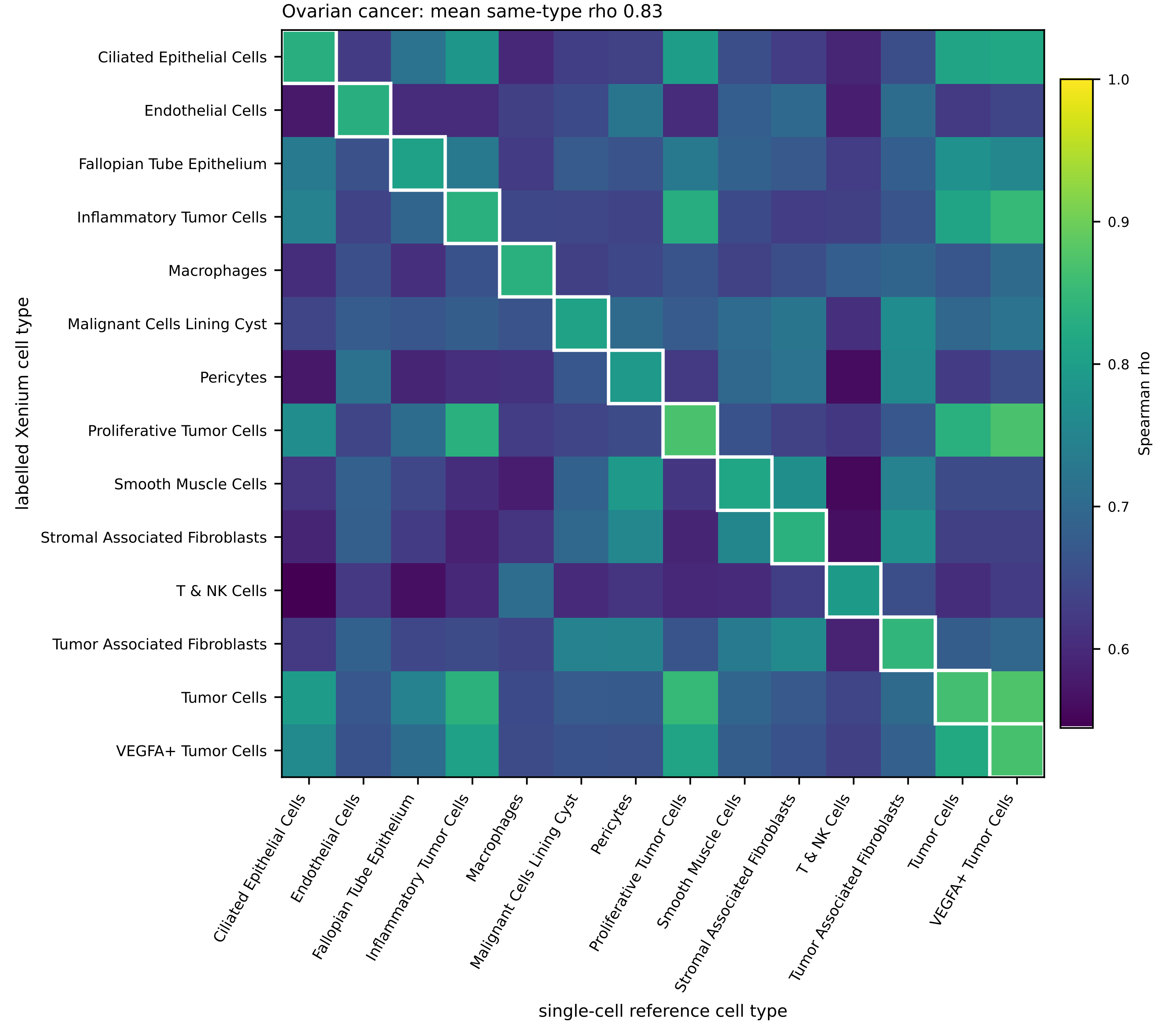

### fig_offtarget_CRX.png

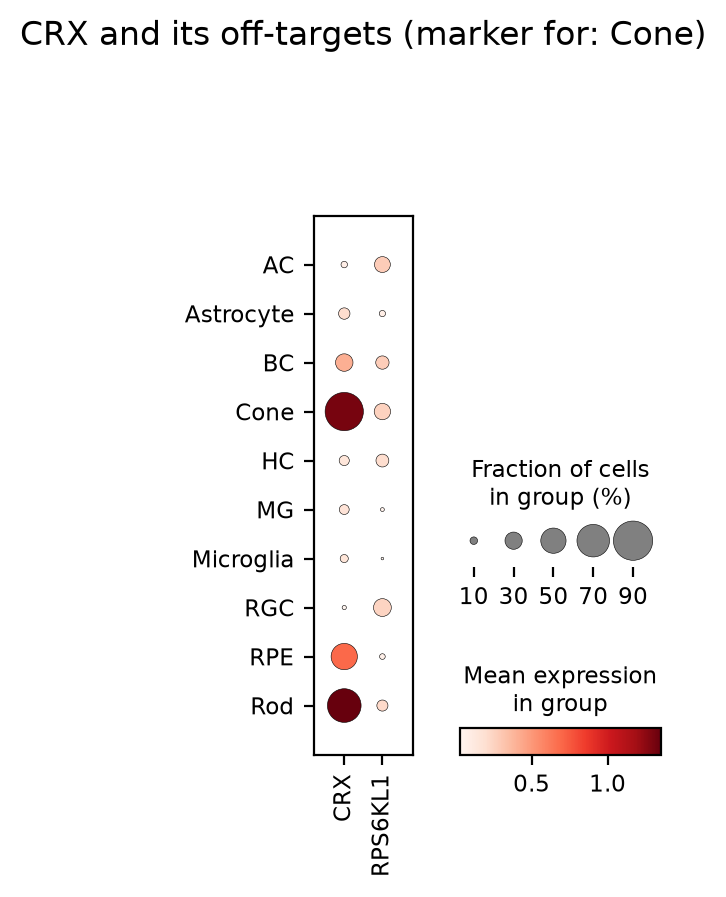

### fig_offtarget_PAX6.png

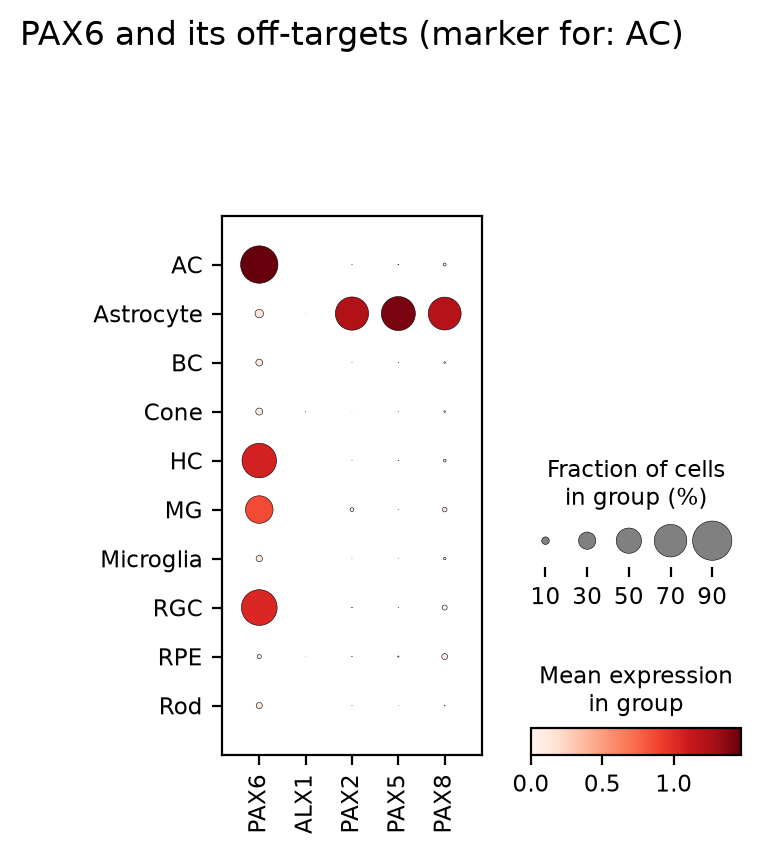

### fig_pkeep_noise.pdf

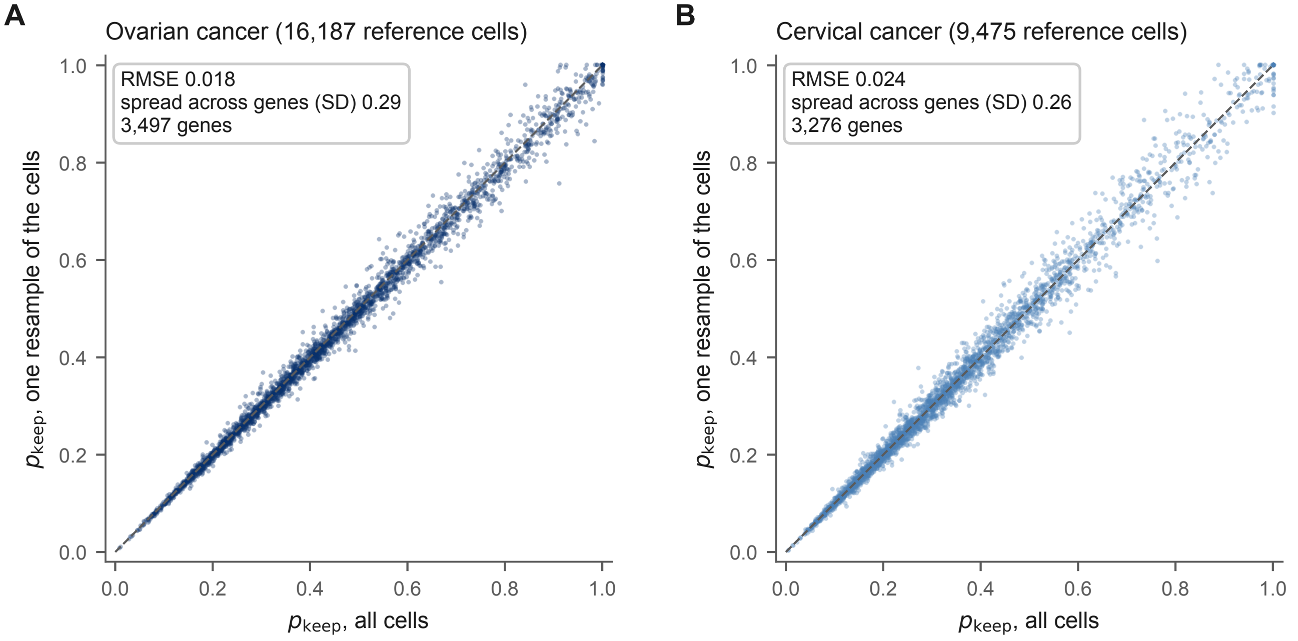

### fig_rgc.pdf

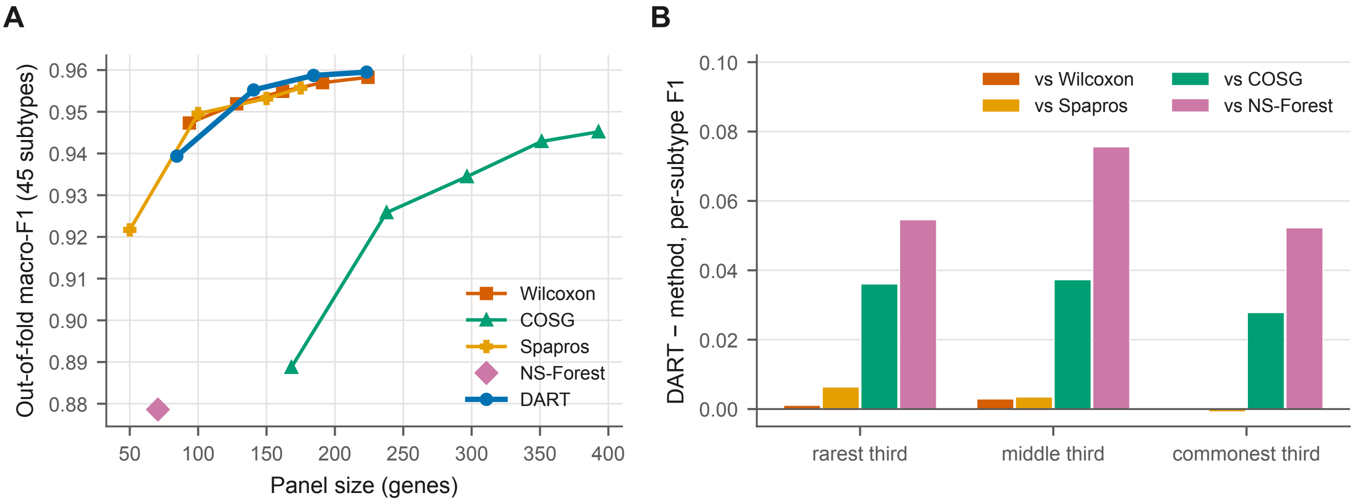
